# OASIS: An interpretable, finite-sample valid alternative to Pearson’s *X*^2^ for scientific discovery

**DOI:** 10.1101/2023.03.16.533008

**Authors:** Tavor Z. Baharav, David Tse, Julia Salzman

## Abstract

Contingency tables, data represented as counts matrices, are ubiquitous across quantitative research and data-science applications. Existing statistical tests are insufficient however, as none are simultaneously computationally efficient and statistically valid for a finite number of observations. In this work, motivated by a recent application in reference-free genomic inference (1), we develop OASIS (Optimized Adaptive Statistic for Inferring Structure), a family of statistical tests for contingency tables. OASIS constructs a test-statistic which is linear in the normalized data matrix, providing closed form p-value bounds through classical concentration inequalities. In the process, OASIS provides a decomposition of the table, lending interpretability to its rejection of the null. We derive the asymptotic distribution of the OASIS test statistic, showing that these finite-sample bounds correctly characterize the test statistic’s p-value up to a variance term. Experiments on genomic sequencing data highlight the power and interpretability of OASIS. The same method based on OASIS significance calls detects SARS-CoV-2 and Mycobacterium Tuberculosis strains de novo, which cannot be achieved with current approaches. We demonstrate in simulations that OASIS is robust to overdispersion, a common feature in genomic data like single cell RNA-sequencing, where under accepted noise models OASIS still provides good control of the false discovery rate, while Pearson’s *X*^2^ test consistently rejects the null. Additionally, we show on synthetic data that OASIS is more powerful than Pearson’s *X*^2^ test in certain regimes, including for some important two group alternatives, which we corroborate with approximate power calculations.

**Significance Statement:** Contingency tables are pervasive across quantitative research and data-science applications. Existing statistical tests fall short, however; none provide robust, computationally efficient inference and control Type I error. In this work, motivated by a recent advance in reference-free inference for genomics, we propose a family of tests on contingency tables called OASIS. OASIS utilizes a linear test-statistic, enabling the computation of closed form p-value bounds, as well as a standard asymptotic normality result. OASIS provides a partitioning of the table for rejected hypotheses, lending interpretability to its rejection of the null. In genomic applications, OASIS performs reference-free and metadata-free variant detection in SARS-CoV-2 and M. Tuberculosis, and demonstrates robust performance for single cell RNA-sequencing, all tasks without existing solutions.

---

**D**iscrete data on contingency tables is ubiquitous in classical and modern data science, and is central in the social sciences and quantitative research disciplines, including biology. In modern applications these tables are frequently large and sparse, leading to a continued interest in new statistical tests for contingency tables (2). One recent motivating application is SPLASH, a new approach to genomic inference which maps myriad problems in genomic sequence analysis to the analysis of contingency tables. These disparate applications include discriminating phylogenetically distinct strains or alternative splicing in single cell RNA-sequencing, among others (1).

There is a rich literature that addresses testing for row and column independence in contingency tables beginning with the work of Pearson, who designed the widely used *X*^2^ test in the early 1900s (3, 4). Other approaches include the likelihood ratio test, permutation or Markov Chain Monte Carlo (MCMC)-based methods (5), and modeling parametric deviations from the null with log-linear models (4).

Despite the prominence of Pearson’s *X*^2^ test, it suffers from multiple statistical drawbacks which limit its utility for scientific inference. First, the *X*^2^ test lacks robustness: it has high power against many scientifically uninteresting alternatives, for example against models where technical or biological noise cause a table to formally deviate from the specified null. We expand on this point in Section 6.A, and provide simulation evidence for noise stemming from biological overdispersion and contamination.

Second, the *X*^2^ test does not provide statistically valid p-values for any finite number of observations. There is substantial work on estimating significance thresholds, primarily centered on an asymptotic theory that assumes a fixed table size with the number of observations tending to infinity. For example, common guidelines (4) indicate that the *χ*^2^ distribution is a bad approximation when more than 20% of the entries take value less than 5. However, in modern tables of interest, this is often the case; the biological tables which motivated this test’s design have many (tens or hundreds) rows relative to the total number of observations per column (similarly in the tens or hundreds), violating the *X*^2^ use guidelines (1).

Other methods such as log-linear models suffer from similar limitations: namely, lack of robustness, and of calibration for finite observations (4). MCMC-based methods have also been developed, which provide statistically valid p-values, but despite significant recent works the lack of a closed form expression renders them too computationally intensive for large tables in practice (2, 5). Additionally, for permutation tests, *B* samples from the null can at best yield a p-value of 1/(B+1). Thus, many permutations are required to obtain sufficiently small p-values to reject the null, especially under the burden of multiple hypothesis correction. Finally, MCMC-based methods suffer from the robustness issues as *X*^2^. To our knowledge, no nontrivial p-values exist for contingency tables that are computable in closed form and are valid for a finite number of observations.

In this work, motivated by recent applications in genomic inference (1), we introduce OASIS, a powerful and general family of interpretable tests. Building on preliminary work (1), OASIS provides p-value bounds that 1) are valid for a finite number of observations, 2) have a closed form expression,3) are empirically robust to small deviations from the null distribution, and 4) empirically enable scientific inference that cannot be achieved with *X*^2^.

To build intuition for a task we seek OASIS to perform, consider the following data. 103 patients are infected with potentially different variants of SARS-CoV-2 (6). For each, a black box produces counts of the nucleotide composition of a segment of the SARS-CoV-2 variant’s genome, which can be considered a categorical variable. This problem formulation and motivation stems from contingency tables generated by SPLASH (1), and is discussed further in Section 6. Under the null hypothesis, each patient is infected with the same variant of the virus, and so all columns in the table will be drawn from the the same distribution over rows. If a new strain has infected a group of patients, then the row distribution for these patients will be different. The desired test should detect this deviation, and provide post-inferential guidance on why the table was rejected, ideally discriminating populations of patients on the basis of which strains infected them.

To understand why a new test is needed, we consider Pearson’s *X*^2^ in more detail. The *X*^2^ test statistic sums squared residuals after normalization (Figure 1B, Eq. (3)) The resulting sum is asymptotically *χ*^2^ distributed under the null, but can significantly deviate from this for a finite number of observations. When it rejects the null, Pearson’s *X*^2^ test provides no framework for interpreting why, leading practitioners to develop and use exploratory data-analysis tools such as correspondence analysis (7–10).

**Fig. 1.**
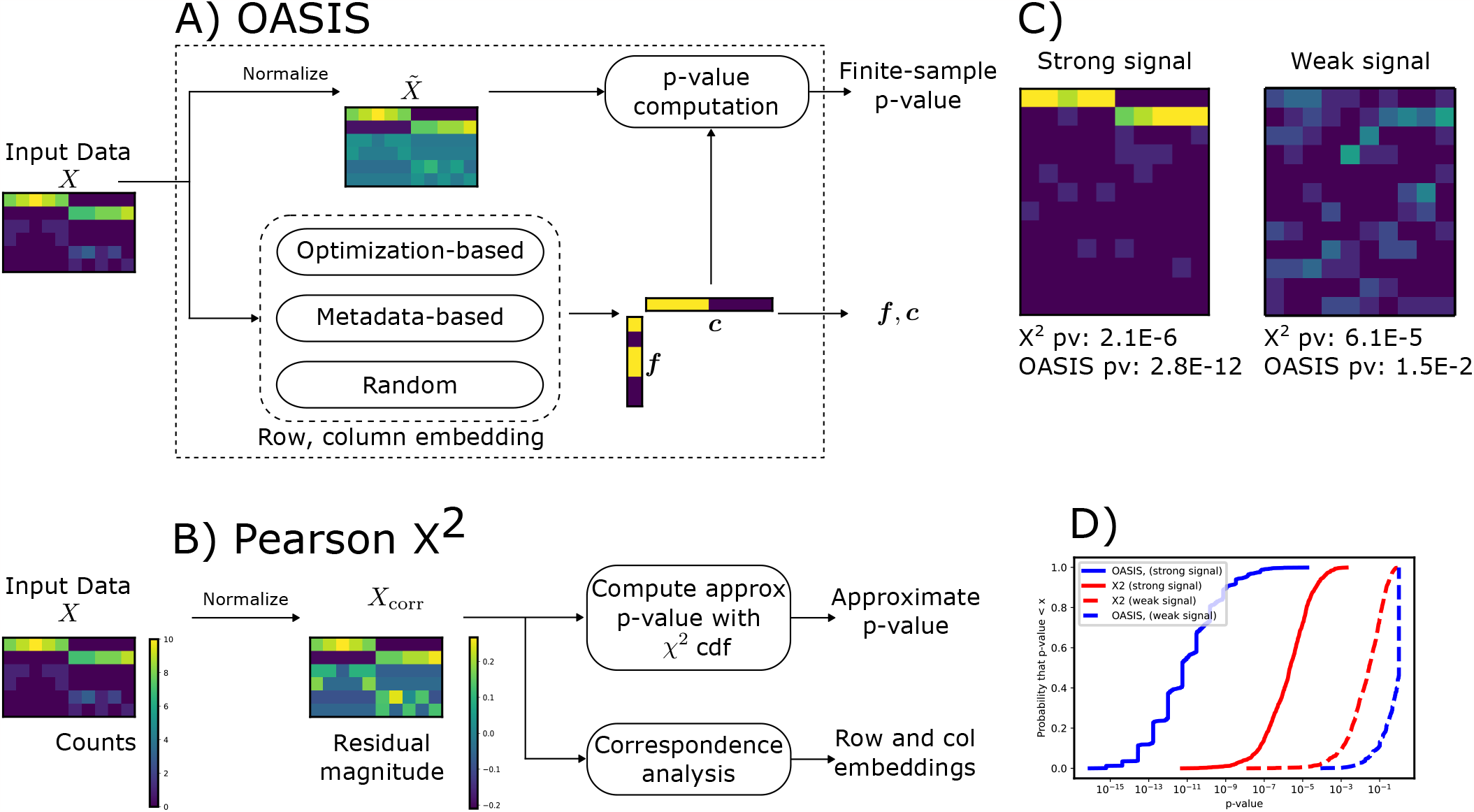
Comparison of OASIS and Pearson’s *X*^2^ test for input matrix *X ∈* ℕ^*I×J*^. **A)** OASIS computes a matrix of residuals 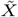 as in Eq. (1). Row (column) embedding vectors ***f***∈ ℝ^*I*^ (***c***∈ℝ^*J*^) are generated by one of several options (metadata can be incorporated if desired, e.g. an ordering of the rows, or similarity information regarding the columns). These vectors are used to compute the OASIS test statistic *S* in Eq. (2), which admits a finite-sample p-value bound using classical concentration inequalities. **B)** *X*^2^ computes a matrix of residuals *X*_corr_ as in Eq. (4), which is sensitive to deviations in low count rows, as seen in the bottom 4 rows in the example matrix *X* and *X*_corr_. The *X*_2_ test then provides an asymptotically valid p-value via a distributional approximation. For interpretability, practitioners often use correspondence analysis (4) to interpret rejection of the null, a procedure with no statistical guarantees, which can fail to detect the desired structure. **C)** depicts two example counts tables. The one on the left corresponds to concentrated (strong) signal, while the one on the right corresponding to diffuse (weak) signal. Both tables have 100 counts distributed evenly over 10 columns, with 12 rows. The generative model for these is detailed in Section 14.A, with additional plots provided. *X*^2^ assigns both of them similar significance, but OASIS assigns a much smaller p-value to the left table than the right, agreeing with our intuition. Figure 9 shows the spectra of the centered and normalized contingency tables, showing that OASIS prioritizes the first table with a spikier spectrum. **D)** plots the empirical CDF of the p-values of OASIS and Pearson’s *X*^2^. This is shown for the two classes of tables; ones with a strong concentrated two-group signal, and ones with a diffuse signal. OASIS yields significantly improved p-values for the case with strong signal, and substantially worse power than *X*^2^ in the weak signal case, which visually looks like noise. *X*^2^ on the other hand yields much more similar performance in the two settings. Here, `OASIS-opt` is shown, which is run over 5 independent splits of the dataset (additional details in Section 14.A).

OASIS seeks to improve these shortcomings by building in interpretability and analytic tractability in its construction of a linear test statistic (Figure 1A, Eq. (2)). This construction enables the use of classical concentration inequalities to yield p-value bounds that are valid for finite numbers of observations. As opposed to *X*^2^ which sums squared residuals, OASIS computes a bilinear form of residuals, similar in spirit but conceptually and statistically distinct from a Lancaster decomposition of *X*^2^ (11) and related polynomial decomposition methods (12). Fluctuations or single large residuals will be averaged, and are unlikely to generate a large test statistic. We make this observation precise later in our discussion of the linear algebraic characterizations of these approaches. The most similar method to OASIS regarding interpretable decompositions of contingency tables is correspondence analysis, (7–10), an exploratory method for post-facto interpretation that provides no statistical guarantees.

One recent work which shares some similarities with ours is (13), which provides a method for estimating graph dimension with cross-validated eigenvalues. Their method, like ours, is based on splitting the data into two portions, generating embeddings on one part, and testing the signal strength on the held-out portion. While general, this method requires additional assumptions on the embeddings generated and used for inference (no one entry is too large), and critically is only able to provide asymptotically valid p-values. The authors discuss an extension of their method to contingency table analysis, but inherently can only provide asymptotic p-values. In this work, with our more analytically tractable test statistic, we are able to provide a closed form p-value bound which is valid for any finite number of observations.

In the rest of this paper, we formalize OASIS, state several of its theoretical properties, contrast it with *X*^2^, and present several variants and extensions of the OASIS test. Here, we do not seek to expound on the full theoretical generality of OASIS, but instead provide an applied exposition and disciplined framework for computing some optimization-based instantiations of the statistic, and illustrate the applied performance of OASIS in simulations and in real biological data. Simulations show that OASIS is a robust test that controls the level in a variety of models where the null is formally violated, but without a biological or scientific meaning. Simulated alternatives show that OASIS has power in many settings of interest, and in fact has more power than *X*^2^ in a variety of settings including for two group alternatives. Biological examples show that, with no parameter tuning, OASIS enables scientific inference currently impossible with Pearson’s *X*^2^ test; for example, OASIS has 100% accuracy in distinguishing discrete patient populations infected with Omicron-BA.1 and BA.2 from those infected with the Delta variant without knowledge of a reference genome or any sample metadata. In a different biological domain, analyzing M. Tuberculosis sequencing data, OASIS precisely partitions samples from two sub-sublineages, again with no metadata or reference genome. Finally, the OASIS framework enables more general tests and analyses which we discuss in the conclusion. A glimpse at such examples include a disciplined alternative statistical framework for matrix decomposition beyond the SVD, and inference on multiple tables defined on the same set of columns.

## 1. Problem Formulation

As is standard in contingency table analysis, the observed matrix of counts is taken as *X* ∈ ℕ^*I×J*^. Defining [*m*] ={ 1, 2, …, *m* }, a contingency table is then defined by pairs of observations of a row ([*I*]-valued) categorical random variable, and a column ([*J*]-valued) categorical random variable. In this setting, we focus on the case where the columns correspond to biological samples (explanatory random variable) and the rows correspond to the response variable (4). Thus, we are interested in whether the conditional row distribution is the same for each column. Define *X*^(*j*)^ as the *j*-th column of *X*, and 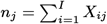 as the total number of counts in column *j*. Without loss of generality, we assume that *n*_*j*_ *>* 0∀*j*, as otherwise this column can be omitted. Let *M* =∑_*j*_ *n*_*j*_ be the total number of counts in the table, equivalently obtainable by summing row or column sums. The null model studied is:

### Definition 1

(Null Model). *Conditional on the column totals* 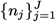, *each column of the contingency table X is X*^(*j*)^ ∼ multinomial(*n*_*j*_, ***p***), *drawn independently for all j, for some common vector of (unknown) row probabilities* ***p***.

Contingency tables are well studied, and we refer the reader to (4) for further background. A classical contingency table example studies the relationship between aspirin use and heart attacks, where the columns correspond to aspirin or placebo use, and the rows correspond to Fatal Attack, Nonfatal Attack, or No Attack. It is found that conditioning on whether the subject takes aspirin or not yields a statistically significant difference in outcome.

Notationally, ∥·∥denotes the vector *ℓ*_2_ norm or spectral norm for matrices, unless otherwise specified. ∥ *A* _*F*_ ∥ denotes the Frobenius norm of a matrix *A*. The operation diag(·) maps a length *n* vector *v* to an *n*×*n* matrix *A*, where all off diagonal entries of are 0 and *A*_*ii*_=*v*_*i*_ for *i* ∈[*n*]. *A* ^(*j*)^ denotes the *j*-th column of a matrix. When applied to a vector, 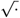 denotes the entrywise square root. **1** is taken as the all ones vector of appropriate dimension. Φ denotes the CDF of a standard Gaussian random variable. Finally, for a random variable we use 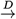 to denote convergence in distribution.

## 2. OASIS test statistic

The OASIS test statistic has a natural linear algebraic formulation, as we develop in Eq. (2). This statistic is computed using two input vectors ***f***∈ ℝ^*I*^ and ***c***∈ ℝ^*J*^, where a significant p-value will be obtained if the contingency table *X* ∈ ℕ^*I×J*^ can be well partitioned according to ***f*** and ***c***. These vectors should be thought of as row and column embeddings respectively, and can be generated with the assistance of metadata (if available), by random selection, or through an optimization framework we develop in Section 4. To compute the OASIS test statistic *S*, we first define the expected matrix *E* ∈ ℝ^*I×J*^ and the centered and normalized table 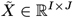:

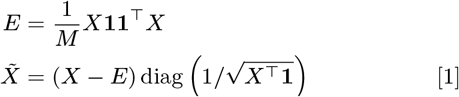

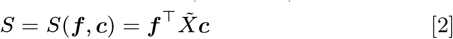

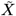 is normalized so that under the null, the variance of 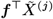 is constant across all *j* up to the dependence between *X*^(*j*)^ and *E*. This linearity enables the construction of a finite-sample valid p-value bound for fixed ***f*** and ***c***, which operate on the rows and columns of the table respectively. In preliminary work these vectors were chosen randomly (1), leading to a specific case of the OASIS test we call OASIS-rand, albeit with a looser analysis and worse p-value bound. In this work we formulate an optimization objective to construct ***f*** and ***c*** that identify latent structure, noting that other approaches based on experimental design or data-adaptive optimization may also be possible.

### A. Comparison with Pearson’s *X*^2^ test statistic

The linear algebraic formulation of Pearson’s *X*^2^ statistic highlights its fundamental differences from OASIS. *X*^2^ is computed as:

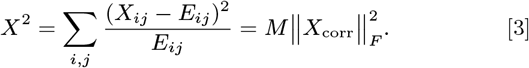

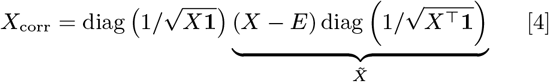

In contrast, OASIS’s p-value depends on *S*^2^ in Eq. (2), highlighting two main differences between *X*^2^ and OASIS. First, *X*^2^ left normalizes using empirical row frequencies, which from a technical perspective makes finite-sample analysis of the test-statistic difficult, and from a practical perspective upweights minor deviations in low count rows. OASIS treats rows and columns asymmetrically and only normalizes by column frequencies, which are given by the model. Second, *X*^2^ squares each residual, and then sums these quantities. OASIS computes a bilinear form of the residual matrix with ***f*** and ***c***, and squares the resulting sum. This allows residuals resulting from unstructured deviations to average out, focusing the power of the test on structured deviations from the null. We make this intuition precise in Section 4.B

Since Pearson’s *X*^2^ test does not provide guidance for the reason the null is rejected, practitioners commonly employ correspondence analysis for this task (8). Correspondence analysis studies the matrix of standardized residuals *X*_corr_ defined in Eq. (4). This method computes the singular value decomposition of *X*_corr_, and projects rows and columns along the first few principal vectors to obtain low dimensional embeddings for both the rows and the columns. As we show, OASIS provides a statistically grounded and equally interpretable alternative to this approach by analyzing 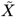 which in out experiments better identifies latent structure.

In addition, the power of Pearson’s *X*^2^ test decays as the table size increases, as *X*^2^ is approximated as being *χ*^2^-distributed with (*I* −1)(*J*− 1) degrees of freedom under the null of row and column independence for an *I*×*J* table. This yields several important classes of alternative hypotheses where OASIS is predicted (and empirically shown) to have higher power than *X*^2^, such as time series or 2-group alternatives when the total number of counts *M* is small relative to *I*× *J* (details in Section 6.B).

## 3. Analysis of OASIS statistic

The bilinear form of OASIS’s test statistic admits both an exact asymptotic p-value, and a finite-sample p-value bound. OASIS’s design provides interpretation for rejection of the null through ***f*** and ***c***, and allows for the construction of an effect size measure which quantifies the magnitude of deviation from the null, deconfounding the total number of observations *M*.

### A. p-value bound

A preliminary version of OASIS was designed so that p-value bounds could easily be obtained via classical concentration inequalities (1); here we improve these bounds, derive the asymptotic distribution of the test statistic, and show that our newly derived finite-sample bounds have a matching form with the asymptotic p-value. For notational convenience we define the quantity *γ* which measures the similarity between the column embedding vector ***c*** and the vector of columns counts ***n*** = *X*^⊤^**1** as

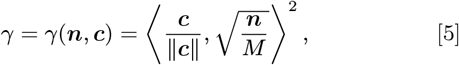

where we drop the dependence on ***n*** and ***c*** when clear from context. Observe that *γ*∈ [0, 1] by Cauchy-Schwarz. While the p-value bound can be computed for any ***f***, ***c***, we provide a constrained variant below for simplicity.

#### Proposition 1.

*Under the null hypothesis, for any fixed* ***f***∈ [0, 1]^*I*^ *and* ***c*** ∈ ℝ^*J*^ *with* ∥***c***∥ _2_ ≤ 1, *if γ <* 1, *the OASIS test statistic S* = *S*(***f***, ***c***) *satisfies*

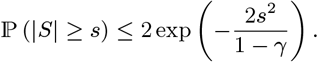

We provide an unconstrained version of this bound along with the proof details in Section 10.B, where we additionally note that for *γ* = 1, *S* = 0 with probability 1.

### B. Asymptotic normality

As intuition predicts, since the OASIS test statistic is the sum of independent increments, it converges in distribution to a Gaussian as the number of observations goes to infinity, as long as *γ*≠1. For fixed ***f*** and ***c***, define 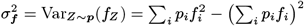, where ***p*** is the common row distribution under the null. Then, we can state the following asymptotic normality result.

#### Proposition 2.

*Consider any fixed* ***f*** ∈ ℝ^*I*^, ***c*** ∈ ℝ^*J*^, *probability distribution* ***p*** ∈ Δ^*I*^ *with* 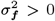, *and any sequence of column counts* 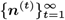 *where each* ***n***^(*t*)^ ∈ ℕ^*J*^, *with* 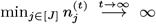 *and γ*(***n***^(*t*)^, ***c***) *<* 1 *for all t. Then, the random sequence of OASIS test statistics* 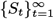, *where S*_*t*_ = *S*(*X*_*t*_, ***f***, ***c***) *and* 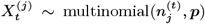 *independently across j, t, satisfies*

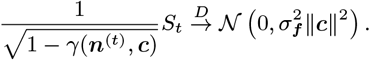

The intuition for this result is that each entry of 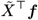 is asymptotically distributed as 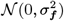, up to the negative correlation stemming from the unknown *μ* = 𝔼_*Z∼****p***_[*f*_*Z*_]. Then,*S* is a linear combination of 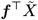 with weights ***c***, and so *S* has variance essentially 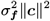, up to the (1−*γ*) scaling. As a direct corollary, by Slutsky’s theorem an asymptotically valid p-value can be constructed using the sample variance 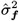, based on the empirical row distribution 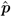.

#### Corollary 2.1.

*Under the conditions of Proposition 2*,

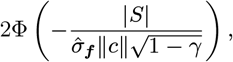

*is an asymptotically valid p-value*.

Using standard gaussian tail bounds this asymptotic p-value can be upper bounded as

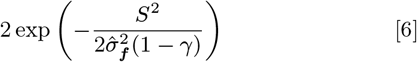

for ∥***c***∥_2_ = 1. This exactly matches the finite-sample upper bound derived in Proposition 1 up to *σ*_***f***_, where the finite-sample bound upper bounds the variance of *f*_*Z*_, a [0, 1]-valued random variable, as 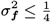.

### C. Effect Size

Through ***f*** and ***c***, OASIS assigns scalar values to each row and column. The preliminary version of OASIS (1) assigned each sample to one of two groups by utilizing *c*_*j*_ =±1, which is simply a partition of the samples into two groups. Building on this, we formalize an effect size measure for OASIS, which is computed as the difference in mean as measured by ***f*** between the two sample groups induced by the sign of *c*_*j*_. Defining *A*_+_ ={ *j* : *c*_*j*_ *>* 0} and *A*_*−*_ ={ *j* : *c*_*j*_ *<* 0 }, the effect size is computed as

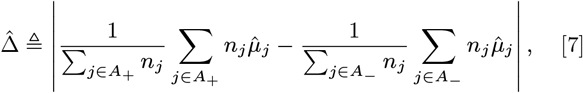

where 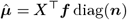. Defining *δ*_TV_(·, ·) as the total variation distance between two probability distributions, and 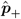 as the empirical row distribution over samples in *A*_+_ (similarly for 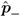), the effect size measure 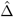 in Eq. (7) satisfies for all ***f***∈ [0, 1]^*I*^ that 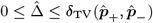.

The effect size is bounded between 0 and 1, and cannot be too large if the empirical distributions on the rows is similar between the two groups. The proof of the relationship between effect size and total variation distance, and the motivation for this effect size measure stemming from a simple two group alternative, are detailed in Section 10.D. Empirically, as we later show, this effect size measure allows OASIS to prioritize scientifically interesting tables.

## 4. Optimization-based approach to constructing *f, c*

Discussion heretofore has focused on studying OASIS’s test statistic for a fixed ***f*** and ***c***. The natural question is then, how to choose ***f*** and ***c***? Many options are possible. Here, we focus on an intuitive method, `OASIS-opt`, that partitions the observed counts into independent “train” and “test” data sets, using the training data to construct ***c*** and ***f*** that optimize the p-value objective, and the held-out test data to compute a statistically valid p-value (bound)as shown in Figure 2A, with analysis in Section 10.E.

**Fig. 2.**
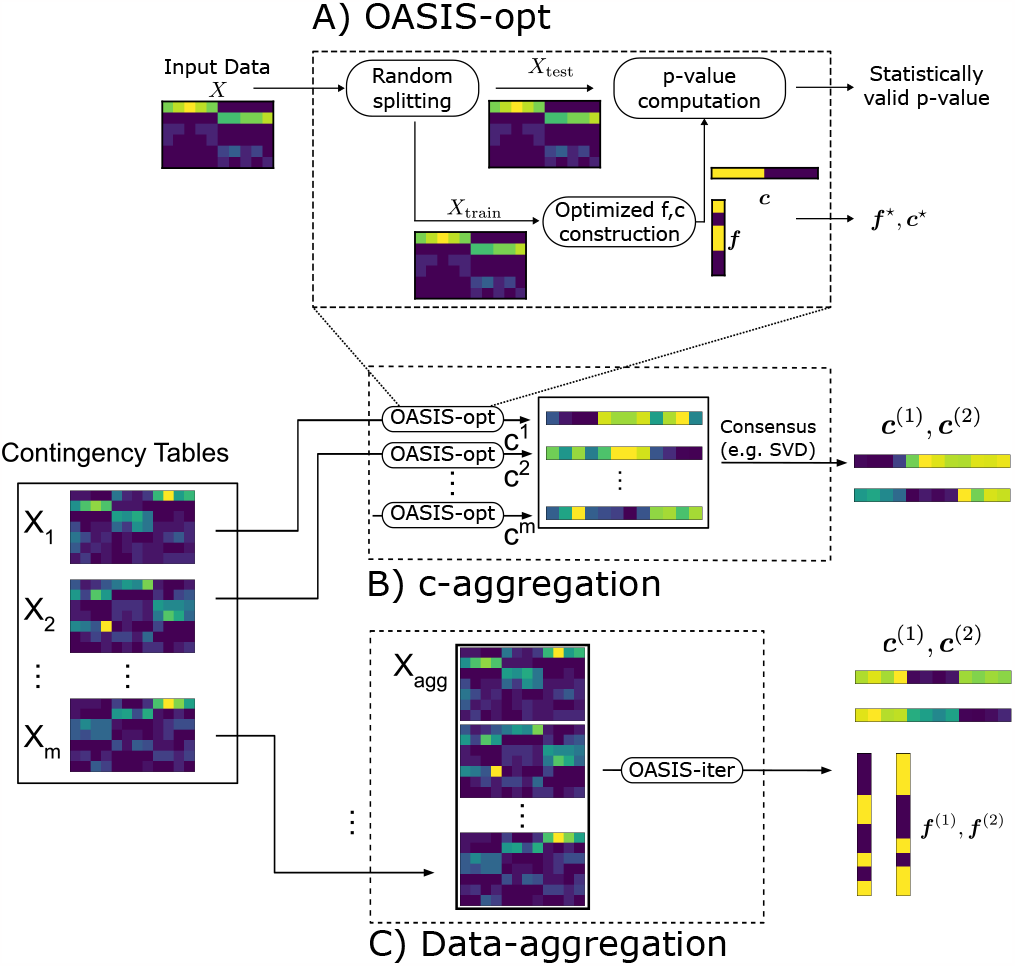
Figure showing the algorithms and methods we build using the OASIS framework. **A)** `OASIS-opt`, the primary instantiation of OASIS used in this work. `OASIS-opt` employs data-splitting to generate optimized data-dependent ***f*** and ***c***, before generating a statistically valid p-value bound on the held out test data. **B** and **C** depict two algorithms we propose for inferring latent structure from a collection of tables defined on the same set of columns. As building blocks, we use `OASIS-opt` for ***c***-aggregation, and `OASIS-iter` (Algorithm 3) for data-aggregation. **B) *c***-aggregation (Algorithm 2) performs inference on each table marginally using `OASIS-opt` and aggregates the resulting ***c*** embeddings. **C)** Data-aggregation (Algorithm 1) stacks the contingency tables into one large matrix *X*_agg_, and performs iterative analysis on this aggregated table using `OASIS-iter`.

**Fig. 3.**
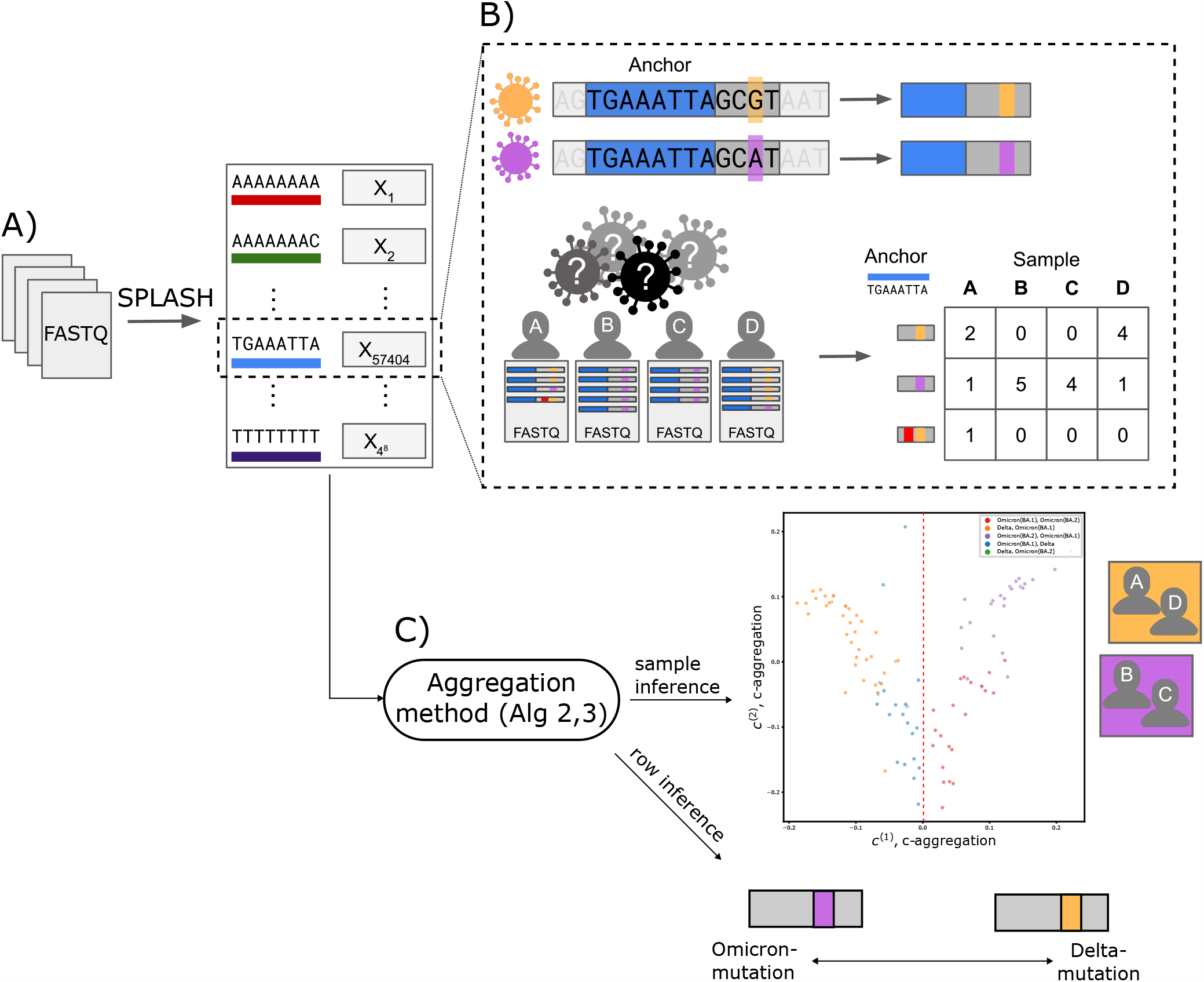
**A)** SPLASH (1) generates contingency tables from genomic sequencing data, here FASTQ files, for all 4^*k*^ possible anchor k-mers (length *k* genomic sequences). **B)** Shown in greater detail is the process for one specific anchor, TGAAATTA. This anchor highlights a mutation between two strains of SARS CoV-2, Omicron (purple) and Delta (orange). Below, viral sequencing data from 4 individuals (samples) infected with COVID is shown. However, it is a priori unknown which strain each individual was infected with, and no reference genome is available. For the fixed anchor sequence (shown in blue), SPLASH counts for each sample the frequency of sequences that occur immediately after (target sequence), and generates a contingency table, where the columns are indexed by the samples and the rows are indexed by the sequences. Shown in B is one read in sample A which underwent sequencing error, highlighted in red, and thus yielded an additional discrete observation – a sequence – resulting in an extra row. Sequencing error leads to tables with many rows with low counts. Note that we cannot know a priori which rows of this table are due to sequencing error, as we are simply observing raw sequencing data. **C)** The contingency tables generated by SPLASH are defined over the same set of samples (patients), so we can use these tables to jointly infer sample origins. Plot shown depicts the results of ***c***-aggregation on SARS-CoV-2 data (6), perfectly predicting whether a patient has Delta or not, and yielding high predictive accuracy (92%) for subvariant classification (Omicron BA.1 versus BA.2). Data-aggregation can also be used to predict the strain of mutated targets, with a 93% correct classification accuracy of whether a target was Delta or not. In the depicted toy example, this would correspond to grouping targets and individuals by strain as shown.

**Fig. 4.**
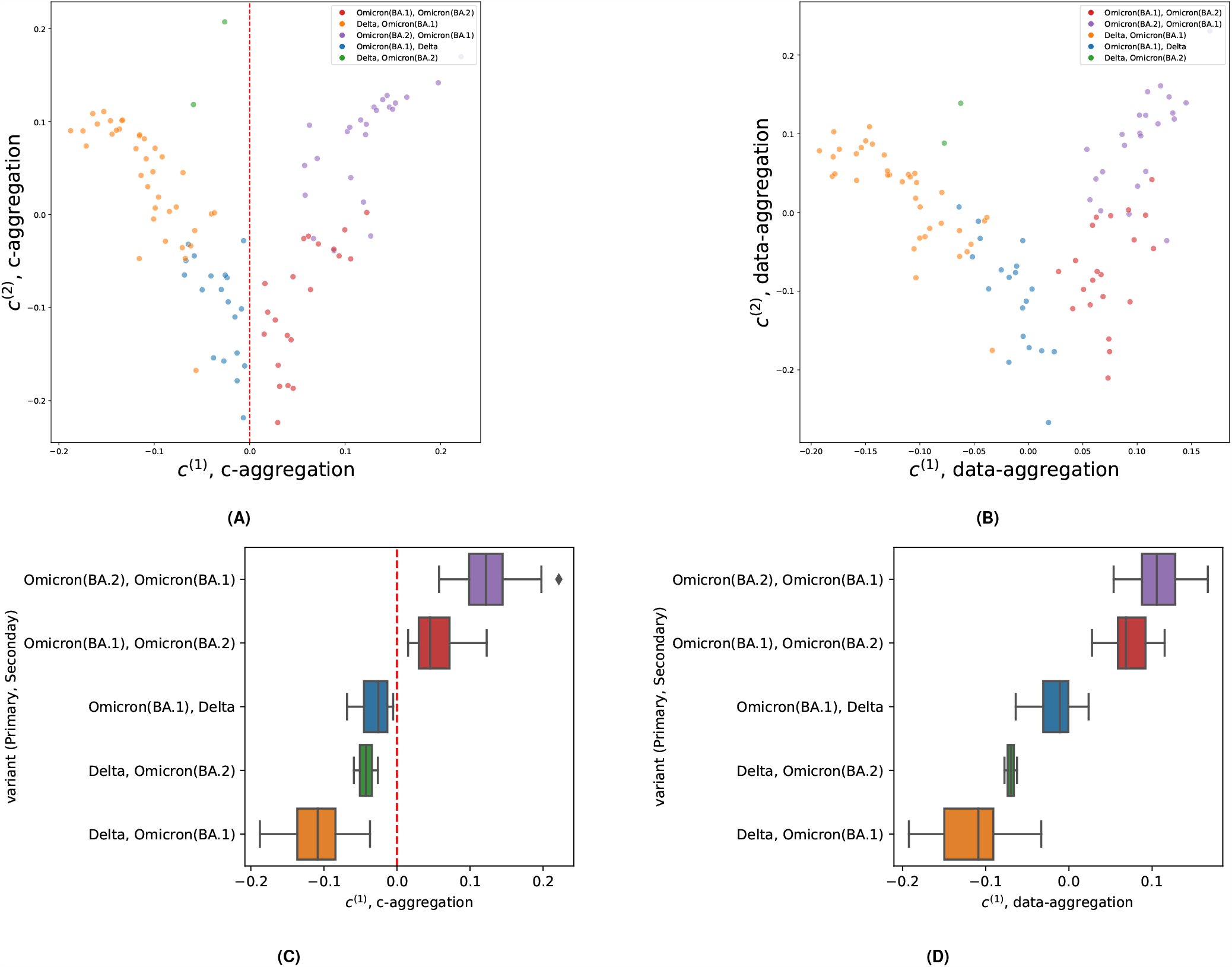
Analysis of SARS-CoV-2 coinfection data (6). Tables were generated by SPLASH (1) and tested with `OASIS-opt`. **(A)** and **(C)** depict the results of ***c***-aggregation (Algorithm 2). **(A)** shows the generated 2D embeddings, which perfectly classify whether a patient has Delta or not at ***c***^(1)^ ≷ 0, highlighted in **(C). (B)** and **(D)** depict the results of data-aggregation (Algorithm 1). Perfect separation of Delta vs non-Delta samples, no longer at ***c***^(1)^ ≷ 0. SARS-CoV-2 analysis details in Section 13.

**Fig. 5.**
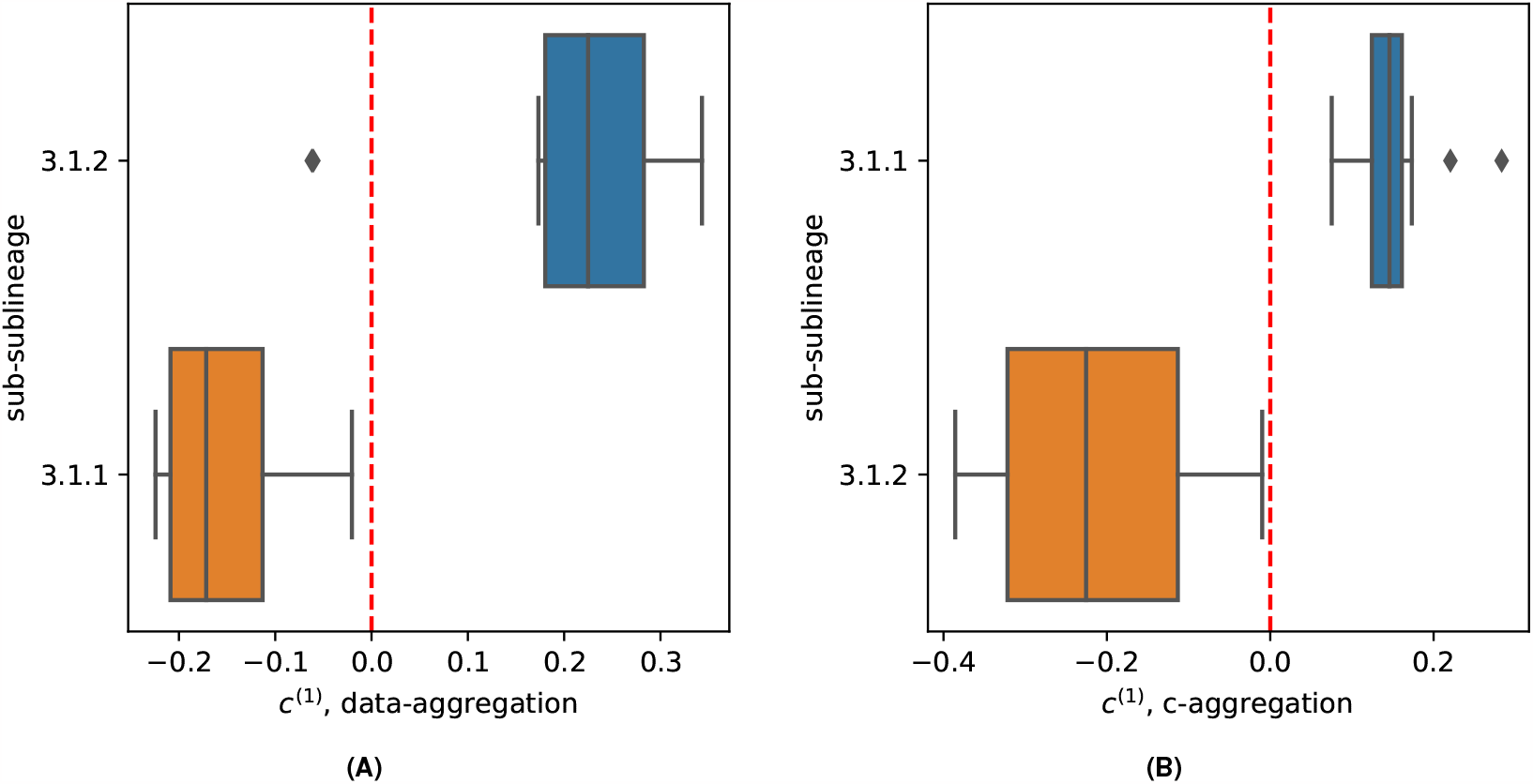
Interpretation of OASIS-rejected null from M. Tuberculosis data (6). Tables were generated by SPLASH (1) and tested with OASIS. **(A)** shows the generated 1D embeddings from ***c***-aggregation (Algorithm 2), which perfectly classifies patients based on sub-sublineage at ***c***^(1)^ ≷ 0. **(B)** depicts the results of data-aggregation (Algorithm 1). Two samples are misclassified (visually, one on top of the other), but with a much larger margin for the rest. 2D plots with ***c***^(2)^ shown in Figure 25.

### A. Minimizing the p-value bound

Analyzing the finite-sample p-value bound, we begin by simplifying the optimization objective, observing that the p-value bound is minimized by maximizing the test statistic. Concretely, considering the two optimization problems:

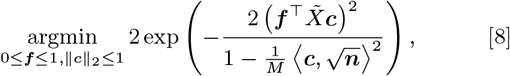

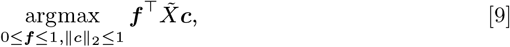

we prove the following lemma (details in Section 11.A).

#### Lemma 1.

*The set of optimal solutions to* Eq. (9) *is contained within the set of optimal solutions to* Eq. (8).

Thus, to identify an optimal solution to Eq. (8), it suffices to optimize Eq. (9). Since the objective is bilinear the maximum value must be attained at a corner point. Further, since ∥ · ∥_2_ is self-dual, Eq. (9) can equivalently be written as

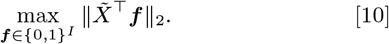

In graph contexts, a similar combinatorial optimization problem arises in the determination of the max-cut, which is known to be NP-complete. In particular, Eq. (10) is in general APX-hard, meaning that no polynomial-time approximation scheme exists for arbitrarily good approximations (14). For small *I*, the problem can be solved exactly by enumerating all 2^*I*^ possible ***f***. For large *I* more sophisticated algorithms can be used, such as SDP relaxations (discussed further in Section 11.E). However, due to the structure of 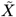 and the biconvex nature of Eq. (9), alternating maximization is computationally efficient and yields empirically good performance.

Alternating maximization converges to a local maxima, computing iterates as 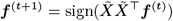, implicitly computing 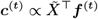. Due to the non-convexity of the overall objective, several random ***f*** initializations are used in practice, which we encapsulate in an algorithm called `OASIS-opt` (additional discussion and analysis details in Section 11). Note that there are two sources of randomness in `OASIS-opt`; statistical randomness in data-splitting, and the randomn ***f*** initializations for approximately solving the inner maximization problem. These are distinct, with the former being necessary and the latter being a computational tool to improve alternating maximization. We show in Figure 8 how utilizing additional random train / test splits and applying Bonferroni correction to the generated p-value bounds improves empirical performance, ensuring rejection of highly significant tables, at the expense of a higher burden of multiple hypothesis correction. We discuss these aspects further in Section 14.A.

### B. Relation to the Singular Value Decomposition

Examining the optimization problem in Eq. (9), observe that if ***f*** were *ℓ*_2_ constrained then the optimal solution would take ***f*** (resp. ***c***) as the principle left (resp. right) singular vector of 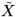, where the optimal value would be the maximum singular value by the variational characterization of the SVD. `OASIS-opt` provides an alternative decomposition of a contingency table to the ubiquitous SVD, within a disciplined statistical framework. While the SVD has a statistical interpretation in some parametric settings, `OASIS-opt`’s alternative decomposition provides a fresh and potentially more statistically tractable and interpretable decomposition.

As in Eq. (3), the *X*^2^ test statistic can be expressed as 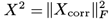. As 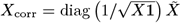 this can be bounded as a function of 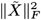, and can be approximated as 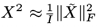 when all rows have similar counts. Comparatively, the OASIS test statistic attains a maximum value (up to the *ℓ*_2_ as opposed to *ℓ*_∞_ constraint on ***f***) of 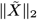. Note that the *ℓ*_2_ ball is contained within the *ℓ*_∞_ ball, and so the optimal value of the OASIS test-statistic is in fact lower bounded by 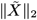. The Frobenius norm is the sum of squared singular values, and so *X*^2^ is essentially summing over all possible directions of deviation. This makes it powerful, but overpowered against uninteresting and unexplainable alternatives such as those with a flat spectrum. OASIS on the other hand computes significance by projecting deviations from the expectation in a single direction. Intuitively, this denoises all the lower signal components, and ensures that OASIS remains robust to biological noise, and yields interpretable results. This is exactly why *X*^2^ fails to distinguish between the tables in Figure 1C, where we plot the spectra of the strong and weak signal tables in Figure 9.

An SVD of 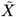 or *X*_corr_ offers one approach to generating ***c, f*** for inference with OASIS, however this construction will in general yield suboptimal ***c*** and ***f***. This choice gives worse p-value bounds (as the SVD is optimizing a fundamentally different objective), with empirically less meaningful ***f*** and ***c*** than `OASIS-opt`, as shown for a toy example in Figure 6, and for SARS-CoV-2 data in Table 3. One reason for this is that the OASIS p-value bound constrains the *ℓ*_*∞*_ norm of ***f*** rather than *ℓ*_2_ as in the SVD. Directly optimizing the objective (with an *ℓ*_*∞*_ constraint) naturally yields improved p-value bounds. `OASIS-opt` provides a promising alternative to correspondence analysis, but further experiments and theoretical analyses are required to validate the quality of these embeddings in more general settings.

**Fig. 6.**
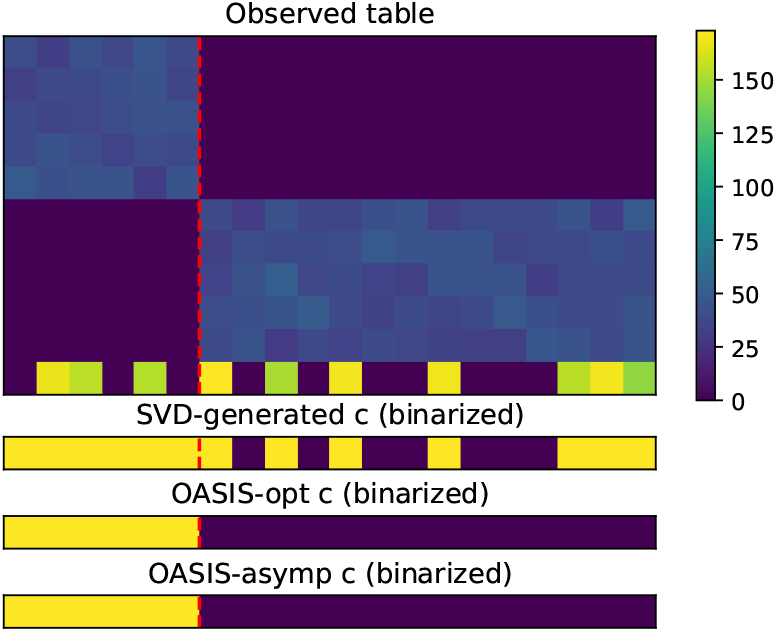
SVD does not identify the visually clear planted structure in the shown toy example. Shown at top is the contingency table *X*, which displays a clear 2 cluster behavior (the left 6 columns versus the right 14 columns). The last row should be considered noise, as this signal does not match with any other row expression. However, taking the SVD of 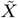 (the centered and right normalized table), the principle right singular vector does not well predict the planted structure. OASIS, both from its asymptotic-optimized ***c*** and its finite-sample bound optimized ***c***, both perfectly identify the latent clustering. The raw values for each methods identified ***c, f*** are plotted in Figure 7.

**Fig. 7.**
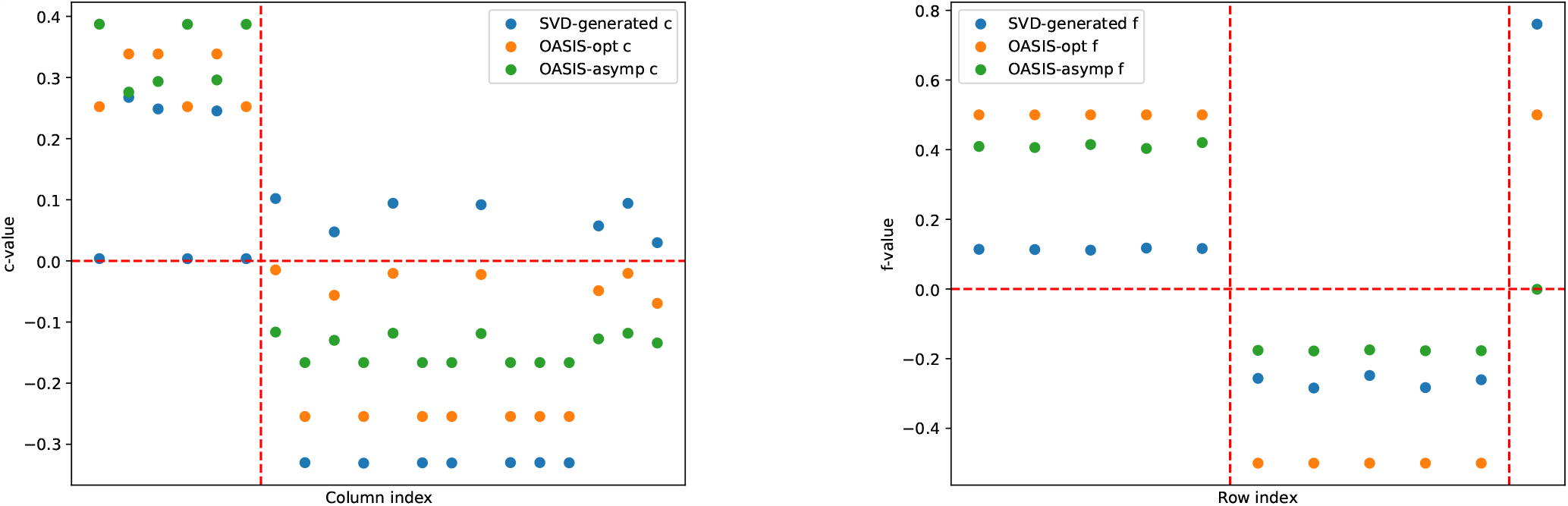
Plots of ***f***, ***c*** vectors identified by SVD, `OASIS-opt`, and OASIS-asymp. `OASIS-opt` ***f*** is shown with .5 subtracted, so that is centers around 0 and can be visually compared with the other ***f*** vectors. On the right, the SVD-based ***f*** does identify the row groupings, but utilizes only half the dynamic range for the first 10 rows. OASIS more clearly identifies the row structure, with the asymptotic objective forcing the bottom row to a value of 0. `OASIS-opt` utilizes the full dynamic range for the first 10 rows. Examining the ***c*** plot on the left, we see that the SVD-identified ***c*** cannot identify the latent structure, as for example *c*_1_ *< c*_7_, while *c*_1_ should belong in the same cluster as *c*_2_ which has a large positive value. `OASIS-opt` clearly identifies the clustering in a well separated manner, and the asymptotic optimized ***c*** yields even greater separation (as it fully discounts the last “noisy” row).

**Fig. 8.**
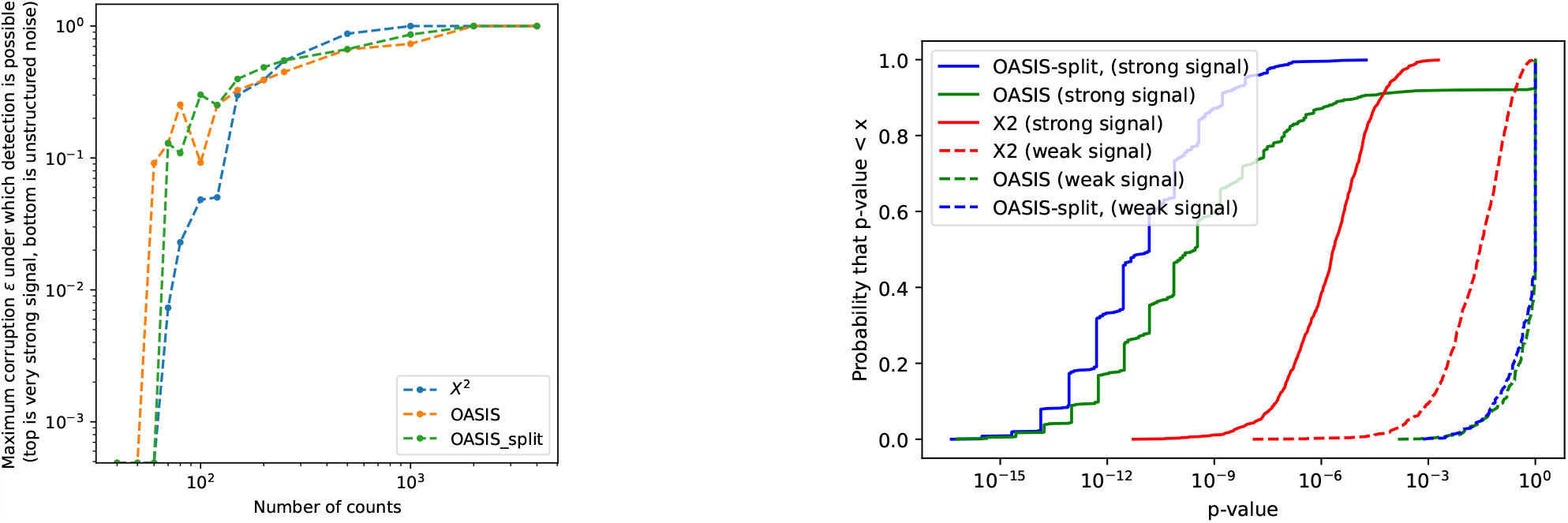
Following Figure 1, this plot shows the performance of `OASIS-opt` and *X*^2^ over the class of alternatives defined in Eq. (43). The plot shows an empirical achievability curve, where the y-axis indicates how structured versus unstructured the alternative is, and the x-axis indicates the total number of counts in the table. The more up and to the left a curve is, the better. For any alternative (any *ϵ*∈ [0, 1] for our model), a statistical test should reject the null with enough observations. Here, we show that empirically, OASIS is more likely to reject highly structured tables with lower counts, whereas Pearson’s *X*^2^ test is more likely to reject unstructured tables with moderate counts. `OASIS-opt` with 5 random data splits is shown. The flat behavior between adjacent data points for `OASIS-opt` is due to the data-splitting: an increase in number of observations does not necessarily translate to an increase in number of observations in the test data, e.g. going from 7 to 8 observations per sample. The right plot shows the subplot in Figure 1A, additionally showing curves when 5 random splits are used for `OASIS-opt`. As can be seen, this removes the tail of the `OASIS-opt` p-value bounds, improving performance in this setting. The jagged behavior in the plot is due to the finite number of observations (*M* = 100 in this plot), leading to a finite number of values the optimized test-statistic will take, and so even aggregating over 1000 trials, the curve does not smooth out as it does for *X*^2^, which can take many more possible values.

**Fig. 9.**
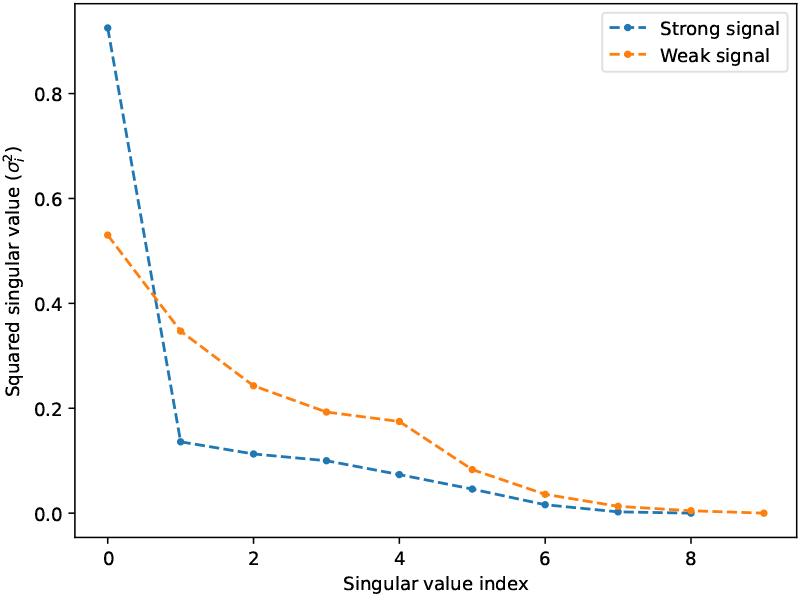
Spectrum of *X*_corr_ for the two example tables in Figure 1C. *X*^2^ yields a similar p-value for the two tables, as the sum of the squared singular values is similar. OASIS prioritizes the “strong signal” table with the more spiky spectrum.

### C. Minimizing the asymptotic p-value

While optimizing the finite-sample p-value bound yields a combinatorially hard optimization problem, the asymptotic p-value can be optimized efficiently. As we detail in Section 11.D, an optimal pair ***f***, ***c*** which minimize the asymptotic p-value objective given in Corollary 2.1 must satisfy 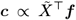 and 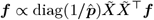. Thus, an optimal ***f*** can be computed as a principle right eigenvector of the asymmetric matrix 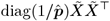. This provides an interesting contrast with the classical SVD; where correspondence analysis would take as the row embeddings a principle eigenvector of the symmetric matrix 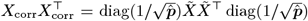. Note that the optimal ***f*** is non-binary in this setting; the discrete result in the finite-sample case is due to the use of Hoeffding’s inequality in the construction of the p-value bound. Alternative optimization problems derived from finite-sample bounds based on Bernstein’s inequality, for example, would also yield non-binary optimal ***f***.

## 5. statistical framework for subgroup classification with OASIS-opt

OASIS at its core is a statistical test for contingency tables, against the null of row and column independence. However, after the null has been rejected, many followup questions often remain. One natural question, after the samples have been found to violate the null when combined using ***c***, is whether there exist different ways of grouping samples to still yield statistically significant deviations. A more general question is whether, studying many tables defined on the same sets of columns, there is some global inference that can be drawn regarding the underlying clustering of the samples.

### A. Iterative analysis of contingency tables with OASIS-iter

Addressing the first question of iterative testing of a single contingency table, we first define a method `OASIS-perp`, which takes in a table *X* and a set of vectors {*v*^(*k*)^ }, and optimizes the objective in Eq. (11).

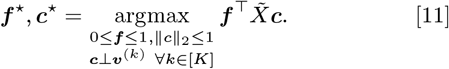

This objective is identical to that of `OASIS-opt`, except that the optimizing ***c*** is constrained to be orthogonal to all vectors in the input set {*v*^(*k*)^ }. This added constraint retains the biconvexity of the original problem, enabling the use of alternating maximization.

To iteratively analyze a table, we propose a simple wrapper on top of `OASIS-perp` called `OASIS-iter`. `OASIS-iter` decomposes a contingency table by first identifying ***f*** ^(1)^, ***c***^(1)^ from `OASIS-opt`, then iteratively identifying ***f*** ^(*i*)^, ***c***^(*i*)^ optimizing Eq. (11) subject to the identified ***c***^(*i*)^ being orthogonal to {***c***^(*j*)^}_*j<i*_. This provides a statistical stopping criterion for cluster identification; for each (***c***^(*i*)^, ***f*** ^(*i*)^) pair computed, we can compute OASIS’s p-value bound, and once the obtained *p*^(*i*+1)^ exceeds the desired significance level (e.g. 0.05), the number of clusters can be estimated as *i*. Each ***c***^(*i*)^ outputted by this procedure represents an orthogonal direction along which this table can be partitioned so as to yield a significant partitioning. The algorithm is made explicit in Algorithm 3, the derivation for this orthogonally-constrained optimization and its alternating maximization updates are in Section 11.C, and simulations on synthetic data are in Section 14.F.

### B. Data-aggregation

With this new primitive of iterative analysis of a single table, one candidate approach to identifying clusters across multiple tables is by aggregating data across many tables, and running `OASIS-iter` to iteratively identify underlying clusters. We design a method for this task called data-aggregation (Algorithm 1), shown graphically in Figure 2B. Data-aggregation takes as input a set of tables, which are defined on the same set of columns and should have shared

#### Algorithm 1 Data-aggregation

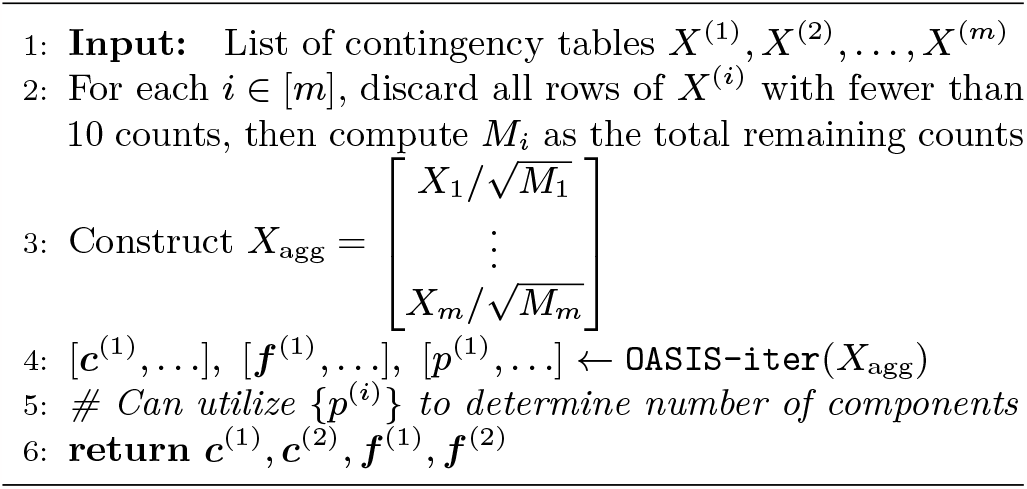

#### Data-aggregation

latent structure. In our setting, these tables are generated by SPLASH, where we filter `OASIS-opt`’s calls for tables with a large effect size, many observations, and a significant p-value bound after performing Benjamini-Yekutieli (15) correction across a much larger list of tables outputted from SPLASH (1). These tables are then vertically concatenated into one larger table, *X*_agg_, upon which `OASIS-iter` is run. For ease of visualization we only utilize the first two outputted ***c***^(1)^ and ***c***^(2)^ as the 2D sample embeddings.

Note that this aggregated table need not satisfy the contingency table null, due to the correlation in entries from the different sub-tables. Algorithmically, we discard rows in this table with fewer than 10 observations, to minimize the computational burden. Additionally, to avoid having a single high-count sub-table dominate all the others in this computation, we normalize the counts in each sub-table by 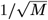, the total number of counts in that sub-table.

### C. c-aggregation

Due to the statistical dependencies and practical issues with normalizing and aggregating tables before performing statistical testing, a more straightforward clustering approach may be desired. To this end, we propose a second method which independently identifies 1D embeddings for each table and then aggregate these results. We call this method ***c***-aggregation (Algorithm 2), which utilizes `OASIS-opt` to generate a vector ***c*** for each table, collects these vectors into a matrix *C*, and then computes a low dimensional embedding for the samples, e.g. with an SVD, as shown in Figure 2C. As before, the tables used as input could be SPLASH outputs filtered for those with a large effect size, many observations, and a significant post-correction p-value bound.

In the construction of *C*, entries may be missing from the matrix, due to samples not appearing in all tables. We impute these with a value of 0 for simplicity, but more sophisticated approaches are possible. Defining the *i*-th right singular vector

#### Algorithm 2 *c*-aggregation

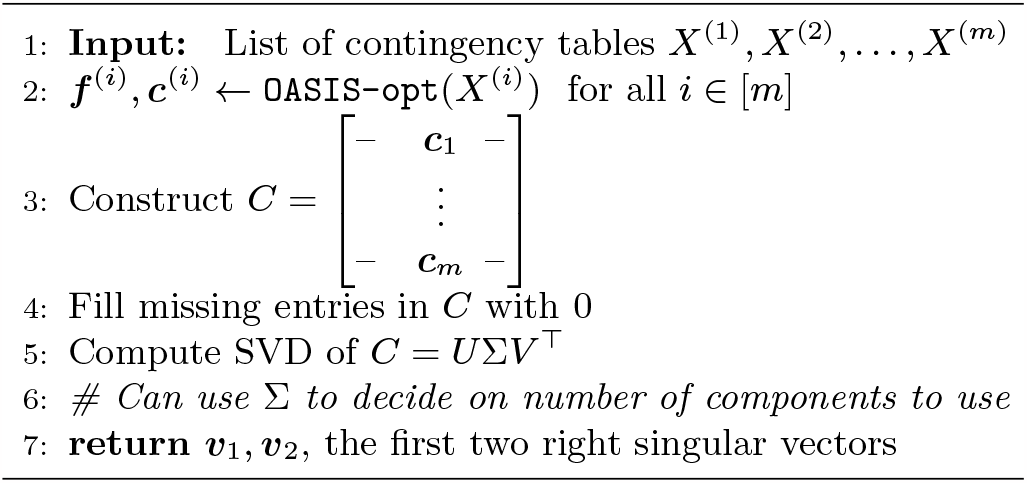

of *C* as ***c***^(*i*)^, ***c***^(1)^ is mathematically equivalent to finding a vector which maximizes the average squared inner product with each ***c***, discussed in Section 13.A.

## 6. Numerical results

Many problems in modern data science, including in genomics, map to contingency tables. Analysis of contingency tables in genomics applications, such as those that treat sequences or genes as counts, is core to biological inference. In these applications, noise is often introduced, sometimes without a good parametric model. We now show that OASIS is robust to some important classes of noise models, and that OASIS has substantial power against several classes of alternative hypotheses of interest. We also show that OASIS can be used to perform classification tasks not possible with Pearson’s *X*^2^, enabling reference-free strain classification in SARS-CoV-2 and M. Tuberculosis. For OASIS, we show the performance of `OASIS-opt` as well as `OASIS-iterOASIS-rand`, utilizing the improved p-value bounds presented in this work.

### A. Empirical robustness against uninteresting alternatives

Next generation sequencing data is commonly treated as matrices of discrete counts, for example with single-cell RNA sequencing data often represented as a cells by genes counts matrix. While the statistical null tests whether observations in each sample are identically distributed, biochemical noise introduced during sampling generates overdispersed counts, violating this null. The field has converged on modeling such data with negative binomial distributions (16, 17). Probabilistically, a negative binomial random variable is equivalent to a Poisson random variable with a random, gamma-distributed, mean. In terms of the observed table, this would yield an alternative probabilistic model where *X*_*ij*_ ∼Pois(Λ), Λ∼Γ(*λ/θ, θ*), *λ* is the expected number of observations, and under the true null *X*_*ij*_ ∼Pois(*λ*) counts would be observed (*θ* = 0). This does not satisfy the contingency table null for *θ >* 0, but also does not represent biologically meaningful deviation.

To test the robustness of OASIS and *X*^2^, we simulate this uninteresting alternative in Table 1, showing the fraction of rejected tables (for *α* = 0.05) at the level of negative binomial overdispersion predicted by the sampling depth (16). As expected, Pearson’s *X*^2^ rejects nearly all tables generated under this negative binomial sampling model, while OASIS retains robustness across a wide range of parameters, due to the unstructured deviations. For all tests, the rejection fraction is monotonic in the number of rows and columns. Taking a table with a mean of *λ*≈129 observations per sample with a uniform row distribution, for 20 rows and 10 columns `OASIS-opt` rejects 7.5% of tables, while Pearson’s *X*^2^ test rejects 95.7% of tables. Additional experiments and simulation details in Section 14.B.1.

**Table 1.** Power against null corrupted by negative binomial noise, as modelled for single-cell sequencing data (16). Uniform target distribution, full plots and details in Section 14.B.1.

| num rows | num cols | $X^2$ | OASIS-rand | OASIS-opt |
| --- | --- | --- | --- | --- |
| 5 | 10 | 0.143 | 0.003 | 0.010 |
|  | 50 | 0.318 | 0.006 | 0.019 |
|  | 400 | 0.318 | 0.006 | 0.019 |
| 20 | 10 | 0.957 | 0.054 | 0.075 |
|  | 50 | 1.000 | 0.065 | 0.193 |
|  | 400 | 1.000 | 0.078 | 0.874 |
| 100 | 10 | 1.000 | 0.947 | 0.996 |
|  | 50 | 1.000 | 0.996 | 1.000 |
|  | 400 | 1.000 | 0.998 | 1.000 |

More generally, an applied task in genomics and other applications is to reject the null only when it is “interesting”. Below, we refer to an alternative as uninteresting if it can be explained by a small number of outlying matrix counts, or from sampling from a set of distributions {***p***^(*j*)^ } where{ ***p***^(*j*)^ }is the row distribution of the *j*-th column, and ∥ ***p***^(*j*)^ − ***p***^(*k*)^ ∥ _1_ *< ϵ* for all *j, k*. Such distributions, while statistically deviant from the null, represent small effects that may be due to contamination or equipment error, and are not a priority to detect. OASIS provides robustness against these sources of unstructured noise that represent biologically uninteresting alternatives. We characterize this with simulations against *ℓ*_1_ corruption of each individual column’s probability distributions in Section 14.B.2.

### B. Power against simulated alternatives

OASIS has substantial power (in some regimes more than *X*^2^) against large, structured deviations from independence, such as when the samples can be partitioned into two groups with nearly disjoint supports. These examples arise in important applications such as detecting viral mutations, recombination in B cells that generate antibodies–V(D)J recombination–and differentially regulated alternative splicing, among others.

We show this empirically in Section 14.C, and provide approximate power calculations in Section 15 that corroborate these numerical results. Approximate power calculations are derived, for a simulated alternative, by considering the toy setting where we observe the expected underlying alternative matrix. Concretely, for sample *j* with row probabilities ***p***^(*j*)^, we assume that we observe *X*^(*j*)^ = *n*_*j*_ ***p***^(*j*)^ instead of the random draw *X*^(*j*)^ ∼multinomial(*n*_*j*_, ***p***^(*j*)^). Analyzing OASIS and *X*^2^ in this deterministic setting, we show that OASIS has power comparable to and in some regimes exceeding *X*^2^, in particular when the number of rows is large, as shown in Table 2. We conjecture that while OASIS may not exhibit the optimal asymptotic rate against certain classes of alternatives (as shown by the unique per-sample expression setting, motivated by V(D)J recombination), it does have power going to 1 as the number of observations goes to infinity across a broad class of alternatives.

**Table 2.**
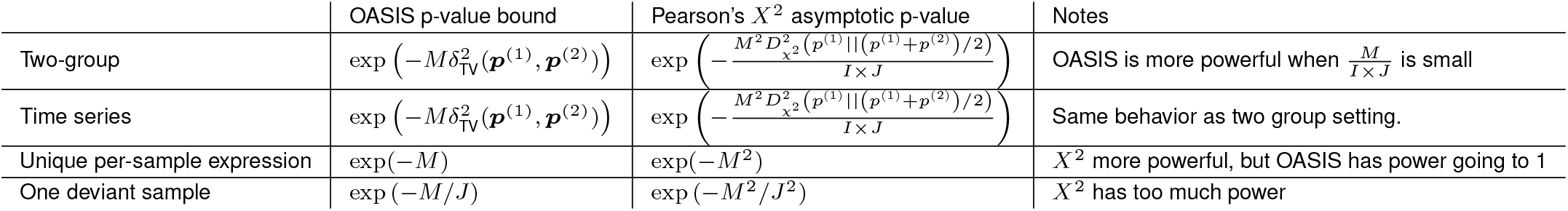
Summary of approximate power calculations in Section 15. Constants omitted for clarity.

### C. SARS-CoV-2 variant detection

To illustrate OASIS’s performance we study a public dataset of SARS-CoV-2 coinfections generated in France, which sequences patients during a period of Omicron and Delta co-circulation (6). We show that OASIS detects variants in SARS-CoV-2 by mapping sequencing data from each of 103 patients’ nasal swabs to a contingency table via SPLASH, as described in (1). SPLASH generates a contingency table for each length *k*-subsequence present (called an anchor *k*-mer) from genomic sequencing (Figure 3). A statistical test with good scientific performance will identify anchors near known mutations in the SARS-CoV-2 genome that distinguish Omicron and Delta. Data processing details are deferred to Section 13.

OASIS and *X*^2^ yield substantially different results on the 100,914 tables generated by SPLASH. We demonstrate the improvement in biological inference enabled by OASIS, utilizing as a measure of biological relevance for each method’s called tables how well these calls can predict sample metadata. For each sample, we have associated metadata indicating the clinical ground truth of whether the patient (sample) was infected with Delta. For each table called by `OASIS-opt`, we use as its 1D sample embedding the vector ***c***, taking the sign of each entry to generate a two-group partitioning. The measure of biological relevance used is then computed as the absolute cosine similarity, 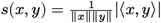, between the sample metadata and the sign of ***c***. This corresponds to the fraction of agreed upon coordinates in *x, y* (up to a global sign flip). For *X*^2^ this process is mirrored, where instead of ***c***, the principle right singular vector for correspondence analysis is used.

Out of the 100,914 tables, OASIS rejects 28,430, and Pearson’s *X*^2^ test rejects 71,543 tables. However, when the tables that *X*^2^ rejects are decomposed, they do not yield signal that correlates well with the ground truth; examining quantiles of the two empirical distributions of absolute cosine similarities, the 0.5 and 0.9 quantiles of these distributions are 0.22 and 0.66 for OASIS, as opposed to 0.10, 0.52 for *X*^2^. We show in Figure 21 that for these significant tables, correspondence analysis yields similarly high correlation, indicating that it is the improved detection of OASIS, rather than the handicap of correspondence analysis versus OASIS’s ***c***, that is yielding the larger similarities. In addition, of the 389 anchors that OASIS calls that *X*^2^ does not, the 0.5 and 0.9 quantiles of the absolute cosine similarity between these binarized ***c*** and the underlying ground truth of whether a patient has Delta are 0.15 and 0.76, indicating that many biologically significant tables that were missed by *X*^2^.

Analyzing all 16,611 tables rejected by *X*^2^ with a p-value of 0 (up to numerical precision), the 1D embeddings obtained from correspondence analysis had low absolute cosine similarity with the biological ground truth, with 0.5 and 0.9 quantiles of 0.087 and 0.40. Looking at specific rejected tables, we select two of the most significant tables rejected by *X*^2^ but not by `OASIS-opt`. Correspondence analysis yields embeddings with minimal correlation with the ground truth, 0.15 and 0.02 for the two tables selected, one of which visually appears to have just detected one deviating sample (Figure 19).

In contrast, `OASIS-opt` yields significantly more biologically relevant calls. ***c*** found by `OASIS-opt` have a significantly higher absolute cosine similarity with the clinical ground truth of whether the patient had a Delta infection. For the 2114 tables that `OASIS-opt` assigns a p-value bound of 0, the cosine similarity with ground truth has 0.5 and 0.9 quantiles of 0.69 and 0.79. We further see that `OASIS-opt`’s top significance calls have significantly higher concordance with clinical metadata. We consider for this analysis tables with effect size in the top 10% and total counts *M >* 1000, as these are predicted to delineate strains, yielding 2495 tables. Of these, the absolute cosine similarity between the identified ***c*** and whether the patient had a Delta infection has 0.5 and 0.9 quantiles of 0.72 and 0.82. In Figure 21, we show the ECDF of the absoltue cosine similarity of all 28,430 of `OASIS-opt`’s calls, and all 71,543 calls from *X*^2^, showing an order of magnitude difference in fraction of identified tables with e.g. *>* .6 cosine similarity.

Comparing with the original statistic used in SPLASH (1), OASIS-rand identifies 5932 significant tables. All except 48 of these are identified by `OASIS-opt`. The two most significant tables that are called by `OASIS-opt` (with an effect size in the top 10% and counts greater than 1000) and not by `OASIS-rand` are shown in Figure 20, both having a high absolute cosine similarity with sample metadata, 0.76, showing that the additional calls `OASIS-opt` is providing are biologically significant. We further show that OASIS provides calls beyond those possible with *X*^2^, showing in Figure 18 the 5 tables in the reduced anchor list above (effect size and counts filtered), which all have large cosine similarities with the ground truth metadata.

#### Data-aggregation

Running data-aggregation (Algorithm 1) on the SARS-CoV-2 dataset, yields important inference on both targets and samples. We select the first two components for simplicity of visualization, and show that the first component contains 57% of the total power as measured by 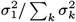, and with the second contained 76% of the cumulative power (full spectrum shown in Figure 22). Focusing first on samples, the first two ***c*** output by data-aggregation have high predictive power over strain information. A simple threshold on ***c***^(1)^ yields perfect prediction of whether a patient has Delta or not. The second vector, ***c***^(2)^, enables classification of Omicron subvariants. A linear predictor on ***c***^(1)^ and ***c***^(2)^ yields 94/103 (91%) accuracy in predicting whether the primary strain is Omicron BA.1, shown in Figure 4.

Running data-aggregation on these matrices also enables joint inference on targets from different tables. The outputted ***f*** ^(1)^ of this procedure has significant predictive power over whether a target corresponds to an Omicron mutation or not. We validate this by using Bowtie (18) to check whether a target noiselessly aligns to the Wuhan reference, a Delta assembly, an Omicron BA.1 assembly, or an Omicron BA.2 assembly (details in Section 13). Comparing with this ground truth vector of whether a target was classified as Delta or not, ***f*** correctly classified 4496/4836 (93%) targets. This is with no parameter tuning. The targets that are incorrectly classified have the potential to uncover novel biology, which is beyond the scope of this paper. Analyzing the first two anchors with targets that are considered misclassified, the first corresponds to a known Omicron deletion not present in the used reference genome, while the second perfectly identifies and predicts an annotated Omicron deletion, however due to parameter choices (SPLASH’s “lookahead distance” (1)) both targets perfectly map to the Wuhan reference. We provide alignments for these results in Figure 24.

#### C.2 c-aggregation

We additionally run ***c***-aggregation (Algorithm 2) on this SARS-CoV-2 dataset, restricting our attention to the first two right singular vectors of *C*. The first singular vector of the matrix *C* perfectly partitions individuals infected with the Delta strain from those that were not (at the threshold ***c***^(1)^ ≷ 0). The second singular vector differentiates between the Omicron BA.1 and BA.2 subvariants as shown in Figure 4 where the x-axis is the principle right singular vector and the y-axis is the second right singular vector. When tasked with predicting whether a sample has BA.1 as its primary variant, a simple linear classifier in this two dimensional space is able to correctly classify 95/103 (92%) of the samples. Further analysis details are in Section 13.A.

Since ***c***-aggregation provides inference on the samples, it can be used to indirectly provide inference on the targets. We show this by utilizing *ŷ* = sign(*X*_agg_***c***^(1)^) as a predictor of whether a patient is infected with Delta or not, which yields correct predictions on 4357/4836 targets (90%). This highlights the power of data-aggregation in performing joint inference directly on the rows.

Together, these analyses suggest that OASIS finds tables representing important biological differences, and can provide auxiliary information on samples (***c***) that can guide scientific inference within a disciplined framework. These features are absent from *X*^2^ or exploratory correspondence analysis.

##### D.M. Tuberculosis strain identification

We tested the generality of `OASIS-opt`’s inferential power to classify microbial variants in a different microorganism, the bacterium M. Tuberculosis. We processed data from 25 isolates from two sub-sublineages known to derive from the sublineage 3.1: 3.1.1 and 3.1.2 (Accession ID PRJEB41116 (19)). Currently, bacterial typing is based on manual curation and SNPs, is highly manually intensive, and requires mapping to a reference genome.

As with SARS-CoV-2, we utilized SPLASH to generate tables which we tested with `OASIS-opt`. 80,519 tables were called by `OASIS-opt` (corrected p-value bound), 258 with effect size in the 90-th quantile and total count *M >* 1000. To avoid biological noise, we preemptively filtered out tables with targets with repetitive sequences (*>* 10 repeated basepairs). `OASIS-opt` was run blind to sample metadata and the M. Tuberculosis reference genome.

##### D.1. Data-aggregation

We run data-aggregation, Algorithm 1, on the filtered Tuberculosis tables described above. The first identified vector ***c***^(1)^ yields a well separated partitioning in terms of sub-sublineage as shown in Figure 5A. Two samples are misclassified, with the rest being perfectly predicted by ***c***^(1)^ ≷ 0, yielding an accuracy of 23/25 (92%).

##### D.2. c-aggregation

We utilized ***c***-aggregation on the same set of filtered tables. The first singular value constitutes 49% of the power in the matrix, and the top 2 singular values comprise 72% of the spectrum’s power. A 1D embedding from the first singular vector leads to a perfectly separation of sub-sublineages, classifying at the threshold ***c***^(1)^ ≷ 0, as shown in Figure 5B.

## 7. Discussion

In this paper we proposed a framework for interpretable and finite-sample valid inference for contingency tables, describing several applications and extensions of OASIS. There are many interesting directions of ongoing and future work, some of which we discuss below. A known failing of *X*^2^ is its inability utilize available metadata regarding the rows and columns of the matrix (e.g. ordinal structure). OASIS however can incorporate this knowledge in its construction of ***f*** and ***c*** (discussed in Section 12.A). Additionally, with its natural effect size measure, OASIS can be used as a coefficient of correlation between two random variables, even continuous valued ones (discussed in Section 12.B). While Hoeffding’s inequality for sums of bounded random variables provides one candidate p-value bound, alternatives can be constructed using different methods, such as empirical variants of Bernstein’s inequality (20) or ones based on Stein’s method (21), leading to alternative optimization objectives for ***f***, ***c*** (discussed in Section 12.C). OASIS’s statistic can also be extended to different nulls (e.g. volume-based (22)) by using alternative concentration inequalities (23). While OASIS currently analyzes 2 dimensional tables, its simple and theoretically tractable approach of centering, normalizing, and projecting can also be generalized to tensors (discussed in Section 12.D). As we have shown, OASIS empirically has power against a wide variety of alternatives, and can be shown in toy models to have more power than Pearson’s *X*^2^ test in some regimes (Section 15); a more precise power analysis would help practitioners know when to best use OASIS. To improve interpretability, regularization can be imposed on ***f*** in the optimization formulations to yield sparser vectors resulting in more parsimonious descriptions of deviations between samples. Finally, as currently presented, in a high dimensional setting with large numbers of tables generated in a single experiment over the same set of samples, OASIS performs testing on each table independently. We have introduced two approaches for jointly analyzing tables, data-aggregation and ***c***-aggregation, to identify latent relationships between samples. Ongoing and future work investigates other approaches, including adaptive ones, to perform this statistical inference and testing on tensors. While we would not be surprised if such tests have been previously introduced, we have not been able to locate them in the literature.

## 8. Conclusion

This paper provides a theoretical framework for and an applied extension of a new test, OASIS, that maps statistical problems involving discrete data to a statistic that admits a closed form finite-sample p-value bound. Here, we focused on practical scientific problems, applying OASIS to data in contingency tables. We develop OASIS with an emphasis towards genomics applications, a rapidly expanding field with diverse scientific applications from single cell genomic inference to viral and microbial strain detection. The field still relies heavily on classical statistical tests and parametric models. OASIS provides an alternative to these approaches that is both empirically robust and scientifically powerful. On real and simulated data, `OASIS-opt` prioritizes biologically significant signals: without a reference genome, any metadata, or specialization to the application at hand, it can classify viral variants including Omicron sub-variants and Delta in SARS-CoV-2 as well as sub-sublineages of Mycobacterium Tuberculosis. Moreover, it is robust to noise introduced during sequencing, the genomic data generation process, including in single cell genomics (24). OASIS provides a tool towards answering questions in mechanistic biology that are manual labor intensive (6) or impossible to address with current statistical approaches. In addition to its scientifically accurate output, this lineage assignment is performed in a rigorous statistical framework that promises further theoretical extensions in clustering.

In summary, OASIS is a finite-sample valid test that has many important statistical properties not enjoyed by *X*^2^ which will enable its use across many disciplines in modern data science. It is computationally simple, robust against deviations from the null in scientifically uninteresting directions, and provides a statistical method for the analyst to interpret rejections of the null hypothesis. In addition to a finite-sample p-value bound, the OASIS statistic admits an asymptotic distributional characterization under the null. Synthetic simulations corroborate the theoretical guarantees provided for OASIS, with experiments on genomic data showing a glimpse of the discovery power enabled by OASIS.

## 9. Data Availability

M. Tuberculosis data is publicly available under Accession ID PRJEB41116 (19). SARS-CoV-2 data is publicly available under Accession ID PRJNA817806 (6). Both datasets were chosen arbitrarily from the sequence read archive (25) and were analyzed with the same method with the same default parameters. Code for SPLASH (previously called NOMAD) is publicly available at https://github.com/salzman-lab/splash, along with code for OASIS inference. An optimized implementation of SPLASH called SPLASH 2 has been developed (26) using the bounds and techniques described in this work, and is available here https://github.com/refresh-bio/R-SPLASH. Tubercuolisis data was generated with this improved implementation of SPLASH.

## 10. Proofs

As discussed, OASIS’s test statistic *S* has a simple linear algebraic characterization. For *X* ∈ ℕ^*I×J*^ the expected matrix *E* can be written as the rank 1 outer product between the row and column sums (dividing by *M*). The centered matrix is then right normalized to account for the unequal sampling depth, dividing by the square root of the column sums to appropriately normalize, yielding the centered and right normalized matrix 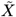. Multiplying from the left by ***f*** yields a deviation statistic for each column, which is aggregated in a ***c***-weighted sum to yield the final test statistic *S*.

To analyze OASIS’s test statistic, we define the random variables {*Z*_*j,k*_} for *j* = 1, …, *J* and *k* = 1, … *n*_*j*_. *Z*_*j,k*_ denotes the row identity of the *k*-th observation from the *j*-th column, so *Z*_*j,k*_ ∈ [*I*]. The table can then be equivalently constructed from the {*Z*_*j,k*_} by taking 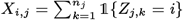. Under the null hypothesis, {*Z*_*j,k*_} are all independently drawn from ***p*** over [*I*]. This ordering *k* = 1, …, *n*_*j*_ represents a random ordering of the counts observed and is for analysis purposes only. This reformulation allows OASIS’s test statistic to be expressed as a weighted sum of independent random variables, enabling the use of classical concentration inequalities.

### A. Proof of original p-value bound

The original p-value bound proposed in (1) is:

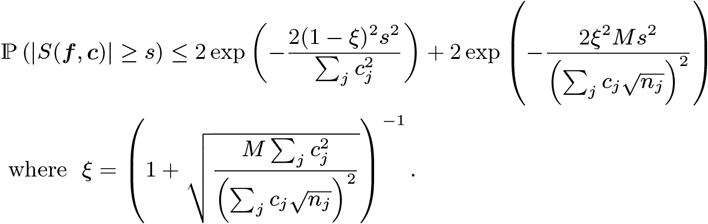

Using the quantity *γ* defined in this paper, this can be expressed as:

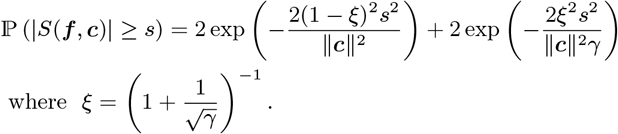

The p-value bound in (1) can be proved in the following manner. First, we estimate the expectation (unconditional on sample identity) of ***f*** on the observations as 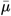. Then, we estimate the expectation of ***f*** for each column *j* separately, as 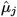. Then, we construct an estimate for the deviation of a column *j* from the table average by constructing 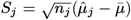, normalizing by 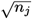 to ensure that each *S*_*j*_ will have essentially constant variance (up to the correlation between 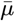 and 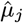). Finally, we construct our overall test statistic *S* as the ***c***-weighted sum of the *S*_*j*_, i.e. *S* =∑_*j*_ *c*_*j*_ *S*_*j*_. In summary:

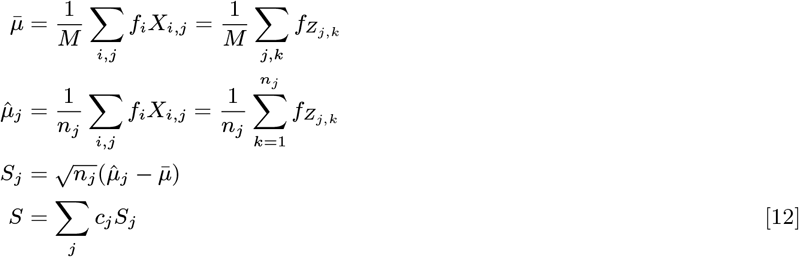

To provide a p-value bound, we use Hoeffding’s inequality:

#### Lemma 2

(Hoeffding’s inequality, Prop. 2.1 (27)). *Suppose that the random variables X*_*i*_, *i* = 1, …, *n are independent, and X*_*i*_ *has mean μ*_*i*_ *and X*_*i*_ ∈ [*a*_*i*_, *b*_*i*_]. *Then for all t* ≥ 0 *we have*

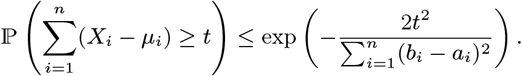

Taking the true (unknown) distribution of the {*Z*_*j,k*_} under the null to be ***p***, we define *μ* ≜ 𝔼_*Z∼****p***_[*f*_*Z*_]. Then, as in (1),

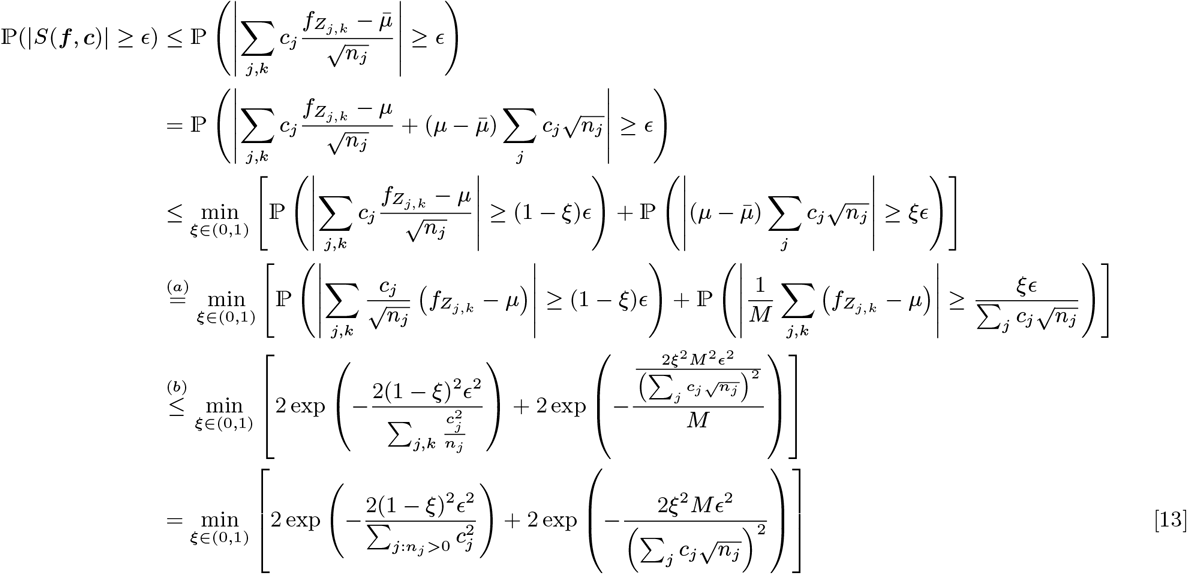

The first inequality comes from a union bound (if neither of the two conditions are met, then |*S*| *< ϵ*. (a) assumes that the denominator 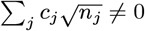, as otherwise *ξ* can simply be taken to be 0. In (b) we utilize Hoeffding’s inequality on the two terms; in the first we note that the *j, k*-th term, 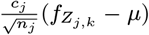 is indeed 0 mean, and is bounded as *μ* ∈ [0, 1] is simply a constant offset that shifts both the min and max by the same amount (retaining the dynamic range of 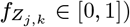. Concretely,

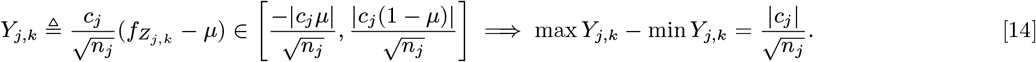

Similarly, for the second term, each summand has a range of 1. This bound can easily be optimized over *ξ* ∈(0, 1) to within a factor of 2 of optimum by equating the two terms, which is achieved when:

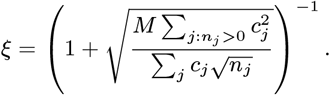

This leads to the stated p-value bound in (1) of

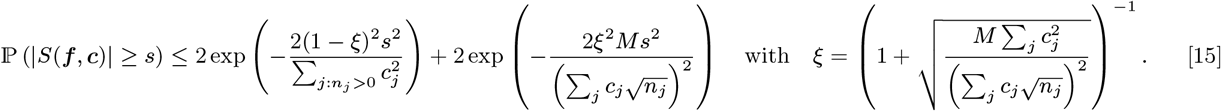

### B. Proof of improved p-values

As before, we define 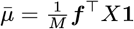 as the estimate of *μ* = 𝔼_*Z∼****p***_ [*f*_*Z*_], where ***p*** is the unknown common row distribution under the null. Due to the structure of the test statistic, the p-value bound in Equation (15) can be improved and simplified. Fixing ***f***, ***c***, the test statistic can be simplified as

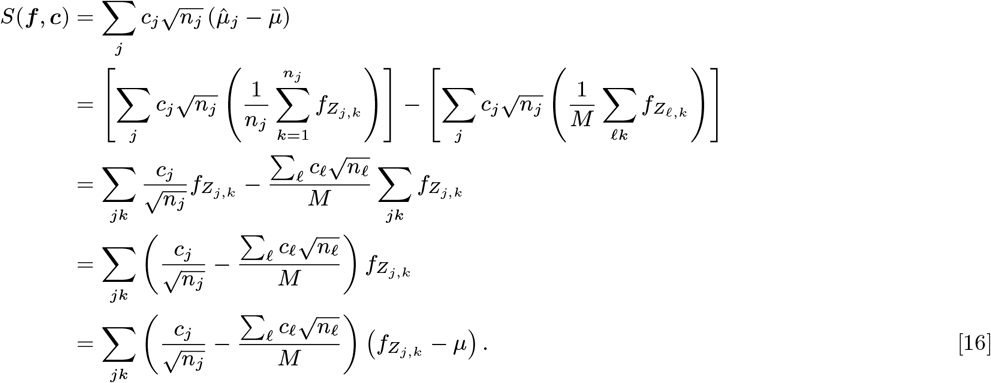

Note that, ignoring 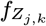, this summation is identically 0, to allow for the test statistic to have mean 0 independent of ***f*** and ***p***. This allows us to subtract the mean *μ* ≜ 𝔼_*Z∼****p***_[*f*_*Z*_] inside the summation in the last line, yielding a sum of mean 0 terms.

Note that *γ* = 1 iff 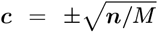 by Cauchy-Schwarz, in which case each coefficient will be equal to 0, i.e.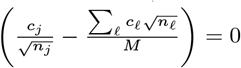 for all *j, k*. Then *S*(*f, c*) = 0 with probability 1.

If *γ <* 1, then examining each term in the summation in Eq. (16) yields

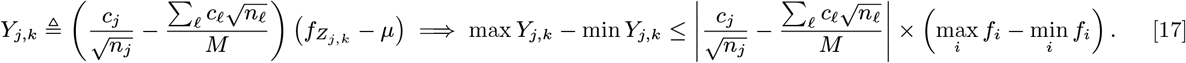

At right we are simply claiming that *Y*_*j,k*_ is almost surely bounded with the provided range. Thus, observing that each *Y*_*j,k*_ has mean 0, and is bounded as above, we can apply Hoeffding’s inequality to the sum of these *Y*_*j,k*_. For the simplification below we consider the case where max_*i*_ *f*_*i*_ − min_*i*_ *f*_*i*_ = 1, but this is without loss of generality as dividing by max_*i*_ *f*_*i*_ − min_*i*_ *f*_*i*_ simply rescales *ϵ*. Note that this condition of *γ <* 1 ensures that the denominator is nonzero.

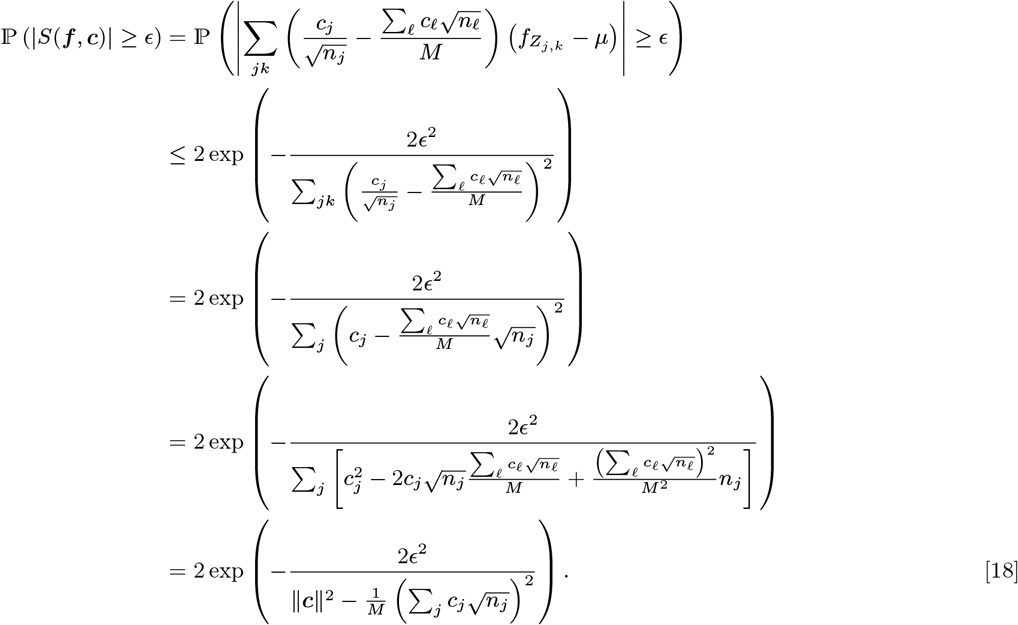

The only inequality used is Hoeffding’s; the rest is simplification and manipulation.

This improves upon the previous result, due to the smaller denominator. It also follows much more similarly to the asymptotic normality argument, where the variance of *S* appears in the denominator, up to the variance of 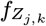. Essentially, this finite-sample valid bound takes the same form as the asymptotic bound, but upper bounds the variance of 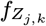 to be at most 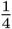, as we discuss later. Thus, our final p-value bound is

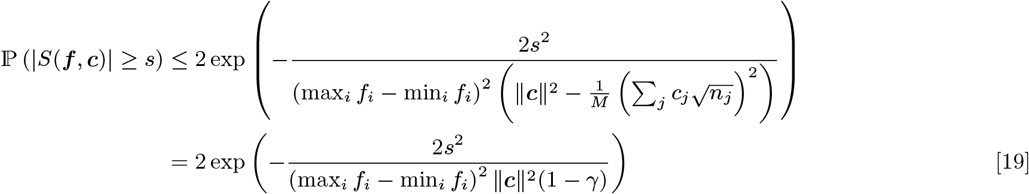

We state this general proposition below, noting that Proposition 1 follows as an immediate corollary.

#### Proposition 3.

*For any fixed* ***f*** ∈ ℝ^*I*^ *and* ***c*** ∈ ℝ^*J*^, *if γ* = 0 *then S* = 0 *with probability 1. If γ <* 1, *then*

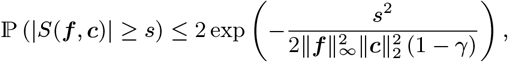

*where* ∥***f*** ∥_*∞*_ *can be tightened to* (max_*i*_ *f*_*i*_ − min_*i*_ *f*_*i*_)*/*2.

### C. Asymptotic distribution of OASIS test-statistic

Here, we show that OASIS’s test statistic is asymptotically normally distributed by using the Lyapnuov Central Limit Theorem.

#### Theorem 1

(Theorem 27.3 (28)). *Suppose that for each M the sequence X*_*M* 1_, …, *X*_*MM*_ *is independent and satisfies*

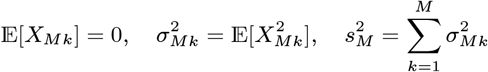

*for all M, k, where the means and variances are assumed to be finite, and* 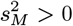 *for large M. Defining* 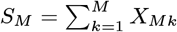,*if*

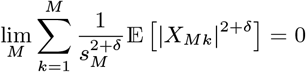

*holds for some positive δ (Lyapunov’s condition), the* 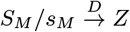where *Z* ∼ 𝒩(0, 1).

*Proof*. Continuing from Eq. (16), the test-statistic *S* can be expressed as

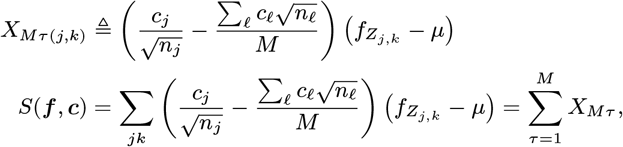

for an appropriate reindexing *τ* which maps *j, k* indices to [*M*]. Since *f*_*i*_ ∈[0, 1] for all *i*, each *X*_*Mτ*_ has finite variance, and has mean 0 due to the centering.

Computing the variance of *S*_*M*_, which is 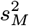 as the terms in *S*_*M*_ have mean 0 and are independent, yields

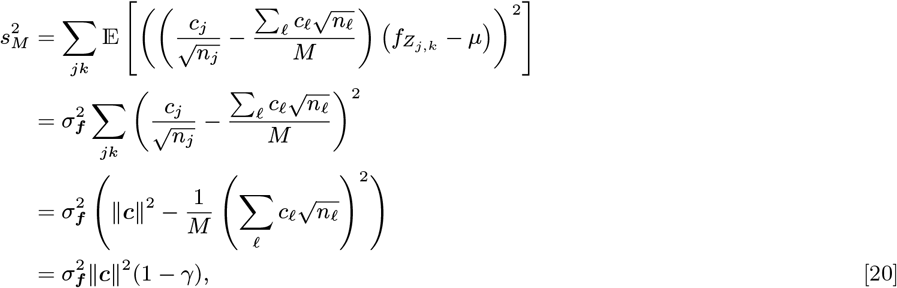

where we use the definition of 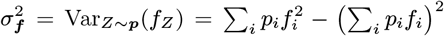. This is greater than 0 by assumption in the Proposition statement, as *γ* is bounded away from 1. Thus, our desired result is

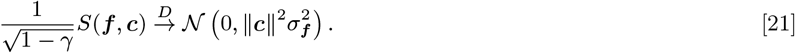

#### Satisfying Lyapunov’s condition

we show that this condition holds for *δ* = 1, which entails bounding third moments. We assume WLOG that 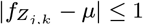 a.s. (***f*** is fixed), and so 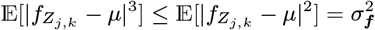. By Cauchy-Schwarz, even the largest coefficient is vanishing:

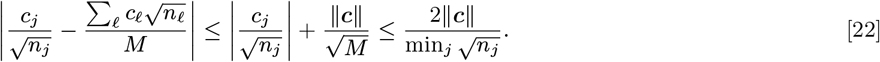

Using this, we can upper bound the sum of the third moments:

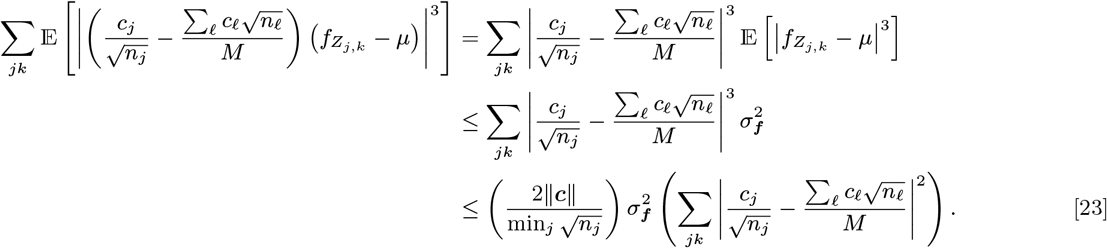

Verifying Lyapunov’s condition:

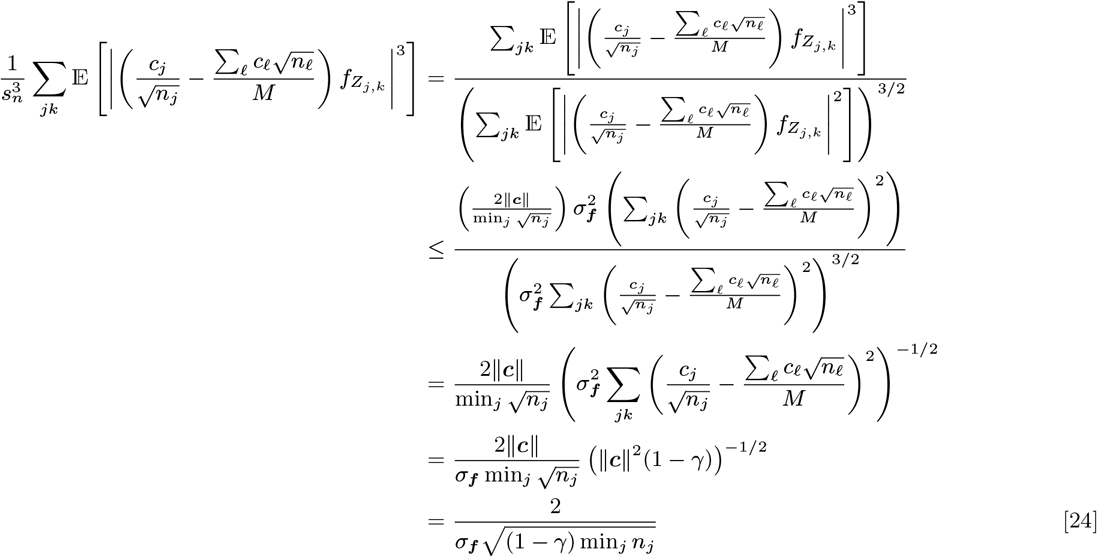

This last line goes to 0 as all *n*_*j*_→ ∞ simultaneously as long as *σ*_***f***_ *>* 0 and *γ <* 1, which were assumed in the Theorem statement, and thus by the Lyapunov CLT we have the desired result. The variance of *S*(***f***, ***c***) has already been computed as 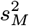, and so

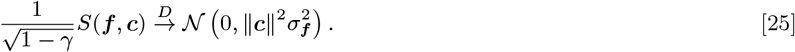

Note that if *σ*_***f***_ = 0 then *S* = 0 w.p. 1, which matches this result under the convention that a Gaussian with variance 0 is a Dirac point mass at 0. □

With this proposition, we can construct an asymptotically valid p-value. For comparison with our finite-sample p-value bound, we construct an asymptotically valid p-value bound using standard Gaussian tail bounds as

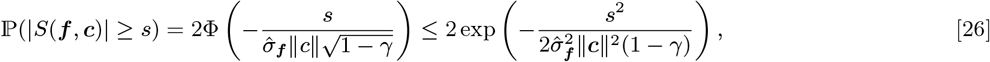

where Φ is the CDF of a standard Gaussian random variable. This holds by Slutsky’s theorem, as 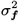 is an asymptotically consistent estimator of *σ*_***f***_ *>* 0. Note that this form precisely matches that of the finite-sample p-value bound in Proposition 1, up to bounding 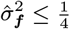 (bounding the variance of a [0, 1] random variable as 1*/*4). Since 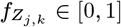, the rate of convergence can be quantified with the Berry-Esseen inequality (29) if desired.

### D. Effect Size

The effect size measure for OASIS is motivated by a simple two group alternative, where each sample originate from either group *A* or group *B*. Each group has a characteristic target distribution, ***p***_*A*_ and ***p***_*B*_ respectively, where observations from a sample in group *A* (resp. *B*) are drawn i.i.d. from ***p***_*A*_ (resp. ***p***_*B*_). Then, denoting *Z* as the identity of the random row drawn from or, the effect size estimate 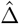 is nothing but the plug-in estimate of 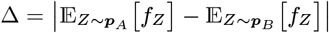 the difference between 𝔼[*f*_*Z*_] where *Z∼****p***_*A*_ or *Z∼****p***_*B*_. This is an intuitive measure of discrepancy, as OASIS declaring a discovery based on a specific ***f*** and ***c*** indicates that when partitioned into groups by ***c***, the two groups of samples have significantly different row distributions as measured by ***f***. This measure Δ is nonnegative, and is bounded above by the total variation distance between the empirical row distributions of columns where *c >* 0 and where *c <* 0. This is attained by taking the supremum over all ***f*** ∈ [0, 1]^*I*^, yielding a value of 0 if and only if the empirical row distributions are the same between the two clusters, and 1 iff they are disjoint. Note that due to the rescaling of each point’s contribution by 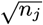(downweighting) in OASIS’s test statistic, constructing ***f*** to minimize the p-value bound will not necessarily yield an effect size equal to the total variation distance between the two empirical distributions given by the sign of ***c***.

*Proof of Effect Size Bound*. The lower bound on 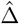 follows trivially from the absolute value. The upper bound follows from the reformulation of 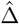 as:

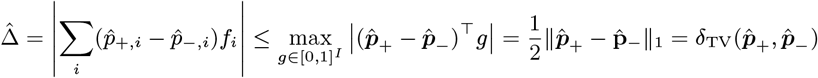

We can rewrite this as an *ℓ*_1_ norm as since 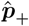 and 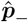 are probability distributions, shifting *g* by a constant does not change the objective value. Thus, we can equivalently maximize over *g* ∈ [−1*/*2, 1*/*2]^*I*^. Then, for any vector *x*,

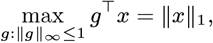

by the dual norm characterization of the *ℓ*_1_ norm.

#### D.1. Beyond binary c

There is no clear extension of this effect size measure beyond binary ***c***. We discuss some candidate measures and their drawbacks below. For simplicity, in this section we consider ***f***∈ [−1, 1]^*I*^ instead of ***f*** [0, 1]^*I*^, to avoid centering. One candidate measure which encourages binary ***f*** is:

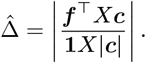

A more gradual and natural alternative which allows for non-extremized ***f*** is:

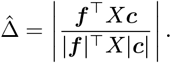

These quantities are both bounded between 0 and 1. The first attains a value of 1 only if *f*_*i*_*X*_*i,j*_ *c*_*j*_ = *X*_*i,j*_ |*c*_*j*_ |(extending the range of ***f*** to [− 1, 1]). The second requires that *f*_*i*_*X*_*i,j*_ *c*_*j*_ = |*fi*| *X*_*i,j*_ *c*_*j*_ |. However, note that these can both be trivially achieved by taking ***f*** and ***c*** to both be all ones vectors. This will not yield a significant p-value, however, due to the structure of 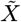.

### E. Statistical validity of data-splitting

The p-value bound outputted by the data-splitting optimization procedure is statistically valid, by a classical data-splitting argument (e.g. (13)), which we detail here for completeness. We begin by defining the random table generated by the null for probability vector ***p*** with column counts 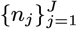. The data generated by the null is 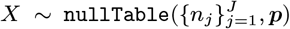. Data-splitting entails selecting a set of 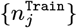 as a function of the 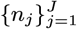 (non-random, these are given by the null). Then, due to the data-splitting procedure, 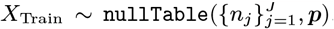,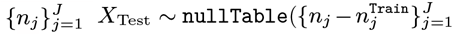, ***p***) and the two are independent. This is due to the fact that each count in the multinomial and *X*^(*j*)^ is an i.i.d. draw from ***p***, and so splitting the counts randomly between 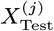 and 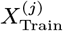 is simply separating these i.i.d. draws into two groups, resulting in a multinomial for each.

As we have shown, for any fixed ***f***, ***c***, the random variable given by the p-value bound *p*(*X*_Test_, ***c, f***) (where only *X*_Test_ is random) is a statistically valid p-value bound. We show that this choice of p-value bound, *p*(*X*) = *p*(*X*_Test_, ***c***(*X*_train_), ***f*** (*X*_train_)) is a valid p-value bound.

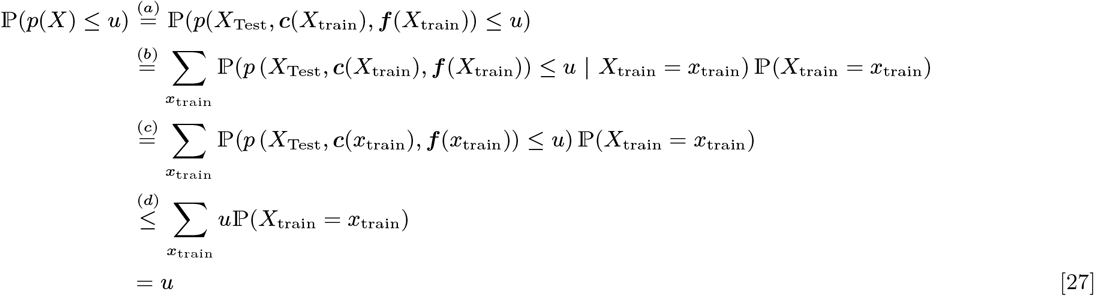

where (a) holds by definition, (b) by law of total probability, (c) by the fact that *X*_Test_ is independent of *X*_Train_, and (d) as once we condition on *X*_Train_, ***c*** and ***f*** are no longer random, and so the p-value bound from the theorem statement can be used. Finally, the last line follows from summing over the probabilities. The resulting p-value bound is not just random with respect to the data *X* (which is to be expected), but also with respect to the random splitting procedure. However, since the p-value bound holds for any fixed ***f***, ***c***, once we condition on *X*_Train_, the p-value bound can be applied.

Note that more generally, there are two sources of randomness used in the algorithm; one from splitting the data into train and test sets, and one from solving the non-convex optimization problem. The first source (splitting into train and test sets) is fundamental, whereas the second is simply for computational efficiency; if computational complexity is not an issue, we could enumerate over all possible ***f*** ∈ {0, 1}^*I*^ and exactly solve the inner optimization problem deterministically. However, the randomness in data-splitting is necessary, and can greatly impact performance. If the training data is not representative of the test data, then no matter how well the optimization problem is solved on the training data, it need not yield a significant p-value bound on the test data. Thus, a natural consideration is to generate multiple random splits of the data, and multiple hypothesis correct over these. To this end, OASIS generates independent random splits of the data by using the Poisson nature of the counts under the null. The Poisson nature of these counts is critically important, as under negative binomial overdispersion the counts observed are no longer independent, and so more sophisticated methods are needed, which do not yield fully independent sets of counts unless the overdispersion parameter is known (30).

## 11. Optimization primitive

Recall the p-value bound derived in Proposition 1 for the test statistic 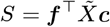, restated below for convenience, where we constrain 0 ≤ ***f*** ≤ 1:

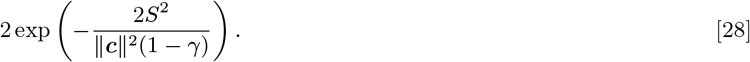

For simplicity, we shift and rescale ***f*** to be bounded as |***f***|≤ 1, which does not change the sstructure of the problem (an optimal solution for the original problem can be obtained by modifying the solution to the shifted and rescaled problem).

### A. Reformulating the optimization problem (proof of Lemma 1)

The optimization problem for minimizing the p-value bound is equivalently:

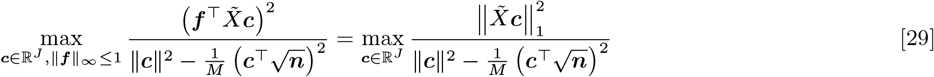

by dual norm characterization, where we recall that

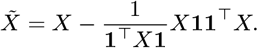

We begin by decomposing ***c*** into the part parallel to 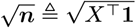 and the part orthogonal to 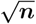, i.e. 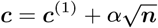 where 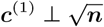 and *α* ∈ ℝ. This yields an objective value of

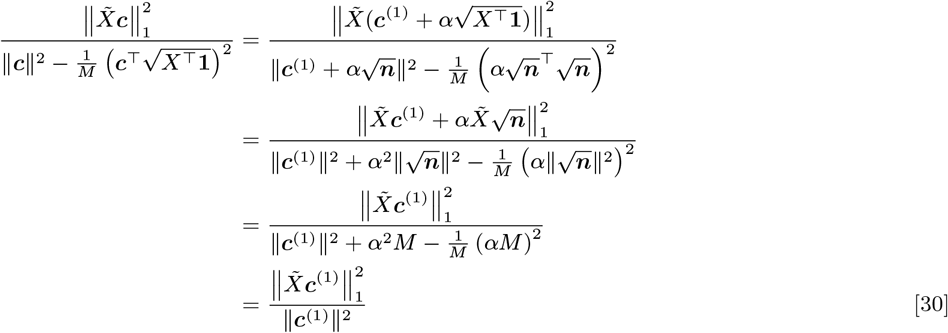

As

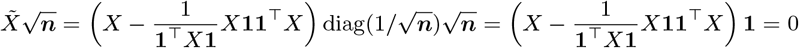

Thus, any component of ***c*** in the direction of 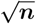 does not contribute to the numerator, and cancels out in the denominator, and so an optimal solution can be obtained by simply optimizing over ***c*** orthogonal to 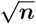. Since the denominator is equal to ∥***c***∥^2^, and the numerator and denominator both scale quadratically in ∥***c***∥, we can simply optimize the numerator subject to ∥***c***∥ ≤ 1 to identify a pair of optimal ***f***, ***c***, as:

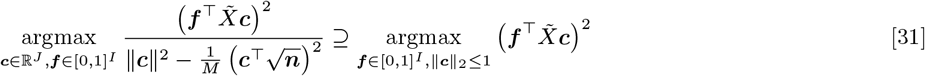

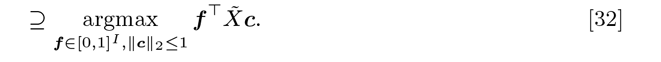

where the last line follows as ∥***c***∥ ≤ 1 ⇒ ∥ − ***c***∥ ≤ 1. This proves Lemma 1.

### B. Alternating maximization for finite-sample bound

As shown in Lemma 1, in order to obtain an optimal pair of ***c, f***, we simply need to optimize Eq. (32). We begin by enlarging the constraint set of ***f*** to be [−1, 1]^*I*^ for symmetry (an optimal ***f*** for the original problem can be obtained from such a 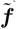 by considering 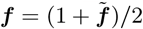 and 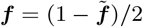, optimizing 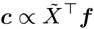, and comparing their objective values). For fixed ***c, f*** is then optimized as 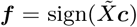, where sign(*x*) is +1 if *x >* 0, 0 if *x* = 0, and −1 if *x <* 0. ***c*** is always optimized as 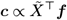. Rolling these updates together, we have that since the sign of an entry doesn’t change based on scaling, we can write our iterates {***f*** ^(*t*)^} as

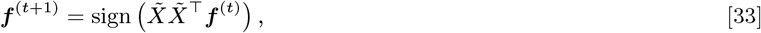

where the iterates {***c***^(*t*)^} are implicitly computed as 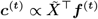.

### C. Iterative testing of contingency tables

Recalling the initial intuition for the OASIS test, where there are two groups of samples each with a characteristic probability distribution over the rows, the optimal ***c*** vector in this case partitions the samples by type (positive on one group of samples and negative on the other). However, these scalar-valued vectors ***c*** are limited by their one dimensionality, and so a natural question is whether OASIS can identify more than 2 clusters within a table. In genomics contexts this translates to detecting subclusters or multiple cell-types.

Considering the matrix viewpoint of OASIS, the test statistic can be thought of as determining whether the centered and right normalized contingency table has a single sufficiently large direction of deviation. We show that a simple extension of OASIS allows for detection of how many ways the contingency table significantly deviates from the null, through a statistical stopping condition.

Consider that one initial partitioning vector ***c***^(1)^ is identified, for example distinguishing Delta vs Omicron samples in the running SARS-CoV-2 example, and we want to determine if there are additional statistically significant ways of partitioning this data not along this same direction ***c***^(1)^. One way of accomplishing this is by attempting to identify new vectors ***c, f*** which yield a significant p-value bound, requiring that this new vector ***c*** be orthogonal to the previously found vector ***c***^(1)^. With COVID, this could be a vector separating Omicron BA.1 and BA.2. The constraint ***c*** ⊥***c***^(1)^ maintains the convexity of the optimization problem.

This naturally generalizes to multiple orthogonality constraints where ***c*** is restricted to be orthogonal to a subspace. As discussed, OASIS’s initial test statistic can be thought of as identifying one direction of substantial signal. Restricting this new ***c*** to be orthogonal to the first one is, analogous to identifying the second largest singular value and its corresponding right singular vector in the case of the SVD. Thus, this iterative procedure of finding ***c*** orthogonal to all previously found ***c*** and terminating when this no longer yields a significant p-value bound, parallels analyzing the spectrum of 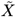 until a significant dropoff occurs.

In this iterative setting, the optimizing ***c*** is constructed as

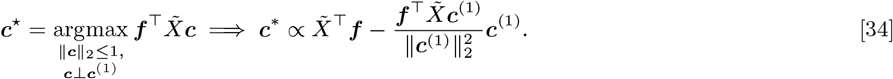

***f*** is constructed as before.

#### Algorithm 3 OASIS-iter

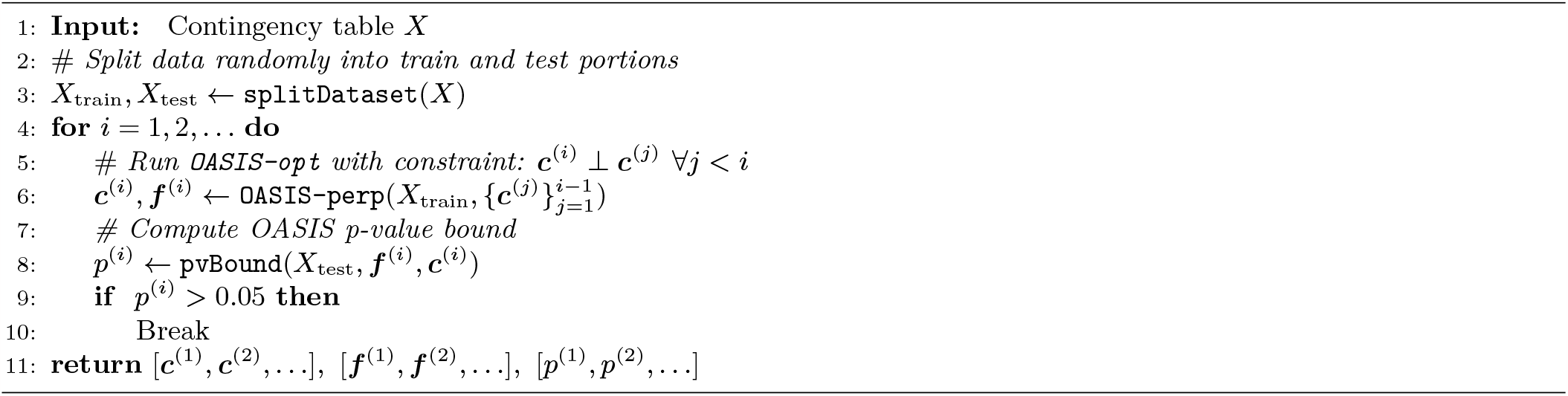

### D. Asymptotic p-value

The asymptotic p-value provided in Corollary 2.1 leads to the following maximization objective

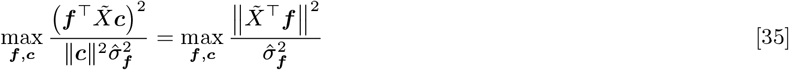

where 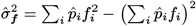 and 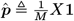.

#### Proposition 4.

*For fixed* ***c***, *an optimal* ***f*** *for* Eq. (35) *can be constructed as*

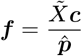

*and* 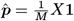.

We show this by taking the partial derivatives with respect to *f*_*i*_, noting that the optimizing ***f*** is shift invariant, and that the objective value does not change with rescaling of ***f***, as we detail below.

*Proof*. First, we show that the objective is shift invariant with respect to ***f***. It is clear that variance is shift invariant, and

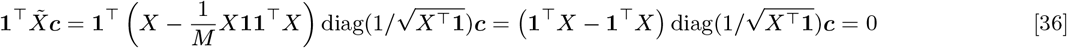

Considering the objective for the asymptotic p-value:

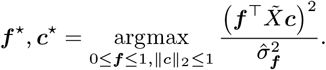

Since the numerator and denominator both scale quadratically in the magnitude of ***f***, we can instead treat the denominator as a constraint. This entails constraining the denominator 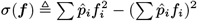 to be at most 1, and maximizing the numerator subject to *σ*(***f***) ≤ 1. Since *σ*(***f***) is a convex function of ***f***, we can simply find a stationary point with respect to ***f*** and this will be the optimum.

Taking the partial derivative with respect to *f*_*i*_ and equating to 0, we obtain (defining 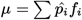)

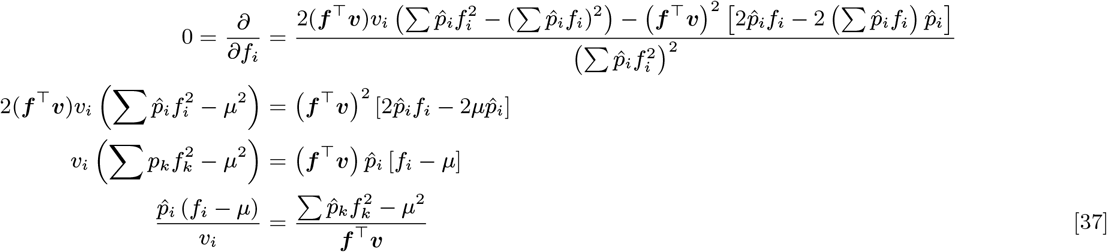

We can write out a system of *I* + 1 equations in *μ* and the *I* unknown *f*_*i*_’s, and solve by setting *μ* = 0 (shift invariant, can be arbitrary), and so obtain that 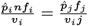 for all *i, j*. This yields 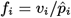 with

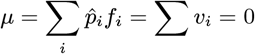

as shown in Eq. (36).

Thus, since 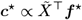, and 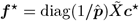, any optimal ***f*** must satisfy

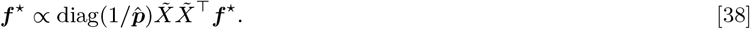

Thus, an optimal ***f*** can be obtained by taking the right eigenvector of 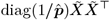 with the largest eigenvalue.

#### D.1. Modification for finite-sample bound

The ***f*** optimizing the asymptotic p-value, derived in the previous section, can be modified to better suit the finite-sample valid p-value bound. This is accomplished by binarizing the vector as 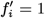 if *f* ≥0, and 0 otherwise. Then, for this ***f*** ^*′*^, the optimal ***c*** can be identified as 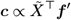.

Introducing randomness, the identified ***f*** from the asymptotic optimization procedure can be randomly rounded to 0, 1^*I*^ with probabilities proportional to the corresponding entry of ***f*** (after normalizing this vector to [0, 1]^*I*^), allowing for multiple trials and a better final objective value.

### E. More computationally intensive optimization methods

Analyzing the reformulated finite-sample optimization problem, observe that the form of the finite-sample p-value bound objective is simply a quadratic program:

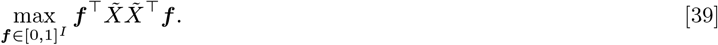

Since 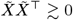 is positive semi-definite, this is an integer constrained quadratic program. This is a known combinatorial optimization problem, and certain algorithms for optimizing it have been devised.

One well-known approach is based on SDP relaxations. This works by relaxing the vector ***f*** into an *I*×*I* matrix optimization variable *A* and optimizing

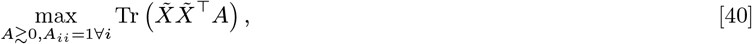

where Tr denotes the matrix Trace operator. The objective value of Eq. (40) upper bounds that of Eq. (39). We can convert the solution to Eq. (40) to an integer solution via Goemans-Williamson rounding (31), which picks a random vector ***v*** on the unit sphere in ℝ^*I*^, and assigns ***f***_*i*_ = 1 if *A*_*i*_***v*** ≥ 0 and ***f***_*i*_ = −1 otherwise. This attains an expected approximation ratio of at least 2*/π*, compared to the optimal solution of Eq. (39). Multiple random ***v*** can be selected to slightly improve the resulting solution.

In practice however, solving an SDP is computationally intensive; in the biological setting of interest, contingency tables have many rows due to sequencing error in observing the targets. This makes methods whose sample complexities scale superlinearly in the number of rows too computationally intensive to use. Future work on computationally efficient methods with provable approximation guarantees can be readily utilized in this optimization framework; the primary contribution of OASIS is in its formulation of minimizing the p-value as an optimization problem, for which we propose an alternating maximization-based solution which empirically yields good performance in an efficient manner.

#### E.1 Performance

We study the performance of these different optimization methods by simulating the “strong signal” setting described in Figure 1 and Section 14.A. The tables generated have 12 rows and 10 columns, and a clear two group structure, with signal focused in the first two rows of the tables. When run on 1000 random tables, the aforementioned SDP relaxation followed by rounding attained OPT 88% of the time. Alternating maximization yields similarly good performance, attaining OPT over 82% of the time. Computing the asymptotically optimal ***c, f*** and rounding (Section 11.D.1) attains OPT 72% of the time on this synthetic dataset. Computing ***f***, ***c*** from the SVD of 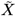 yields OPT 0% of the time, even if we postprocess ***f*** by rounding its entries

However, solving the SDP relaxation is significantly more computationally intensive and requires more complicated machinery (cvxpy (32)), while alternating maximization requires only matrix vector products, and the asymptotically optimal ***f*** requires an eigendecomposition. To test the empirical times required, we increase the counts per column from 10 to 100 to make sure that all rows have counts. We then test for various numbers of rows, to show the poor scaling of SDP solvers. Results are described in Table 4.

**Table 3.** Absolute cosine similarities between ground truth clinical metadata (whether a sample is infected with Delta or not) and the binarized sample embeddings identified by **X2** (correspondence analysis) and `OASIS-opt` (***c*** vector). For each set of calls, we display the number of anchors called, the median cosine similarity, and the 90-th quantile of cosine similarity.

| | Pearson $X^2$ | OASIS | OASIS called, $X^2$ didn’t | $X^2$ pv = 0 | OASIS pv = 0 | OASIS filtered |
| --- | --- | --- | --- | --- | --- | --- |
| Number of anchors | 71,543 | 28,430 | 389 | 16,611 | 2114 | 2495 |
| 0.5 quantile cosine similarity | 0.10 | 0.22 | 0.15 | 0.09 | 0.69 | 0.72 |
| 0.9 quantile cosine similarity | 0.52 | 0.66 | 0.76 | 0.40 | 0.79 | 0.82 |

**Table 4.** Computational time (in seconds) for computing the optimized ***c, f*** using the stated method, on 1000 matrices. As can be seen, the SDP solver quickly becomes infeasible. Forming the matrix 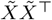 also becomes costly as the number of rows increases. Alternating maximization stays efficient and performant. Timing results generated on a MacBook Air (M2, 2022), runs declared as a failure and stopped after 5 minutes.

| number of rows ($I$) | SDP+rounding | Alternating maximization | Asymptotically-optimal<br>(eigenvector + rounding) |
| --- | --- | --- | --- |
| 12 | 14.5 | 0.86 | 0.20 |
| 24 | 194 | 1.2 | 0.25 |
| 48 | - | 1.26 | 0.59 |
| 96 | - | 1.31 | 5.9 |
| 192 | - | 1.48 | 16.7 |

**Table 5.** Power at λ ≈ 129, noise model as in (16). Exponentially distributed target distribution, described in text. Full plots in Figure 10.

| num rows | num cols | $X^2$ | OASIS-rand | OASIS-opt |
| --- | --- | --- | --- | --- |
| 5 | 10 | 0.370 | 0.005 | 0.007 |
|  | 50 | 0.765 | 0.009 | 0.006 |
|  | 400 | 0.995 | 0.001 | 0.057 |
| 10 | 10 | 0.792 | 0.011 | 0.015 |
|  | 50 | 1.000 | 0.007 | 0.046 |
|  | 400 | 1.000 | 0.010 | 0.309 |
| 20 | 10 | 0.999 | 0.065 | 0.080 |
|  | 50 | 1.000 | 0.058 | 0.213 |
|  | 400 | 1.000 | 0.100 | 0.866 |
| 100 | 10 | 1.000 | 0.871 | 0.978 |
|  | 50 | 1.000 | 0.983 | 1.000 |
|  | 400 | 1.000 | 0.994 | 1.000 |

The asymptotic approach identifies the principle right eigenvector of 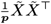 where 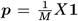, which naively requires forming the *I* × *I* matrix. However, using power iteration, only matrix vector products with 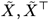 are required, which can be much more efficient.

### F. Comparison with SVD

The SVD is frequently used for matrix decomposition and interpretation tasks, but doesn’t inherently minimize the p-value bound of Proposition 1 or yield desirable partitionings. Mathematically, the SVD yields ***f***, ***c*** which maximize

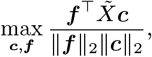

as opposed to the p-value bound objective of

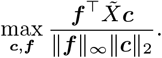

An optimal ***f*** exists at a corner point, and that if entries of 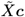 are not equal to 0 then this optimizing ***f*** is unique up to the absolute value (***f*** and 1 −***f*** yielding the same objective value). For any ***f*** with entries not in 0, 1, at least as good of an objective value can be attained by extremizing ***f*** according to the entries of 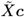. As discussed in the previous section, computing ***f***, ***c*** from the SVD of 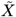 yields OPT 0% of the time, even if we postprocess ***f*** by rounding (extremizing) its entries.

Even if the SVD-based ***f*** is shifted and extremized to be in [0, 1]^*I*^, this is still not optimizing the desired objective. One simple toy example where an SVD does not yield the desired partitioning is shown in Figure 6. Figure 6 simulates a planted model, where there are two subgroups of columns, one which expresses rows 1-5, and the other which expresses rows 6-10. The values in these entries is approximately 40. However, there is a confounding final row, which randomly groups the columns into two new groups, and yields counts either 160 or 0. Multinomial sampling provides slight noise to make this more realistic. While both methods do call this table as significant (extreme example chosen for clarity of exposition), an SVD-based approach is unable to identify the latent column structure, due to its *ℓ*_2_ normalization of ***f***. The latent structure is identified by a vector ***f*** which takes value 1 on the first 5 rows, 0 on the next 5, and 1/2 (i.e., disregarding), the final row. However, an SVD restricts to an *ℓ*_2_ constraint, and since the last row has more counts, it is selected as the dominant component in the SVD-based ***f***. Taking an SVD, the magnitude of this last component in ***f*** is 0.76, where all other components have magnitude at most 0.28. In comparison, OASIS-asymp, which maximizes the asymptotic objective, gives this last row a value of less than 9 × 10^−4^ in magnitude, where all other components have a magnitude of at least 0.17. These are depicted in more detail in Figure 7.

## 12. Additional discussion

### A. Metadata-based construction of *f, c*

OASIS is stated generally, and treats the rows as the set [*I*] and the columns as the set [*J*], with no additional information regarding shared structure. However, considering the motivating biological application, rows have additional information (the associated target sequence in {*A, T, C, G}* ^27^ (1)). Considering this, one potential direction to construct more biologically meaningful inference is to ensure that similar sequences have similar *f*_*i*_ values. This can be accomplished by constructing ***f*** to be Lipschitz with respect to the Levenshtein distance between the targets of the associated rows.

Similarly, in the presence of sample metadata (e.g. cell-type information) in the form of a vector 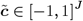, the user may wish to incorporate this information into constructing ***c***. In the simplest case, the metadata is fully trusted, and we want to see if this sample partitioning yields a significant p-value bound. In this case, 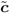 is utilized, along with the optimizing ***f*** ^⋆^ from the split data, for inference.

One additional approach of potential interest is if metadata 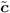 is given, but we are still free to reweight samples: not all samples are equally representative. If we require that each *c*_*j*_ retains the same sign as 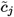, then we can formulate this as:

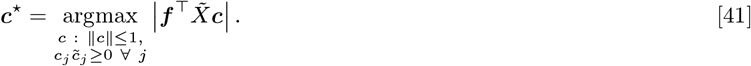

While the overall objective is non-convex, it is still biconvex. The maximization of ***f*** is the same as before, and ***c*** is now maximized as, defining 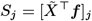 :

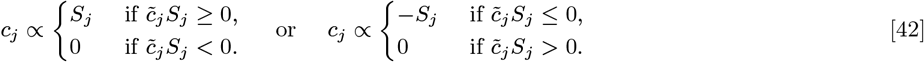

To optimize the overall truncated objective, we can still run alternating maximization.

Many other forms of metadata (e.g. catgory of cell-type beyond just binary) can yield different metadata-aided approaches, and can be framed as optimization problems that admit efficient alternating maximization solutions.

### B. OASIS as a coefficient of correlation

OASIS can also be used to test the dependence between two random variables. Consider observations of two real valued random variables (*X, Y*), where we observe draws from the joint distribution (*X*_*i*_, *Y*_*i*_) and want to test against the null where *X* and *Y* are independent. One recent measure of dependence was proposed (33). OASIS can also be used to test against this, by quantizing the random variables (e.g. into deciles), and constructing a contingency table from these categorical (quantized) observations. For any quantization, OASIS can yield a finite-sample valid p-value bound against the null of independence. This empirically performs quite well, and can yield an effect size estimate comparable to Xicor (33). With no tuning of quantization (fixed at deciles), we show OASIS’s effect size and power in Section 14.H.

### C. Bernstein-based p-value bounds

The finite-sample valid p-value bounds provided in Proposition 1 suffer from the fact that they do not incorporate the variance of *f*_*Z*_ where *Z* ∼ ***p***, and simply rely on the boundedness of *f*_*Z*_. One natural concentration inequality that utilizes both the boundedness of a random variable as well as its variance is Bernstein’s inequality, of which an empirical variant was derived in (20). An improved p-value bound can potentially be obtained by bounding the true variance 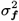 with high probability by some function of the empirical variance 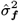 on the training set, and utilizing Bernstein’s inequality instead of Hoeffding’s, applied to the same test statistic.

### D. Tensor analysis

In computational genomics, it is of high priority to discover effects or partitionings that are reproducible. In the setting of (1) this motivates going beyond treating each table independently, and instead analyzing the anchor x sample x target tensor. OASIS’s framework enables this, with metadata-guided construction of ***f***, ***c***, to be performed jointly.

Note that more generally OASIS, with its simple linear structure, suggests a simple test for tensors. If we have a 3 dimensional tensor, we can construct and analyze a test statistic similar to OASIS by centering the tensor, appropriately normalizing by sampling depth, then taking inner products along each of the 3 dimensions to reduce to a scalar.

## 13. SARS-CoV-2 Analysis details

The SARS-CoV-2 analysis was based on dehosted sequencing data of nasopharyngeal swabs with coinfections available under accession PRJNA817806 (6).

SPLASH contingency tables were tested using `OASIS-opt` with 25% train/test split. We then utilize anchors whose BY corrected p-value bound is less than 0.05, whose effect size (when *c*^⋆^ is binarized) is in the top 10% of anchors, and have a total number of counts *M >* 1000. This yields 2495 anchors. We then iterate over these 2495 anchors, and rerun the alternating maximization procedure to generate *c*^⋆^, *f*^⋆^ on the full table.

Note that in addition to the aforementioned thresholds, there are several potential avenues towards filtering that we did not finetune, including how balanced the clustering is (ensure that the table is not simply one deviating column), how similar a ***c*** vector is to others (not detect random splits, e.g. due to SNPs), and reducing similar anchors (noting that one single base-pair mutation may yield up to *k* significant anchors that are one base-pair offsets of each other).

The alignment was performed using Bowtie (18). We used 4 publicly available COVID reference genomes for the strains circulating during this time period: the Wuhan reference genome (NCBI assembly NC_045512.2), Delta (OK091006.1), Omicron BA.1 (OX315743.1), and Omicron BA.2 (OX315675.1). A bowtie index comprised of the four COVID genomes was constructed (bowtie-build on a fasta file containing four reads, one for each assembly). We then construct a fasta file from the anchors passing the above filtering steps, and their targets which comprise *>* 5% of the total counts for that anchor. This fasta is aligned to the bowtie index using (-v 0 -a) options to ensure that we only get exact matches, and record all possible matches.

### A. c-aggregation

To aggregate the ***c***^(*a*)^ vectors obtained from experiments on different anchors *a* (different tables), one natural objective is

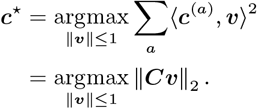

Here, we stacked the ***c***^(*a*)^ vectors to obtain a matrix ***C***. Analyzing this objective, an optimizing ***v*** is attained by the principle right singular vector of ***C***. To analyze this further, we also plot the second singular vector, which can be understood as finding a vector with maximal sum of squared inner products with ***c***^(*a*)^ subject to being orthogonal to the first ***c*** identified, and see that this separates Omicron BA.1 from BA.2.

### B. Data-aggregation

Data-aggregation iteratively finds orthogonal ***c*** vectors on the stacked contingency tables. To merge the tables, we discard all targets that have fewer than 10 counts in a given table. Then, we normalize each table by 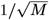, its total number of counts, to ensure that high count tables do not completely outweigh low count ones, and stack the resulting tables together. This yields a visually similar clustering to ***c***-aggregation, but does not perfectly predict with ***c***^(1)^ ≷ 0 whether the sample has Delta or not. There is still perfect classification accuracy on ***c***^(1)^ for predicting whether a sample has Delta or not, and 91% accuracy for predicting from ***c***^(1)^ and ***c***^(2)^ whether a sample has Omicron BA.1 as its primary strain or not (correctly classified 94/103).

For predicting target strain, of the 51,557 anchor-target pairs (rows of *X*_agg_), we filtered for those targets which constitute at least 5% of the total counts for that anchor. Then, we label a target as Delta if it maps to the Delta assembly and neither of the Omicron ones, and Omicron if it maps to at least one of the Omicron assemblies but not the Delta one. This yields 4836 sufficiently abundant anchor-target pairs which map, constituting 2,479 of the 2,495 anchors tested.

## 14. Additional simulations

For brevity and flow we defer some additional plots from the main text to here.

### A. Comparison of OASIS and *X*^2^ in planted setting

1. In this section we define and provide additional simulations for the example in Figure 1. We define a class of alternatives parameterized by their total number of counts *M* and a corruption parameter *ϵ* as follows (the number of rows and columns are fixed as 12 and 10 respectively). In our planted model the the probability vector for the first half (5) of the samples is the first standard basis vector ***p***^(*j*)^ = *e*_1_ for *j* = 1, …, *J/*2, and the latter half of the samples have ***p***^(*j*)^ = *e*_2_ for *j* = *J/*2 + 1, …, *J*. To vary between structured and unstructured signal, we mix each columns probability distribution with another probability distribution ***q***^(*j*)^, which is generated independently for each column. Concretely,

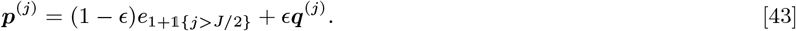

The observed matrix is generated by drawing *X*^(*j*)^ ∼ multinomial(*M/J*, ***p***^(*j*)^), independently for each of the *J* columns. Each ***q***^(*j*)^ was generated by taking each entry to be independently uniformly distributed between [0, 1], and then normalized to a valid probability distribution.

Considering in more detail the specifics of Figure 1, in subplot C) we showed two tables highlighting the shortcomings of Pearson’s X2 test. In both examples, power is hampered by the large number of rows. If the number of rows with nonzero counts doubles, and the value of the test statistic changes minimally, then the p-value X2 provides will decrease dramatically (square root of the original p-value, 10^−10^ → 10^−5^. Additionally, X2 normalizes by the square root of the row sums, downweighting abundant rows and upweighting rare rows (potentially due to sequencing errors).

This is a critical drawback, as seen in the strong signal setting: because the first two rows are abundant, X2 downweights their deviations, and yields only a moderately significant p-value. In the weak signal setting, even though no one row constitutes a large deviation, X2 upweights the deviation in each row, and sums, yielding a similarly significant p-value. In contrast, OASIS identifies the strong and reproducible deviation in the the first setting, yielding an extremely significant p-value bound by focusing on the difference in expression of the first two rows. In the weak signal setting, OASIS struggles to find a strong separation between the samples, and only yields a slightly significant p-value bound. A spectral analysis of *X*_corr_ for the two tables is shown in Figure 9, highlighting the flat spectrum of the weak signal table and spiky spectrum of the strong signal table. In Figure 1D, results are shown using 5 random splits. We additionally show results using just one random split in Figure 8.

### B. Robustness against simulated alternatives

OASIS provides robustness against many undesirable alternatives, beyond just negative binomial overdispersion. We provide additional simulation evidence, showing OASIS’s robustness against corruption of each individual columns probability distributions Figure 12.

**Fig. 10.**
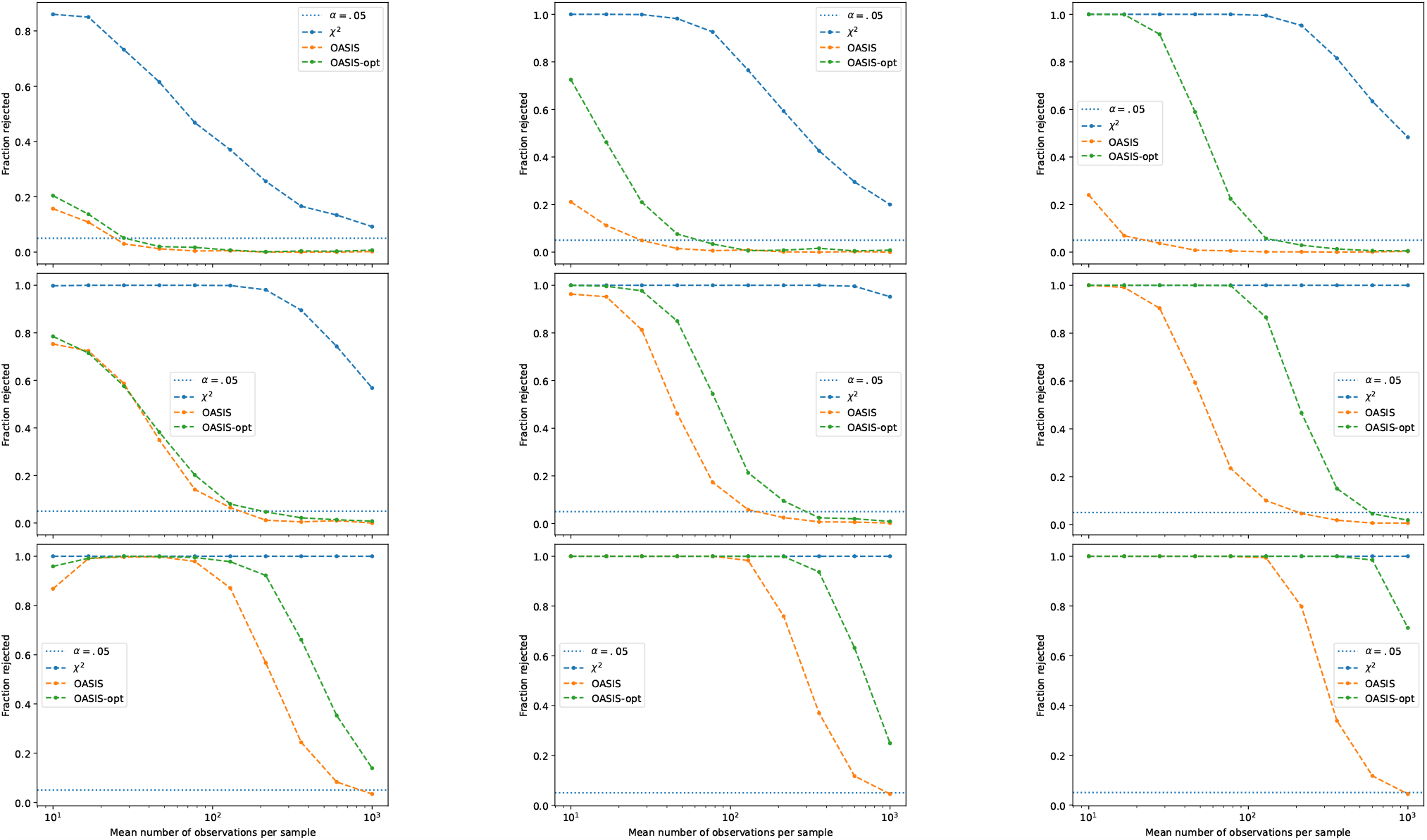
Simulated data for negative binomial overdispersion. Null target distribution generated from exponential distributions, described in Section 14.B.1. OASIS has significantly better control of the false discovery rate than the *X*^2^ test, and requires substantially fewer samples per column to control the FDR. Left to right, the columns correspond to tables having 10, 50, and 400 columns. Top to bottom, the rows indicate 5 rows, 20 rows, and 100 rows.

**Fig. 11.**
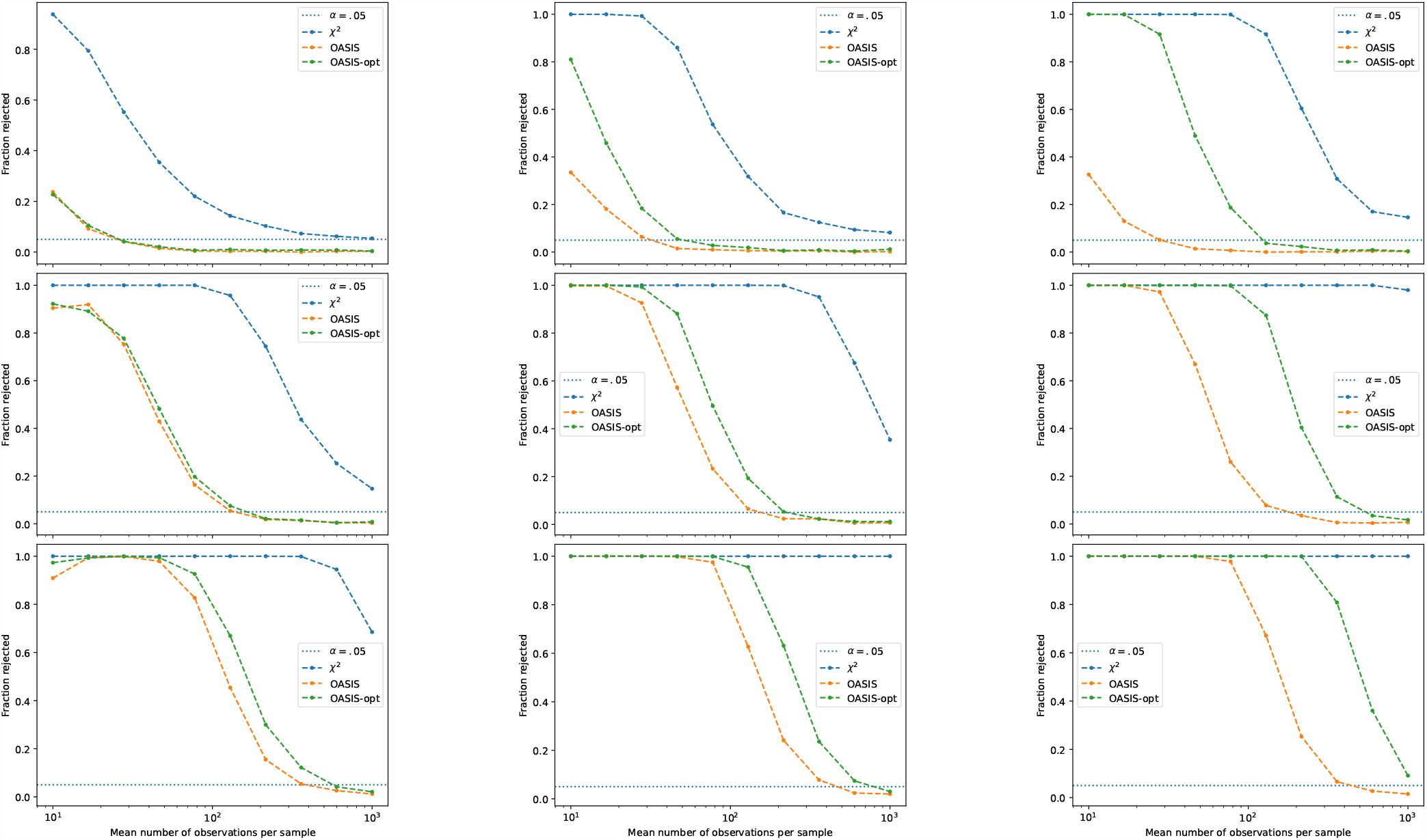
Simulated data for negative binomial overdispersion. Null target distribution is uniform over rows. OASIS has significantly better control of the false positive rate than the *X*^2^-test, and requires substantially fewer samples per column to control the FDR. Left to right, the columns correspond to tables having 10, 50, and 400 columns. Top to bottom, the rows indicate 5 rows, 20 rows, and 50 rows.

**Fig. 12.**
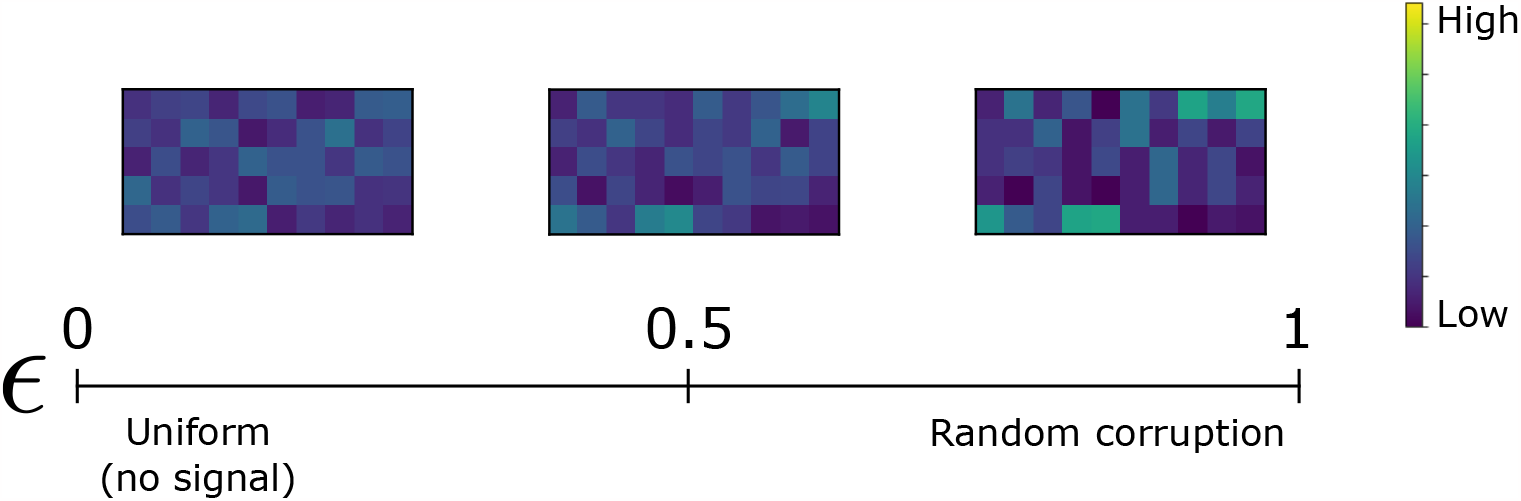
Simulated data under independent *ℓ*_1_ corruption to each columns probability vector. *X*^2^ has significant power against this alternative, even though it represents contamination and a “biological null”.

#### B.1. Negative binomial overdispersion

As discussed, OASIS is robust to negative binomial overdispersion, as modelled in (16). Under the statistical null, for a common row distribution ***p***, if the number of observations in each column is Poisson distributed with rate *λ*, then the observed counts satisfy *X*_*i,j*_ ∼ Pois(*λp*_*i*_), independently drawn for each entry. However, as shown by (16), sequencing data is overdispersed in practice, and should be modelled as negative binomial. For an overdispersion parameter *θ* (where *θ* = 0 corresponds to the true null), we thus generate synthetic data as *X*_*ij*_ ∼ Pois(Γ(*λp*_*i*_*/θ, θ*)). Following the trends depicted in (16), we model our overdispersion parameter as a function of the expected number of counts to be observed *n*, as *θ*(*n*) = 3*/n* if *x <* 150, and *θ*(*n*) = 0.02 otherwise. In these plots, the number of observations per column was varied from 10 to 1000 in log-scale through 10 equally spaced points. We focus on one which shows most clearly the full dynamic range, index 6/10 with a value of 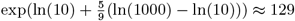.

To generate a random target distribution from the null, we simulated two settings. One parameter free method is to take a uniform target distribution. Another way, modelling the more realistic setting where some targets are more likely than others, is to generate a vector 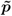 where each entry is drawn independently from some distribution that has nontrivial tails, and renormalize this to a probability distribution, 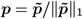. Here, we utilized an exponential distribution with rate parameter 5. In both settings, with uniform or exponential target distributions (Figures 10 and 11), OASIS provides significantly better control of the discovery rate against this uninteresting alternative stemming from overdispersion.

### B.2. Robustness again ℓ_1_ corruption

In this setting, we study an alternative where each column is uniformly distributed, but is then mixed (with weight *ϵ*) with a probability distribution ***q***^(*j*)^, which is generated independently for each column. Concretely,

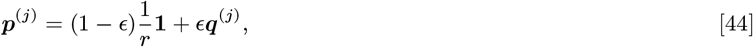

where the synthetic data is generated by taking a Poisson number of observations from each column, multinomial with that columns probability vector ***p***^(*j*)^, independently for each column. To avoid additional parameters, a uniform target distribution was used as the true null. ***q***^(*j*)^ were generated independently by taking each entry to be uniformly distributed between [0, 1] and normalizing. This is shown pictorially in Figure 12.

In simulations, we observe that OASIS is extremely robust to these types of perturbations; whereas *X*^2^ immediately identifies that these tables deviate from the null, OASIS does not prioritize these unstructured deviations, and maintains control of the discovery rate for much larger corruption magnitudes, especially for larger tables.

### C. Power against simulated alternatives

OASIS has power against alternatives with unique per-sample expression (motivated by V(D)J recombination) like those in Figure 14, and can be shown to have more power against splicing-type alternatives (Figure 16) than the *X*^2^ test in some settings.

**Fig. 13.**
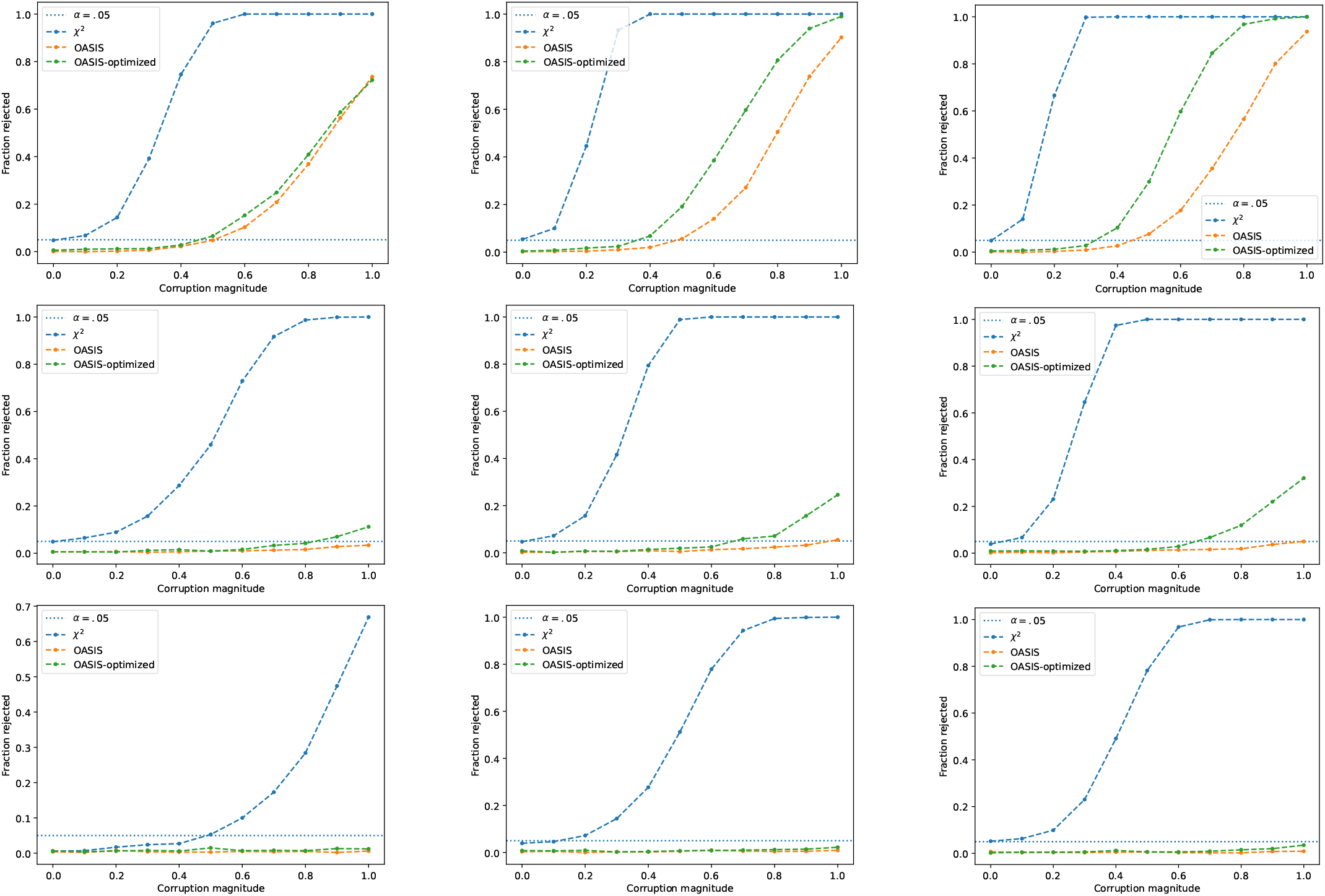
Simulated data for independent *ℓ*_1_ corruption of each column. First row corresponds to 20 rows, second to 100, and third to 500. First column corresponds to 10 columns, second to 50 columns, third to 100 columns. Number of observations in each column is Poisson distributed with mean 100, independent across columns. Observations are drawn independently, and are multinomially distributed for each column with probability vector as defined in Eq. (44).

**Fig. 14.**
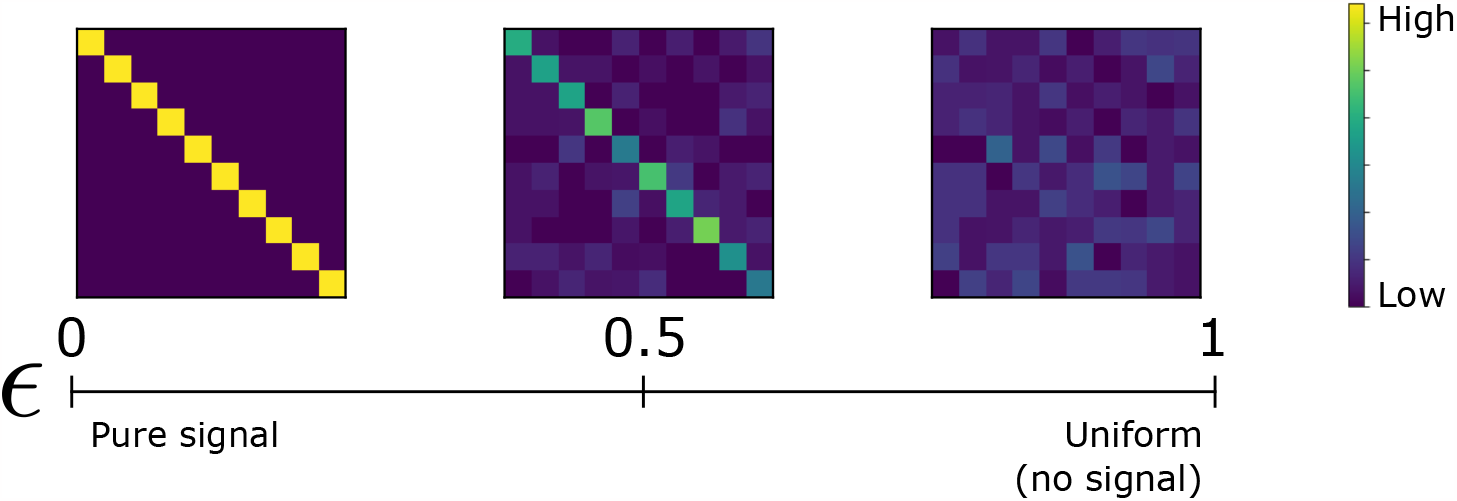
Simulated data for an alternative with unique per-sample expression, motivated by V(D)J recombination. All methods have power.

#### C.1. Unique per-sample expression

In this setting, the probability distribution of column *j* ∈ [*J*] is ***p***^(*j*)^ ∈ [0, 1]^*I*^, with *I* = *J*:

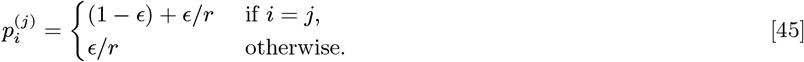

This setting is shown pictorially in Figure 14, where a corruption *ϵ* = 0 corresponds to pure signal, and *ϵ* = 1 corresponds to a uniform target distribution for all columns (no signal). We provide simulation results for this setting in Figure 15, where in most settings Pearson’s *X*^2^ has more power than OASIS, but not by a significant margin.

**Fig. 15.**
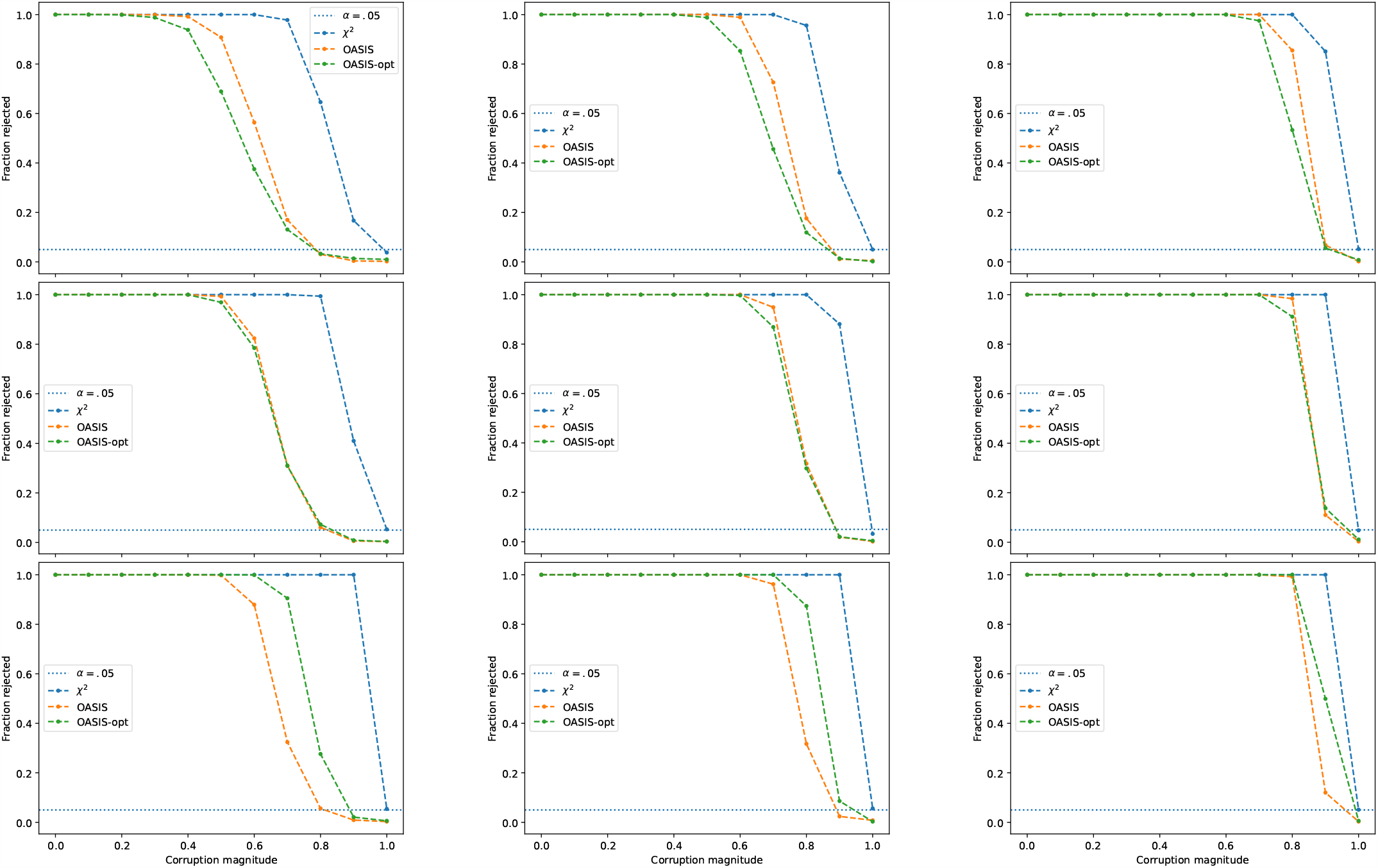
Simulated data for unique per-sample expression alternative. First row corresponds to 10 rows and columns, second to 25, and third to 100. First column corresponds to a mean of 10 observations per column, second to 20, and third to 50. Number of observations in each column is Poisson distributed with the given mean, independent across columns. Probabilistic model in Eq. (45).

**Fig. 16.**
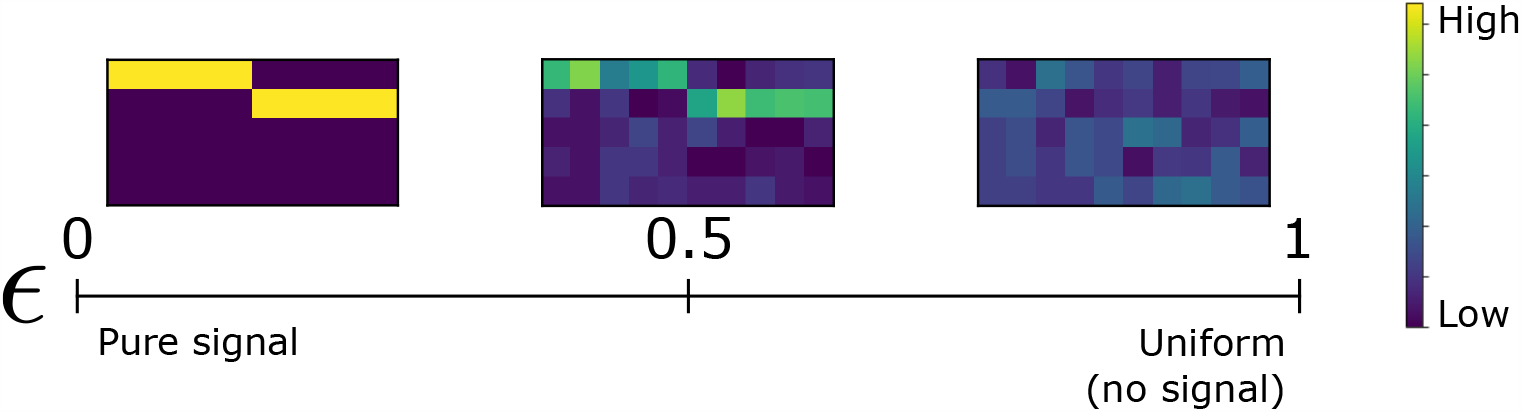
Simulated data for a differentially regulated alternative splicing-type alternative. OASIS has more power than *X*^2^.

#### C.2. Two-group setting

Here we show OASIS’s power against alternative splicing type alternatives across a broader range of parameters. Alternative tables were generated as taking two groups of samples of equal size, and defining the two group’s probability distributions as, for *k* = 1, 2

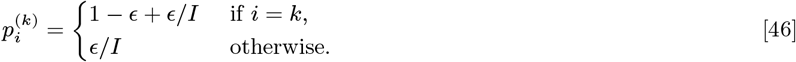

This indicates that ***p***^(1)^ has most of its counts in the first row, and ***p***^(2)^ in the second. This is shown pictorially in Figure 16.

The number of observations per sample is drawn independently from a Poisson distribution with mean 20 for each sample. Figure 17 shows that OASIS can have more power than *X*^2^ in a variety of parameter regimes. For OASIS-rand, 50 ***c*** vectors were drawn uniformly at random from 1, +1 ^*J*^ and 10 ***f*** vectors uniformly at random from 0, 1 ^*I*^, and all 500 pairs were tested.

**Fig. 17.**
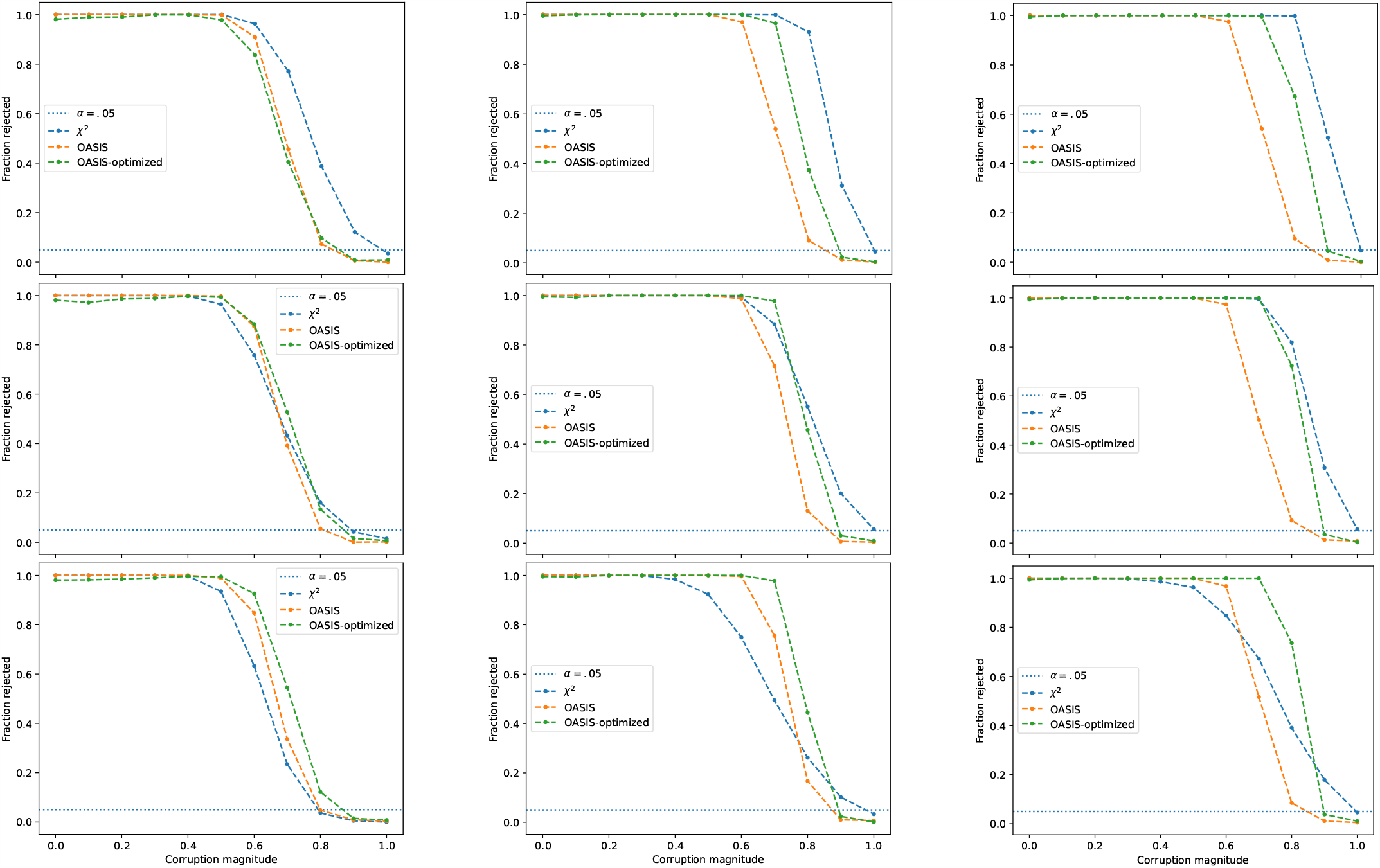
Simulated data for alternative splicing-type alternative. First row corresponds to 20 rows, second to 100 rows, third to 400 rows. First column corresponds to 10 samples, second to 50 samples, third to 100 samples. Number of observations in each column is Poisson distributed with mean 20.

### D. SARS-CoV-2 additional plots

In this section we provide additional results regarding the SARS-CoV-2 analysis provided in the main text.

We observe in Figure 21 that the improved predictive power of OASIS with respect to sample metadata is primarily driven by the tables OASIS prioritizes. When performing correspondence analysis on these tables, similar results are obtained, but when looking at all calls made by Pearson’s *X*^2^, many have worse correspondence with sample metadata.

As seen in Figure 23, iterative testing performed on the stacked data matrices yields significant p-value bounds (of value 0, up to machine precision) through 30 iterations, due to the large number of counts, and inherent latent structure. Since there are so many counts, even vectors ***c*** that do not fully align with the latent structure will yield significant p-value bounds. Thus, while for smaller matrices significance testing provides a good cutoff, for larger tables one may still need to find an “elbow” in the curve. We demonstrate this in Figure 23, where we observe that the first two p-value bounds are substantially smaller, with the rest approximately fitting a simple linearly decaying trend (when the log p-value bounds are plotted on log scale). This threshold can be programatically identified by looking at the ratio of successive log p-value bounds, and finding the maximum.

**Fig. 18.**
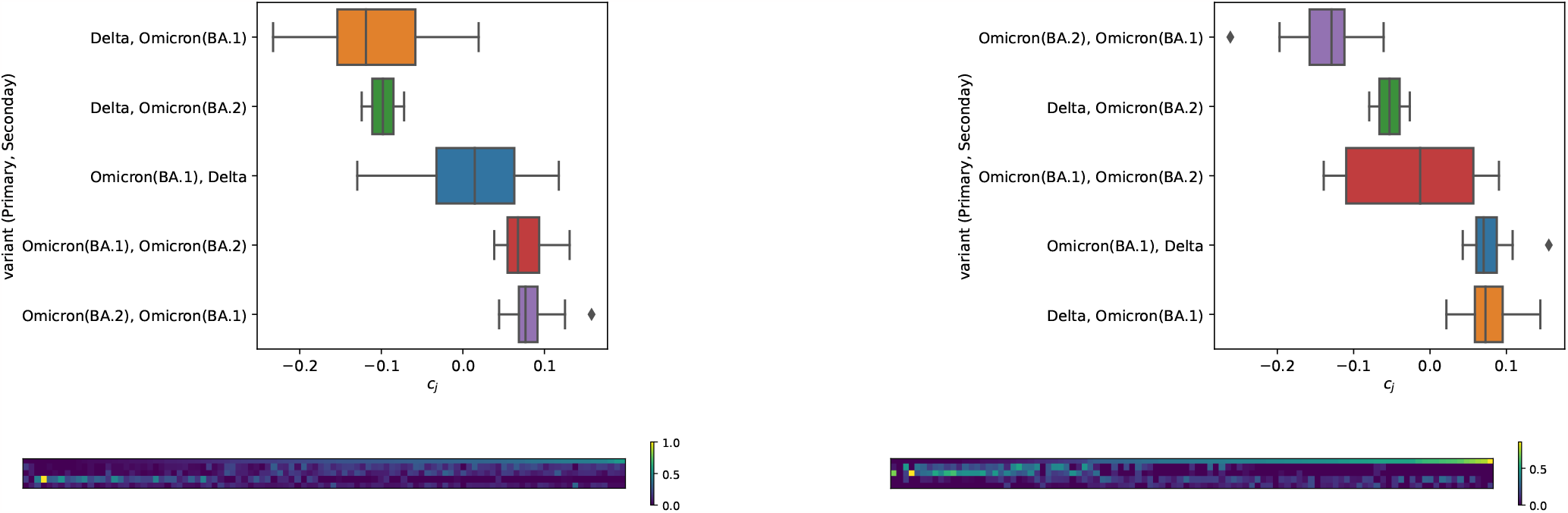
Top 2 tables in terms of `OASIS-opt` p-value bound, not called by OASIS-rand. Both tables yield a vector ***c*** with an absolute cosine similarity of 0.76 with ground truth metadata (was the patient infected with Delta or not).

**Fig. 19.**
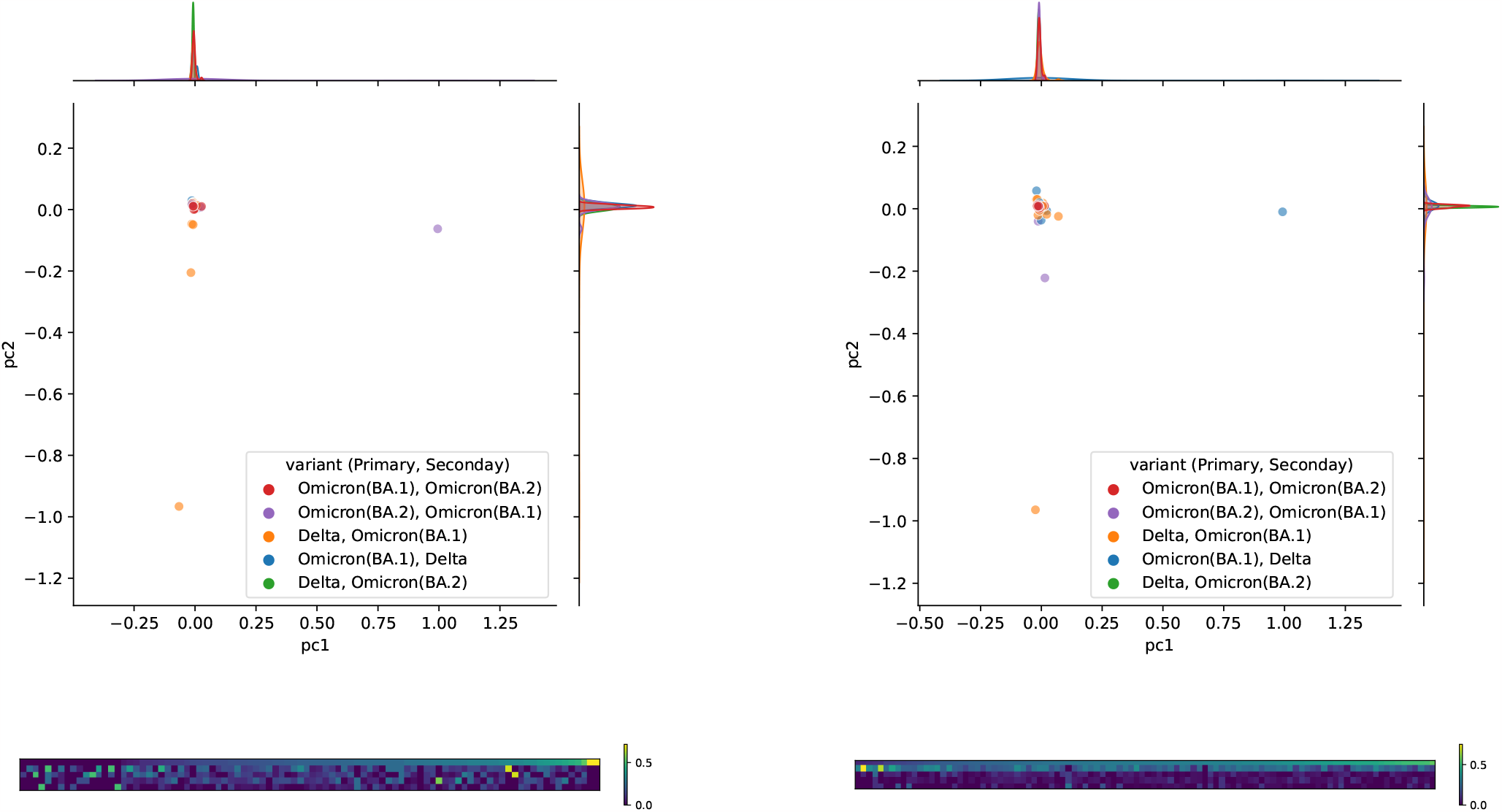
Top 2 tables in terms of *X*^2^-p-value (both with a q-value of 0), not called by `OASIS-opt` (q-value *>* 0.05). Since there were more than 2 tables with a *X*^2^ p-value of 0, we chose the two with the smallest `OASIS-opt` p-value bounds. The output 1d embedding by correspondence analysis yields an absolute cosine similarity of 0.15 and 0.02 with ground truth (whether a patient has Delta or not). Even when extended to 2 dimensions, the clusters do not separate.

**Fig. 20.**
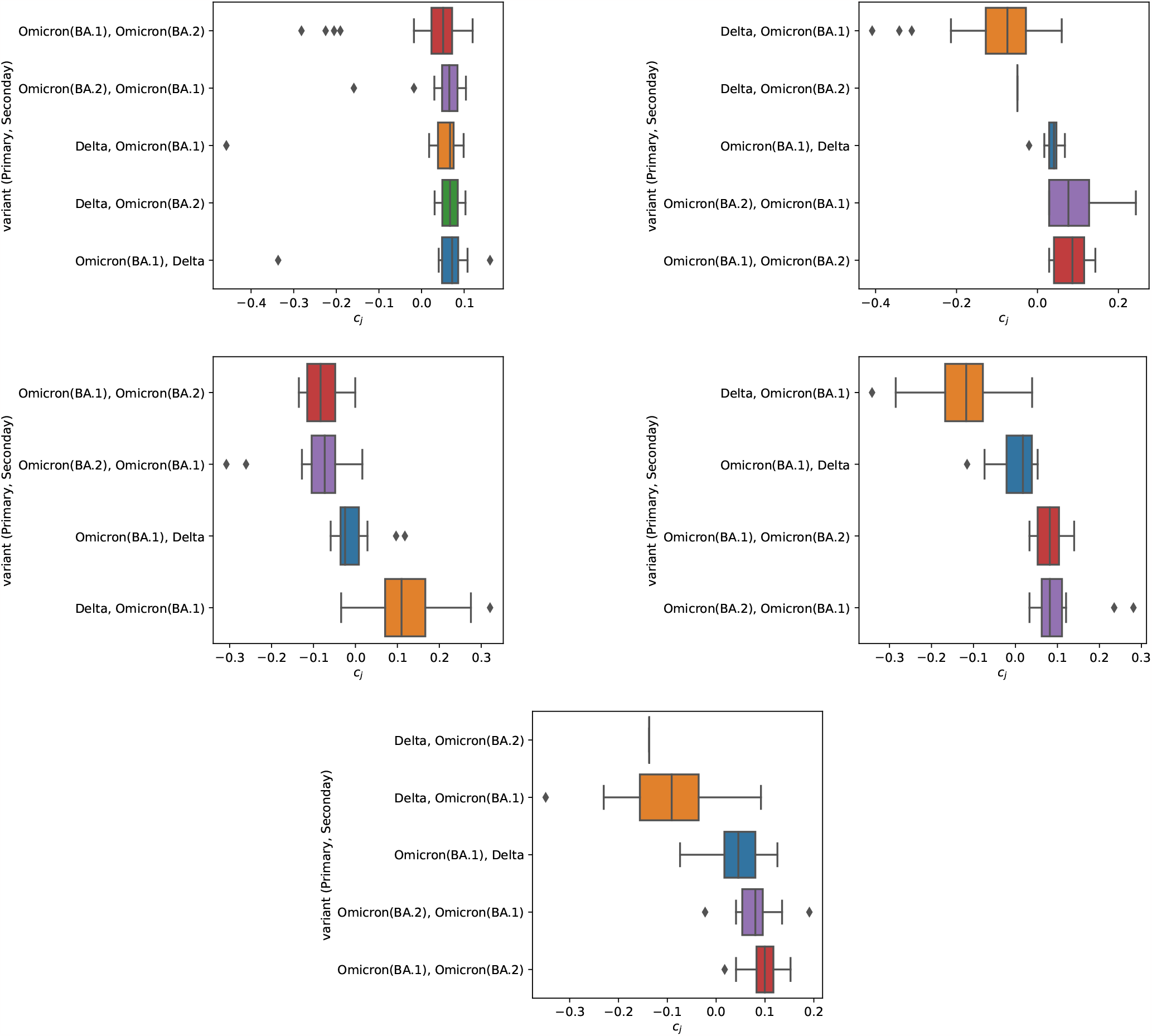
5 tables in `OASIS-opt`’s reduced list of anchors (effect size in the top 10%, *M >* 1000, significant `OASIS-opt` p-value bound). First is 482 *×* 100, 3688 counts, cosine similarity of 0.2. Second is 98 *×* 77, with 2540 counts, cosine similarity of 0.61. Third is 74 *×* 71, 1486 counts, cosine similarity of 0.69. Fourth is 67 *×* 69, 1299 counts, cosine similarity of 0.62. Fifth is 88 *×* 91, 1256 counts, 0.63 cosine similarity.

**Fig. 21.**
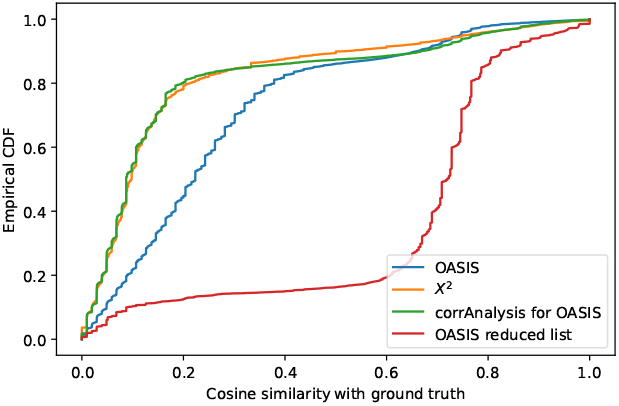
Results on 100,914 tables generated by SPLASH (1) from SARS-CoV-2 data (6). Metric plotted is absolute cosine similarity between binarized sample embedding vector per table (***c*** for OASIS, principle right singular vector from correspondence analysis for *X*^2^) and ground truth vector indicating whether a patient (sample) has Delta or not. Empirical CCDF of absolute cosine similarity between identified vectors per table and ground truth vector is plotted for the tables that each method declares significant after multiple hypothesis correction. Out of the 100,914 tables, OASIS rejects 28,430, and Pearson’s *X*^2^ test rejects 71,543 tables. However, when decomposed, the tables that *X*^2^ rejects do not yield signal that correlates well with the ground truth; examining quantiles of the two distributions, the 0.5 and 0.9 quantiles of these empirical distributions are 0.22 and 0.66 for OASIS, as opposed to 0.10, 0.52 for *X*^2^.

**Fig. 22.**
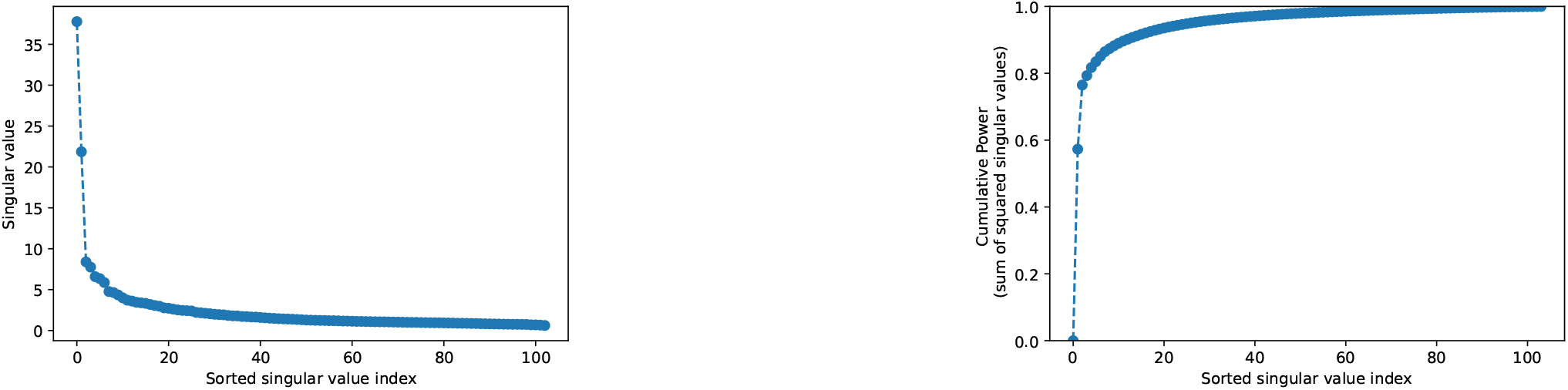
Spectrum of *C* matrix from ***c***-aggregation for SARS-CoV-2 data, showing why 2d embedding is sufficient.

**Fig. 23.**
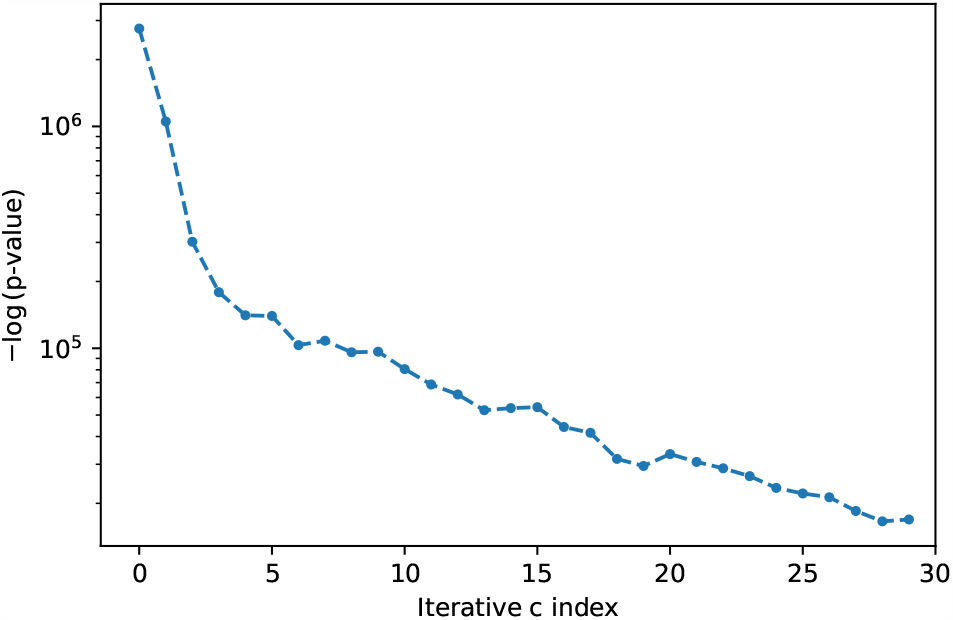
Negative log of p-value bounds for data-aggregation on stacked data matrices for SARS-CoV-2. As can be seen, after the second index, the remaining points follow a slow linear trend (the elbow), indicating that only 2 components should be utilized.

**Fig. 24.**
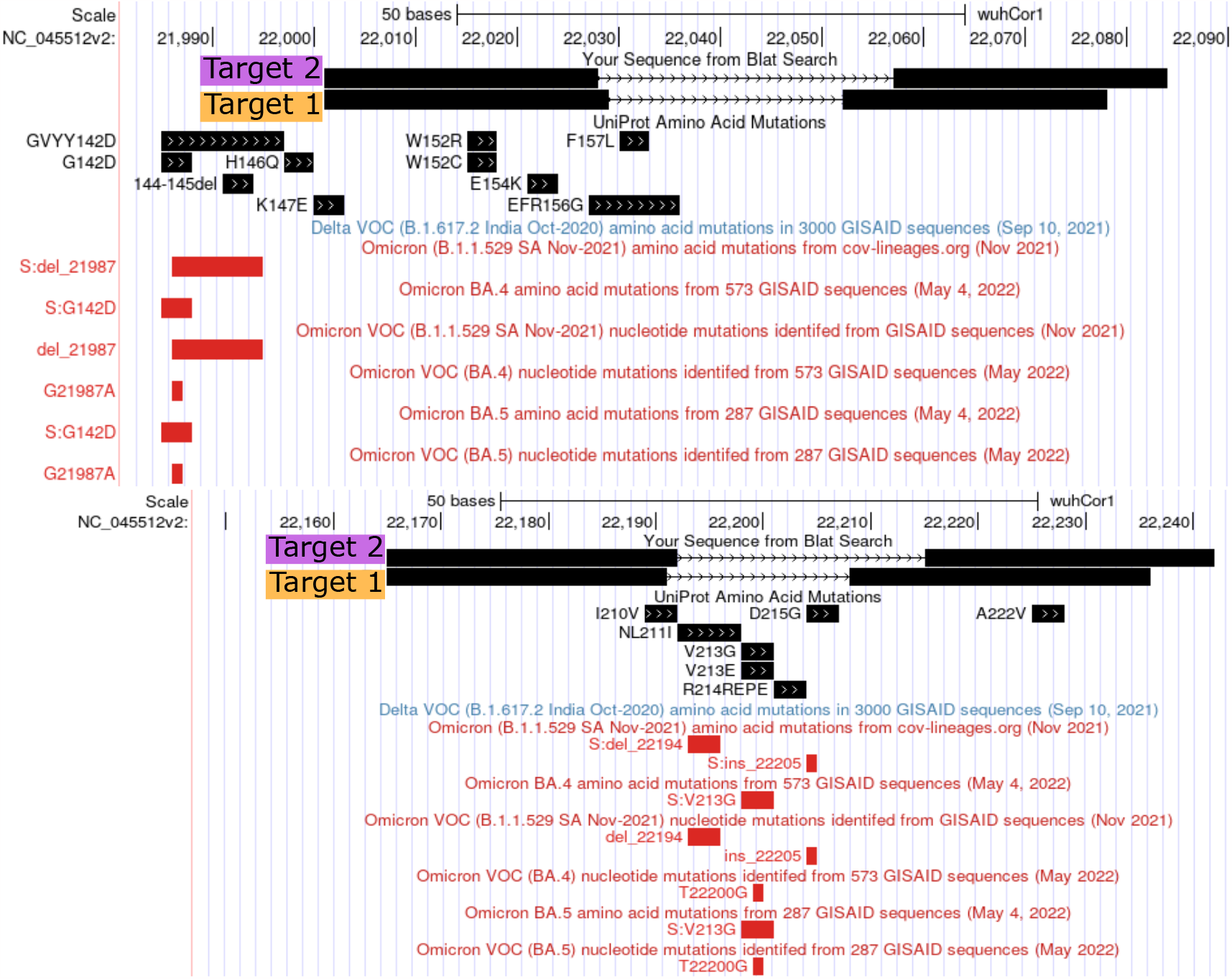
BLAT (34) to SARS-CoV-2 genome via UCSC genome browser (35). For each target, we align the anchor concatenated with the target, where for each of these two anchors, only 2 targets had abundance greater than 5%. Note that these targets do not come immediately after the anchor, but are taken a fixed distance ahead (called the lookahead distance (1)). In both examples, target 1 takes a value *f*_*i*_ = 1, and target 2 takes a value *f*_*i*_ = 0. The reported 93% agreement predicts that *f*_*i*_ = 1 corresponds to Delta and *f*_*i*_ = 0 corresponds to Omicron. Thus, T1 is predicted to be Delta, and T2 to be Omicron. Not shown in the first example, analyzing the raw reads from these samples shows that those from Delta samples follow the genome exactly, whereas those from Omicron samples exhibit a 6 basepair deletion in this gap, leading to the resulting contingency table. This deletion isn’t annotated in the UCSC genome browser, but corresponds to a known Omicron deletion. In the second example, we observe the same behavior (Omicron has a larger gap between anchor and target), this time due to an annotated Variant of Concern (VOC), a deletion.

**Fig. 25.**
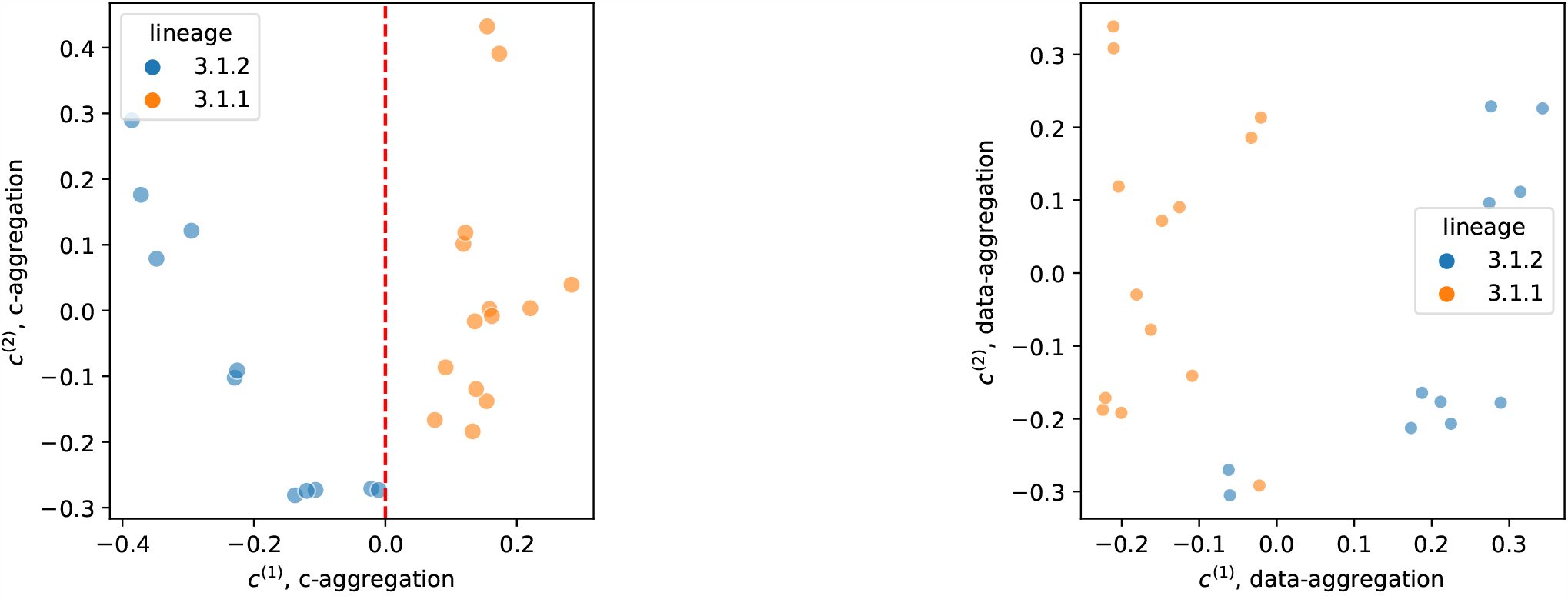
2D plots of Tuberculosis data. Left is ***c***-aggregation, right is data-aggregation.

**Fig. 26.**
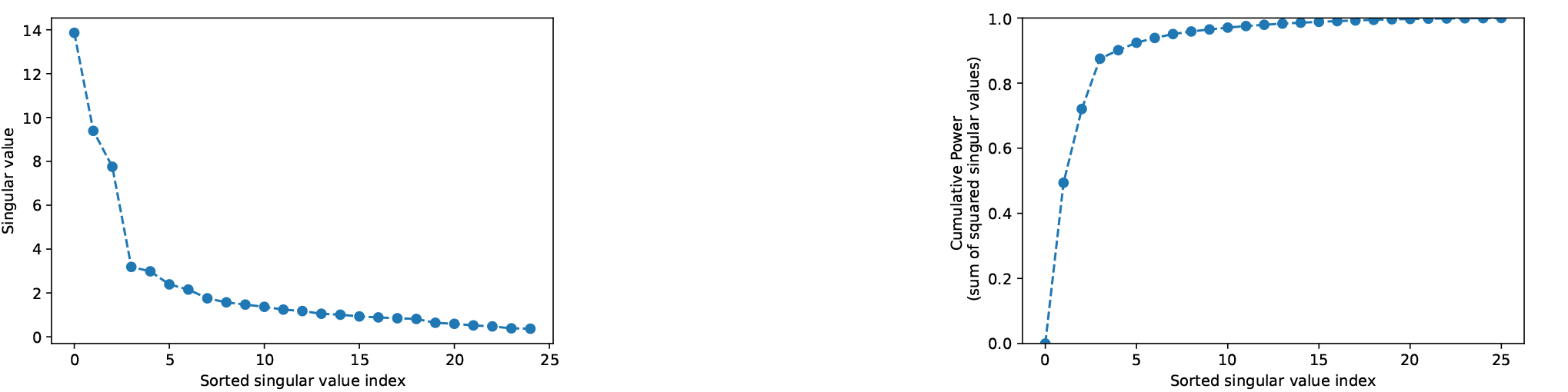
Spectrum of *C* matrix from ***c***-aggregation for Tuberculosis data, showing why 2d embedding is sufficient.

### E. Tuberculosis additional plots

For the analysis of Tuberculosis data, a newer version of the SPLASH pipeline was run, SPLASH2 (26), to generate the contingency tables. Then, the same optimization procedures were run to generate the optimized p-value bounds.

### F. Multiple group

In Figure 27 we observe that OASIS, unaware of the underlying multi-group structure, is able to give a statistically valid stopping criterion for subclustering, correctly identifying the number of underlying clusters as 3 55% of the time when each column has Poisson with mean 20 observations per column, and up to 69% with mean 30 observations per column.

**Fig. 27.**
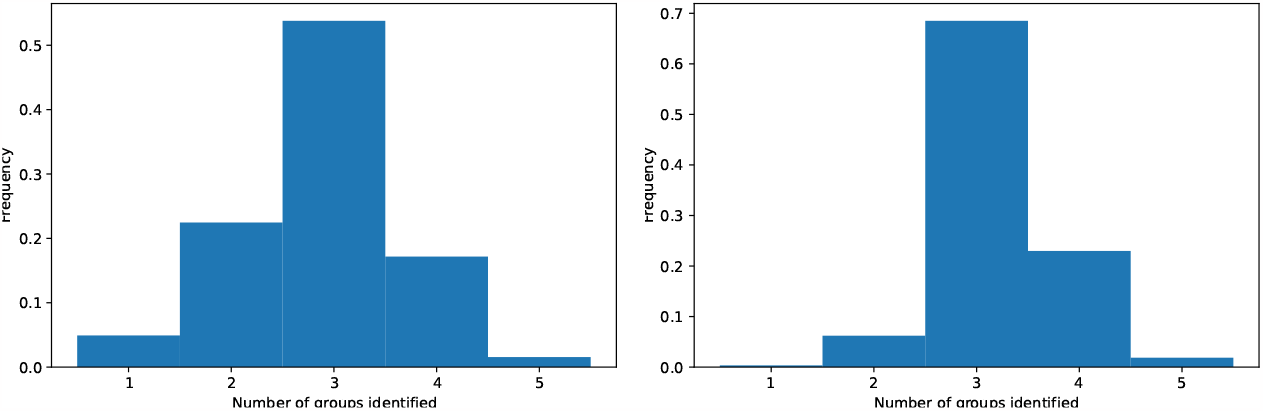
Evaluation of OASIS for group discovery, in a planted setting with 3 groups, 10 rows, and 20 columns. Plotted is the number of identified ***c*** vectors using `OASIS-iter`; finding orthogonal ***c***^(*k*)^ until p-value is no longer significant. Left figure shows 20 observations per column, right shows 30.

### G. Asymptotic normality

As proved in Proposition 2, OASIS’s test statistic is asymptotically normally distributed. However, with asymptotia, one natural question is “how large do counts need to be before the approximation is reasonable”? In Figure 28, we show that even for very small numbers of observations (e.g. 10 counts per column, 8 columns), the normalized test statistic is very close to being normally distributed.

**Fig. 28.**
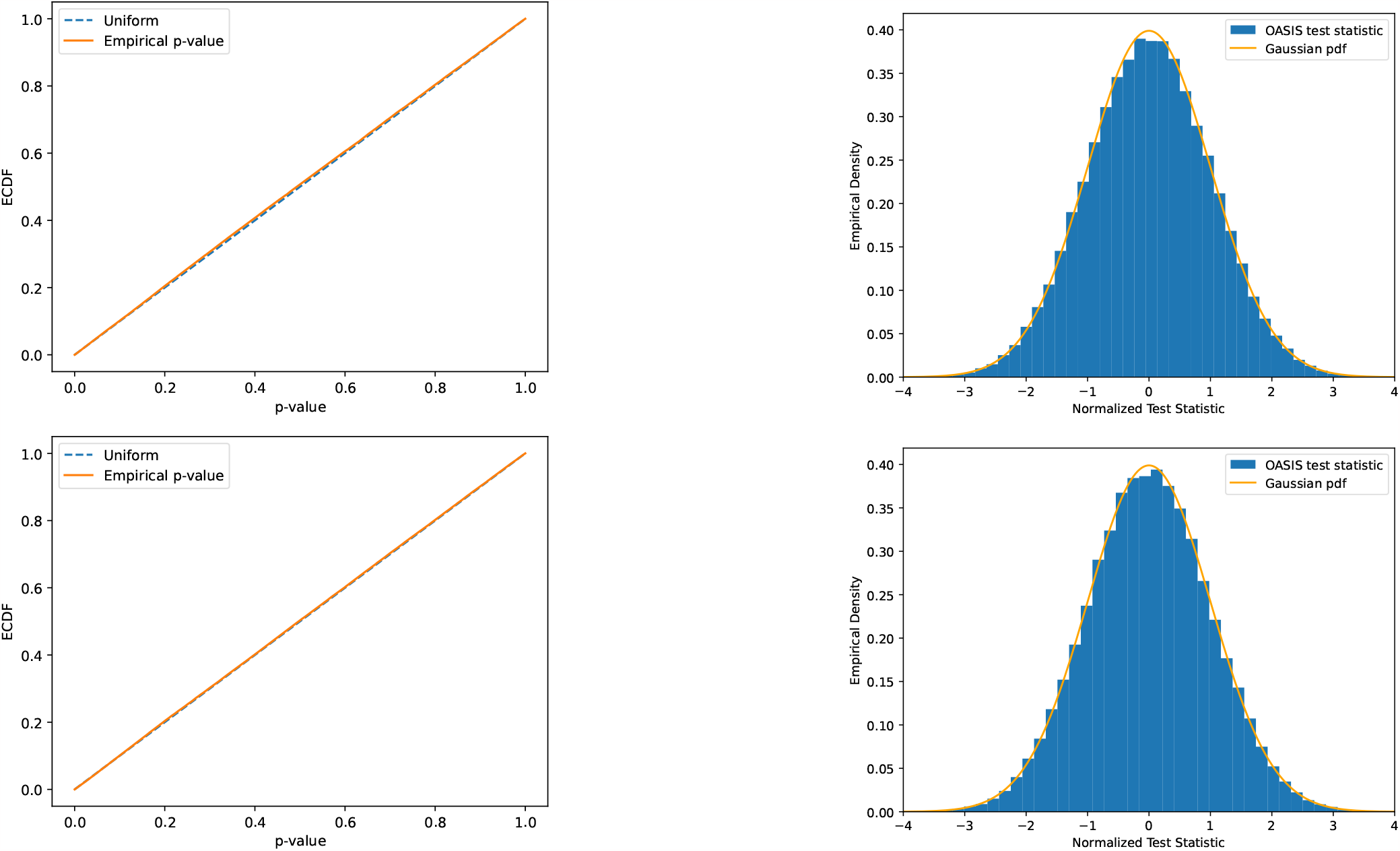
Distribution of OASIS test statistic 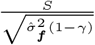. As predicted by theory, this follows a Gaussian distribution, leading to uniformly distributed asymptotic p-values. Important to note is that these tables do not have many counts; tables shown have 5 rows and 8 columns, where in the first there are 10 counts per column in expectation and the second 30. Plotted is simulated data over 100k trials with 5 rows, 8 columns, random target distribution (each entry i.i.d. uniform, normalized). Column counts for first row are independently drawn from a Poisson with mean 10, while second row has mean 30. For each random trial a different random ***p*** was generated, *n*_*j*_ were drawn randomly, *X* was generated from this, then ***c, f*** were independently randomly generated (***f*** coordinate i.i.d. uniform on {0,1}, ***c*** each entry i.i.d. uniform [− 1, 1] then ***c*** normalized. This process was repeated until a table with 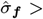 0 and *γ <* 1 was generated to yield a valid run (as otherwise the test statistic is identically 0). KS distance (maximum distance between OASIS’s asymptotic p-value’s ECDF and the uniform distribution of .0079 for *n* = 10 counts per column, .0038 for *n* = 30 counts per column.

As a correlation coefficient, OASIS favorably compares against xicor (33) in some settings, as shown in Figure 29. The example functions used are detailed in (33), where we show that OASIS has additional power against balanced, structured alternatives, such as circular and heteroskedastic.

**Fig. 29.**
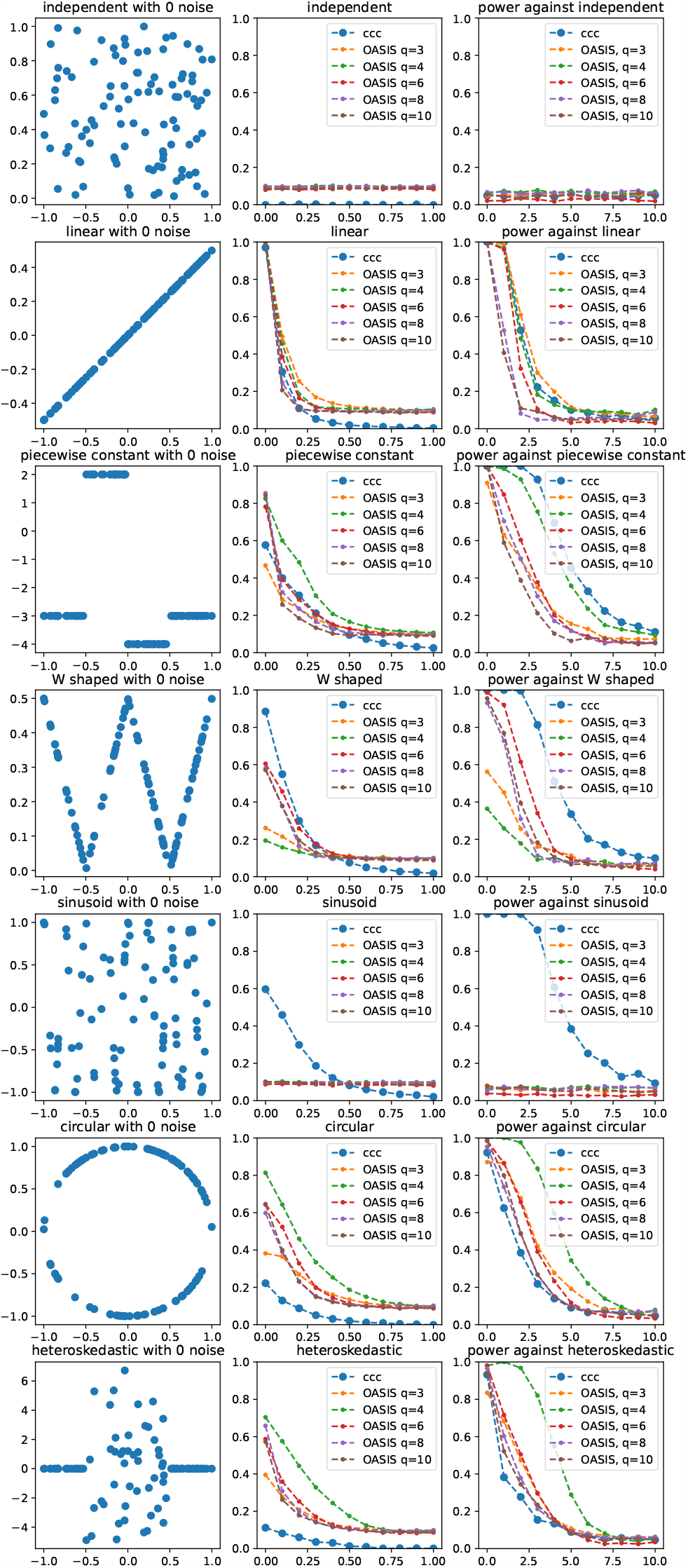
Comparison of `OASIS-opt` and XICOR (33). OASIS asymptotic p-value used, 25% train test split. Each row represents a different setting, where the left column shows the noiseless problem instance *y* = *f* (*x*), the middle column shows the magnitude of the correlation coefficient averaged over 1000 iterations per point, and the right column shows the fraction of time the null was rejected. For OASIS, the middle column shows the effect size, and the right column shows the rejection fraction utilizing the asymptotic p-value. The quantization level *q* is varied across instances; to map the input continuous-valued random variable to a discrete categorical one. This is performed independently for the row and column random variables, and binned into *q* bins of equal counts (1*/q* quantiles).

## 15. Power analysis

Due to the simple closed form expressions for the *X*^2^’s asymptotic p-value, and OASIS’s finite-sample p-value bound, we can perform approximate power calculations across different alternatives. Concretely, we analyze several toy settings where we observe a scaled version of the underlying alternative matrix, and compute OASIS and *X*^2^ p-values, showing that OASIS indeed has substantial power in these settings, and can be more powerful than *X*^2^ in certain regimes. We conjecture that while OASIS may not exhibit the optimal asymptotic rate against all classes of alternatives, e.g. the unique per-sample expression example, it does have power going to 1 as the number of observations goes to infinity across a broad class of alternatives.

In our power computations, we additionally assume that OASIS identifies an optimal pair of ***c, f***. It is reasonable to assume that this occurs in the asymptotic regime, and further since OASIS’s test statistic is bilinear in ***c, f***, its p-value bound is continuous in these parameters, and so OASIS only needs to identify a pair of near optimal ***c, f*** to yield a similar p-value bound.

The *X*^2^ p-value for *k* degrees of freedom is asymptotically distributed as 𝒩(*k*, 2*k*). Since the *X*^2^ test statistic is 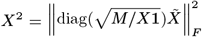, plugging in = (− 1) × (− 1), and examining large values of the test-statistic,

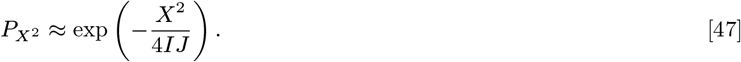

### A. Unique per-sample expression

In this setting, the alternative matrix is *c* × *c*, where the *j*-th sample expresses target (row) *j* with probability 1 −*α*, and with probability *α* is uniform over the rest of the *c* −1 rows. The number of observations per column are assumed to be the same. Formally, the alternative matrix *A* is as below:

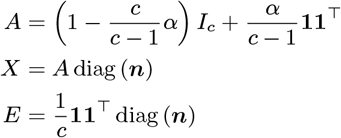

In this setting, Pearson’s *X*^2^ statistic is

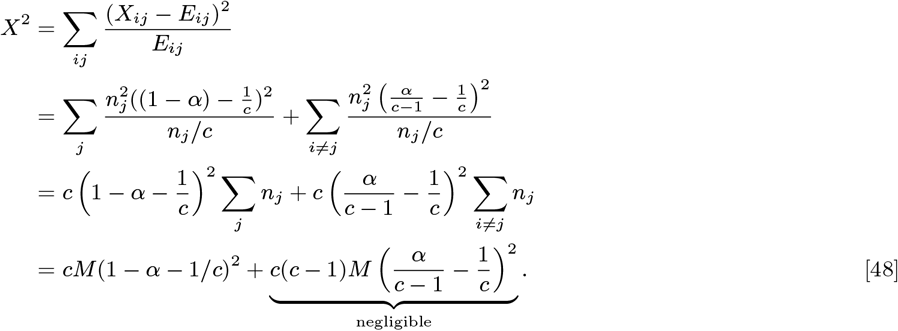

Thus, for fixed *c*, as *M*→ ∞, the *X*^2^ p-value can be bounded as

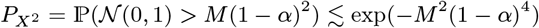

Analyzing OASIS, we assume that it identifies an optimal pair of ***f***, ***c***, where since *n*_*j*_ are balanced

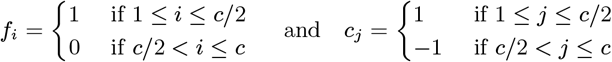

constitute an optimal pair. Since *n*_*j*_ = *M/c* for all *j*, 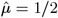, and so

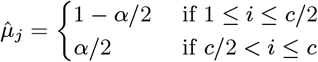

This yields

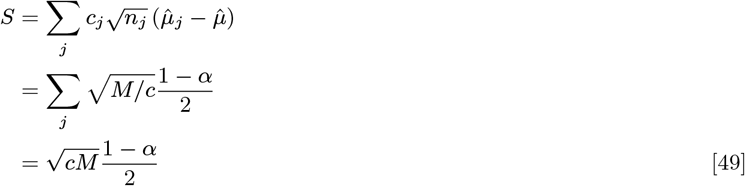

and a corresponding p-value bound of

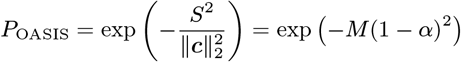

#### A.1. Summary

In this toy-setting of unique per-sample expression, *X*^2^ will yield a p-value of

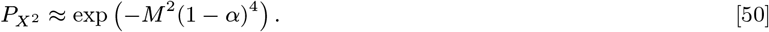

Comparatively, assuming OASIS identifies a pair of optimal ***c, f***, it will obtain a p-value bound of

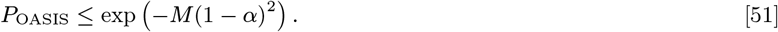

While this is worse for constant *α* and *M* tending to infinity, OASIS still has power going to 1 as *M* increases.

### B. Two-group alternative

Here we assume for simplicity that all *n*_*j*_ are equal, and that the clusters are of equal sizes. This can be relaxed by using a class-weighted mean estimate, which computes the mean *f* value over samples with *c*_*j*_ *>* 0, and averages this with the mean *f* value over samples with *c*_*j*_ *<* 0.

We describe the results in the context of differentially regulated alternative splicing, but the analysis holds for general 2 group alternatives with the below structure. For samples *j* = 1, …, *J/*2, the target (row) distribution is ***p***^(1)^, and for the second half of columns ***p***^(2)^. In this setting we have:

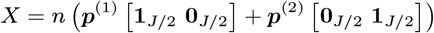

and thus

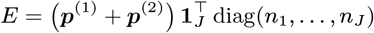

The Pearson *X*^2^ statistic is then, using 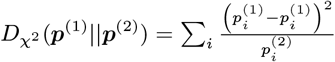 :

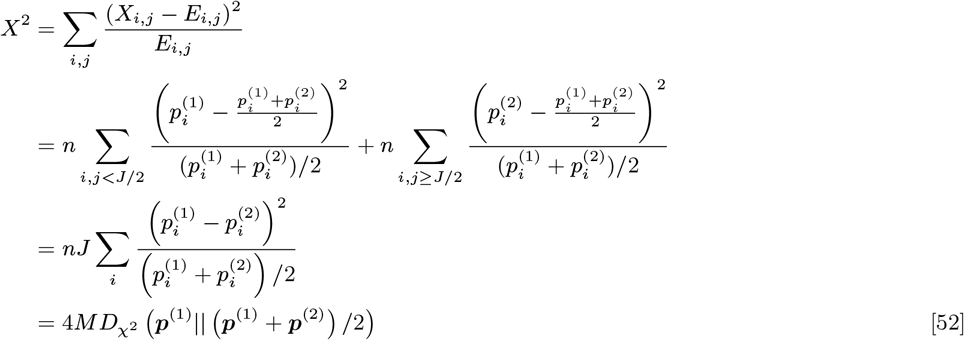

For disjoint ***p***^(1)^, ***p***^(2)^, the *χ*^2^ distance is equal to 1. From the (*I* −1) × (*J*− 1) degrees of freedom, this yield an approximate p-value of

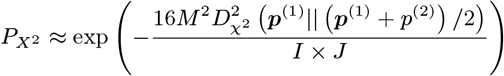

For OASIS, a p air of optimal ***c*** and ***f*** include where ***c*** is +1 for the first *J/*2 components, and −1 for the latter half, and ***f*** is sign ***(p***^(1)^ − ***p***^(2)^). Then, the OASIS test statistic is

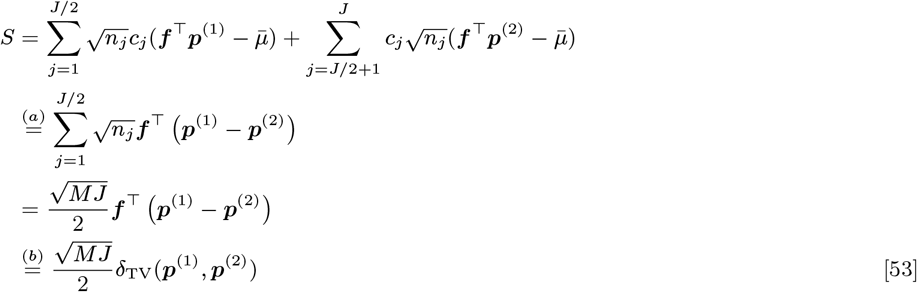

where in (a) we used the fact that all *n*_*j*_ were the same and that the clusters were equally balanced. If we use the class-weighted mean estimate sketched earlier then we can overcome the equally balanced classes problem. The *n*_*j*_ all being the same can be overcome by still assuming that they all go to infinity, but applying Jensen’s inequality to lower bound the test statistic as above. In (b) we used the optimal ***f*** = sign(***p***^(1)^ − ***p***^(2)^).

#### B.1. Summary

The *X*^2^ p-value can be bounded as

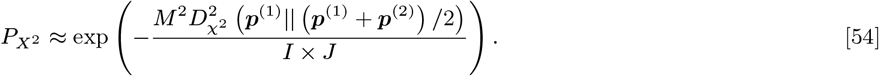

OASIS has significant power against alternative splicing-type alternatives. Assuming that OASIS identifies the optimal ***c, f***, which will be the case asymptotically, it will yield a p-value bound of

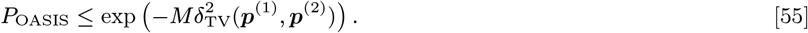

Assuming the total variation distance and *χ*^2^ distance between the distributions are constant, OASIS has more power than *X*^2^ when *I* × *J* ≥ *M* ; this is often the case we are in, where there are many rows with very low counts.

### C. Time series

Consider the simple case where our data represents cells at progressing times during their cell-cycle. At time *t* = 0, they have target distribution ***p***^(1)^, and at time *t* = 1 they have target distribution ***p***^(2)^, smoothly interpolating between them. Assuming for simplicity that the samples are evenly spaced along this time course (i.e. cell *j* is at time *t*_*j*_ = (*j* − 1)*/*(*J* + 1)), and that all are observed equally (same *n*_*j*_), the optimal *c*_*j*_ = 2*t*_*j*_ − 1, so *t* = 0 ⇒ *c*_*j*_ = −1 and *t* = 1 ⇒ *c*_*j*_ = 1.

Consider the simplest time series setting, where at time *t* = 0 cells have target distribution ***p***, at time *t* = *T* distribution ***p***^(2)^, and vary smoothly in between, i.e. at time *t* the mixture distribution ***p***^(1)^ + *t*(***p***^(2)^ − ***p***^(1)^) = (1− *t*)***p***^(1)^ + *t****p***^(2)^. For *T* + 1 evenly spaced time intervals, define the vector ***t*** = [0, 1*/T*, …, (*T* − 1)*/T*, 1]. We observe that, for *n* observations per column, our expected data matrix *X*, and null mean matrix *E*, would be

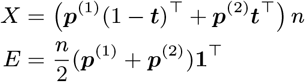

The p-value obtained from Pearson’s *X*^2^ test is:

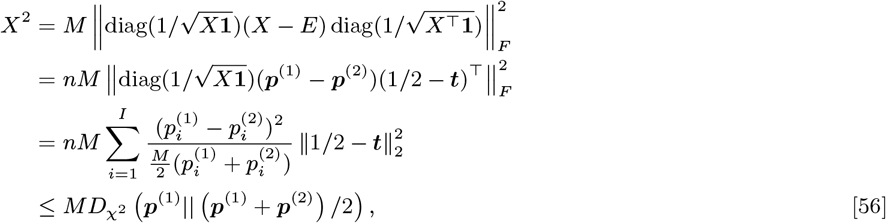

since Frobenius norm of rank one outer product simplifies. 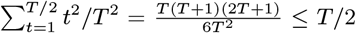 for large enough *T*, and so 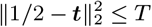.

This leads to an approximate p-value of

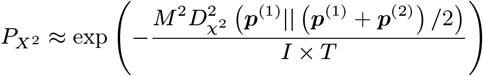

To compute OASIS’s power, we first compute *S*_*j*_

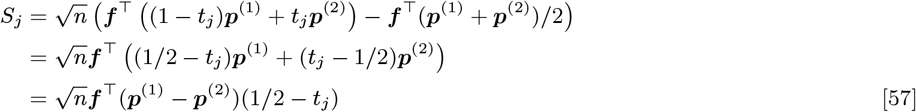

We know that *c*_*j*_ ∝ *S*_*j*_ are optimal as

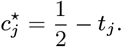

The maximization of ***f*** is simple now, as the same term shows up in each column:

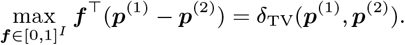

Computing *S* for optimal ***c, f*** :

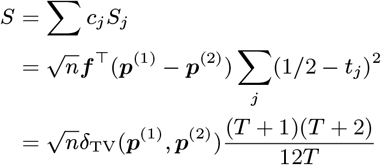

This gives OASIS a p-value bound of (for large enough *T*)

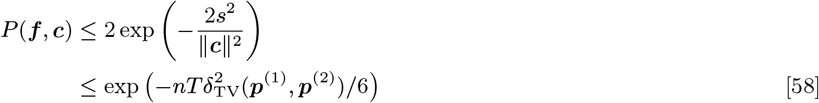

#### C.1. Summary

*X*^2^ provides an approximate asymptotic p-value of

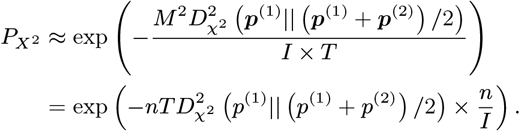

OASIS yields a p-value bound of

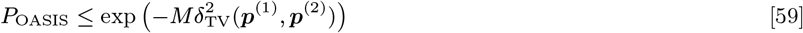

This setting displays the same behavior as in alternative splicing, where OASIS can yield improved performance when 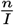 is small.

### D. One deviant sample

Consider the setting where one sample is maximally deviating, i.e. the observed matrix is:

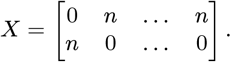

Then, the expected matrix is

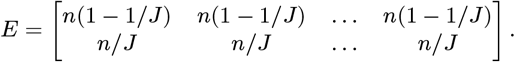

Expressing the summation as grouped by *X*_00_, *X*_10_, *X*_0,1:_, *X*_1,1:_

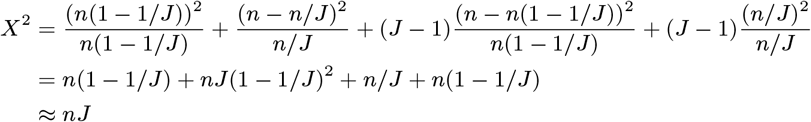

Thus, the Pearson *X*^2^ test will yield an approximate p-value of

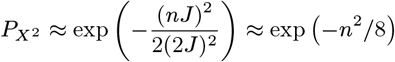

Comparatively, OASIS will choose = [1 0]^⊤^, which yields 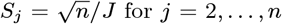 for = 2 and 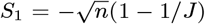. This yields an optimizing ***c*** = [−(1 − 1*/J*) 1*/J* … 1*/J*], where ∥***c***∥^2^ = 1 − 1*/J* here. This yields a test-statistic value 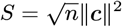

Thus,

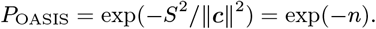

#### D.1. Summary

*X*^2^ provides a p-value of

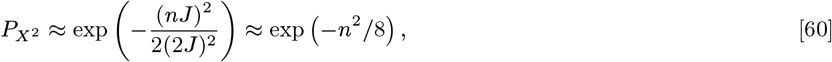

in comparison to `OASIS-opt`’s p-value bound of

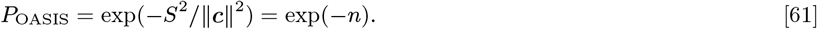

This shows how *X*^2^ can be significantly overpowered against alternatives where only one column deviates.

### E. Conclusions from power analysis

As can be seen, there are several regimes where the Pearson *X*^2^ test has less power than OASIS, primarily when the table is sparse, i.e.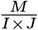 is small. This highlights the difficulty with the *X*^2^ test; its p-value depends heavily on the number of rows present, even if many of them are inconsequentia

